# Integrative analysis of epigenetics data identifies gene-specific regulatory elements

**DOI:** 10.1101/585125

**Authors:** Florian Schmidt, Alexander Marx, Marie Hebel, Martin Wegner, Nina Baumgarten, Manuel Kaulich, Jonathan Göke, Jilles Vreeken, Marcel H. Schulz

**Affiliations:** Cluster of Excellence for Multimodal Computing and Interaction, Saarland Informatics Campus, 66123, Saarbrücken, Germany; Max Planck Institute for Informatics, Saarland Informatics Campus, 66123, Saarbrücken, Germany; Graduate School of Computer Science, Saarland Informatics Campus, 66123, Saarbrücken, Germany; International Max Planck Research School for Computer Science, Saarland Informatics Campus, 66123, Saarbrücken, Germany; Institute of Biochemistry II, Goethe University Frankfurt - Medical Faculty, University Hospital, Frankfurt am Main, Germany; Institute for Cardiovascular Regeneration, Goethe University, 60590 Frankfurt am Main, Germany; German Center for Cardiovascular Regeneration, Partner site Rhein-Main, 60590 Frankfurt am Main, Germany; Frankfurt Cancer Institute, Frankfurt am Main, Germany; Computational Genomics and Transcriptomics, Genome Institute of Singapore, 60 Biopolis Street, 138672 Singapore; Helmholtz Center for Information Security, Saarland Informatics Campus, 66123 Saarbrücken, Germany

**Author notes:** Corresponding authors: Florian Schmidt, Marcel H. Schulz.

## Abstract

Understanding the complexity of transcriptional regulation is a major goal of computational biology. Because experimental linkage of regulatory sites to genes is challenging, computational methods considering epigenomics data have been proposed to create tissue-specific regulatory maps. However, we showed that these approaches are not well suited to account for the variations of the regulatory landscape between cell-types. To overcome these drawbacks, we developed a new method called STITCHIT, that identifies and links putative regulatory sites to genes. Within STITCHIT, we consider the chromatin accessibility signal of all samples jointly to identify regions exhibiting a signal variation related to the expression of a distinct gene. STITCHIT outperforms previous approaches in various validation experiments and was used with a genome-wide CRISPR-Cas9 screen to prioritize novel doxorubicin-resistance genes and their associated non-coding regulatory regions. We believe that our work paves the way for a more refined understanding of transcriptional regulation at the gene-level.

## Background

Elucidating the diversity of transcriptional regulation is a prevalent problem in computational biology. While there is a plethora of mechanisms involved in regulating transcription [9], especially the binding of Transcription Factors (TFs) to regulatory elements (REMs) such as *Promoters*, *Enhancers*, and *Repressors* has been shown to be essential for orchestrating cellular development and identity [54, 57]. Importantly, enhancers have been closely linked to several diseases [18] and recent research suggests that enhancers might be therapeutic targets [57, 45].

n order to describe how REMs might influence their target genes in a systematic way, two models have been proposed: the scanning model and the looping model [57, 4]. According to the scanning model, a REM is usually affecting a gene that is located in close genomic distance, whereas in the looping model, REMs can influence a gene that is located several kilobases (kb) away from the actual regulatory site via chromatin looping. Because biological evidence has been found for both models, it is likely that both do occur *in-vivo* [60, 29].

To elucidate regulatory function, two main problems need to be solved: Firstly, REMs, need to be identified genome wide and secondly, they need to be assigned to their target genes. The first problem, identifying REMs genome wide, has been addressed by international projects, e.g. ENCODE and Roadmap. There, REMs were identified using DNase1-Hypersensitive Sites (DHS), e.g. sites of accessible chromatin [48, 53], via distinct patterns of Histone Modifications (HMs), i.e. the co-occurrence of H3K27ac, H3K4me1 while H3K4me3 is absent [23], or TF-ChIP-seq experiments of TFs such as EP300 [55]. Typically, such data sets are analysed with peak calling algorithms. Although, there is a plethora of peak callers available, designed for ChIP-seq [52] and chromatin accessibility data [28], peak callers still have several limitations. For instance, the selection of the cut-off to determine peaks over background is not trivial, and also cell cycle stage [30] or cell numbers [15] can prevent us from accurately detecting all truly enriched regions. Furthermore, it is often not clear what level of enrichment is needed such that a region can be seen as biologically active [7]. Besides, as illustrated in Supplementary Fig. S1, integrating peak calls across several diverse samples is not straightforward [31]. However, an integrated set of peaks is required if machine learning approaches should be utilized to associate a defined set of candidate REMs to potential target genes across many samples. Note that automated integration of replicates, as offered e.g. in the peak caller JAMM [24], is not designed for such an application. It is rather meant to provide stable, reproducible peak calls across replicates of the same cell-type or tissue.

In addition to the efforts taken by ENCODE and Roadmap, putative enhancers were identified in the Fantom5 consortium via the identification of distinct bidirectional expression patterns in CAGE (Cap Analysis of Gene-Expression) data [1].

Overall, many different ways have been proposed to identify putative REMs using distinct chromatin signatures. Nevertheless, the problem of linking those regions to the genes they regulate is still not straightforward to solve. In literature, especially in instances were only few replicates are available, putative REMs are often linked to their nearest gene according to genomic distance [16], or aggregated using window based approaches [42, 33]. However, several studies suggest that especially enhancers and repressors do not regulate their nearest gene but may influence more distant genes [1, 21, 41]. On top of that, REMs are highly tissue-specific [36], underlying that a purely distance based detection of REMs is error prone.

*Yao et al.* [57] describe two approaches attempting to overcome these limitations: (1) methods based on physical interaction, i.e. capture Hi-C [25], or Chromatin Interaction Analysis by Paired-End Tag sequencing (ChIA-PET) [13] and (2) methods based on associating gene-expression to the activity of REMs, e.g. using DNase1-seq [53, 21], or HM abundance [10].

While methods based on physical interaction are laborious, time consuming and experimentally challenging, e.g. in terms of providing a sufficient resolution of long-range contacts [38], association based methods are predestined to use the plethora of available epigenetics data to link REMs to their target genes.

Using machine learning, *Cao et al.* propose to integrate predicted REMs into cell-type specific interaction networks [6], similar to *Hait et al.*, who also provide regulatory-maps derived from statistical associations between the activity of REMs and target gene-expression [21]. *Shooshtari et al.* combined chromatin accessibility data with Genome-Wide Association study Studies (GWAS) to better pinpoint regulatory events in autoimmune and inflammatory diseases [47]. In the Fantom consortium, putative REMs have been linked to their target genes by associating enhancer activity to gene-expression [1]. *Gonzales et al* use a nearest gene linkage of DHS sites in an iterative manner within gene-expression models to link REMsto their target genes [16].

## Results

### A novel method for the gene-specific identification of regulatory sites

We present STITCHIT a novel segmentation based method to identify gene-specific REMs. Unlike other approaches [6, 21], STITCHIT is a peak-calling free approach interpreting the epigenetic signal in relation to the expression of a distinct gene. Basically, STITCHIT solves a classification problem by segmenting a large genomic area around the target gene. The resulting segmentation highlights regions exhibiting epigenetic signal variance, which is linked to the expression of the analyzed gene. As illustrated in Fig. 1, STITCHIT is superior to peak-based approaches as it is able to pinpoint REMs with high resolution and accuracy. Here, we apply STITCHIT to a large collection of paired, uniformly reprocessed DNase1-seq and RNA-seq samples from Blueprint, ENCODE, and Roadmap to determine gene-specific REMs. These datasets are very different, e.g. the Blueprint dataset is rather homogeneous representing a wide spectrum of the haematopoietic lineage, the ENCODE dataset is composed of a few diverse samples, and the Roadmap dataset is a large, highly diverse, heterogeneous dataset. Thus, these three datasets are ideal to test the capabilities of STITCHIT, which we did in various validation and application scenarios.

**Figure 1:**
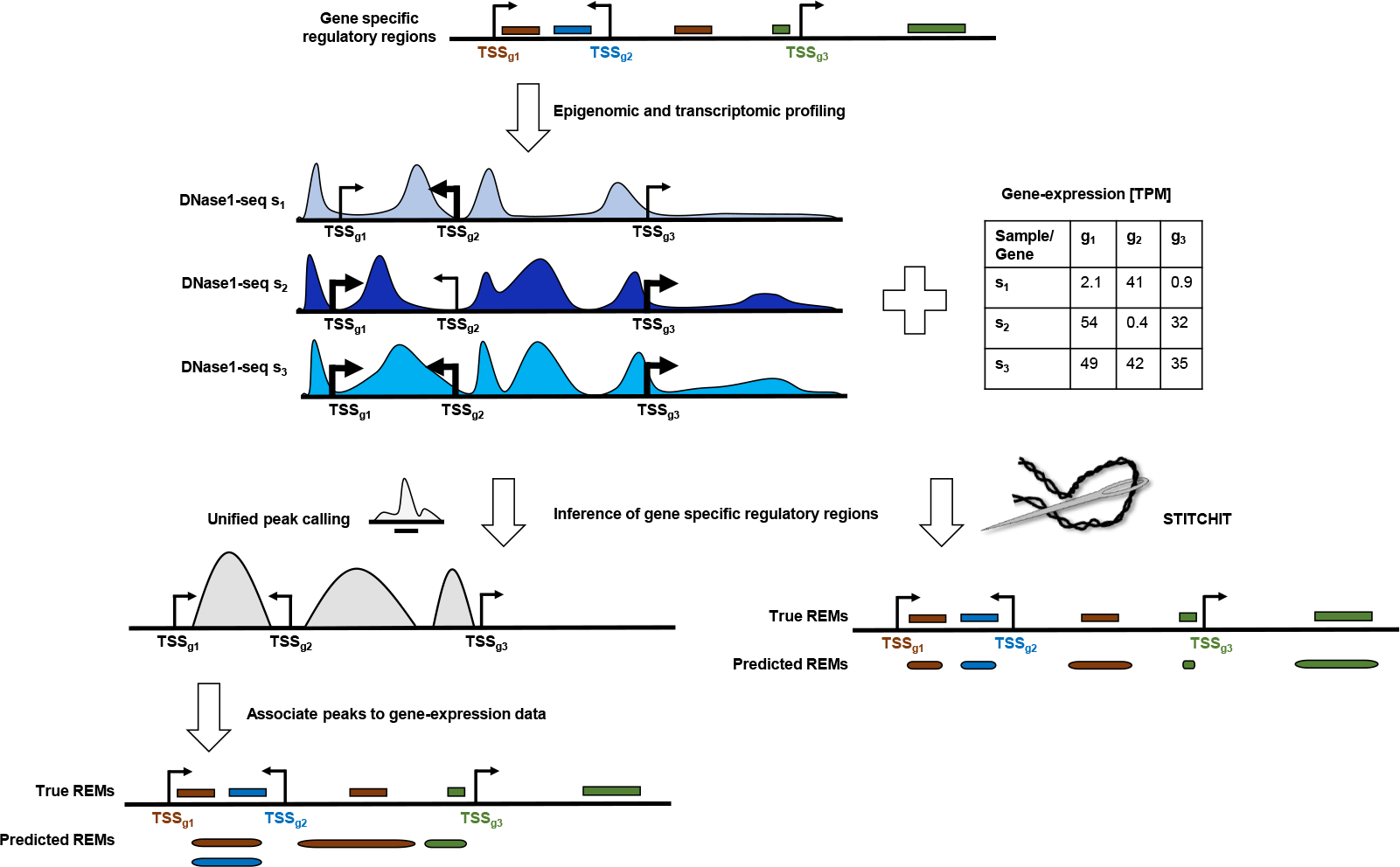
In this illustration, we consider three genes (*g*_1_, *g*_2_, *g*_3_), with distinct, colour coded regulatory regions indicated at the top of the Figure. These genes are profiled in three samples using DNase1-seq and RNA-seq, revealing distinct patterns in chromatin accessibility and in gene-expression, respectively. Two approaches are used to derive regulatory elements (REMs) for these genes. One the left, REMs are derived based on correlation tests between called peaks and the expression of the target genes. As shown, the peak based linkage is inaccurate. For instance, the neighbouring REMS for *g*_1_ and *g*_2_ are not distinguished accurately due to insufficient resolution during peak calling. Similarly, a distal REM for *g*_1_ is predicted to be too large and an intragenic REM of *g*_3_ is missed, due to loss of sensitivity by the peak caller. STITCHIT instead, uses the OC and gene-expression data in a unified statistical framework to define gene-specific REMs with greater accuracy, as shown on the right hand site.

STITCHIT has two main parameters that influence performance and runtime: the *segment-size* and the *resolution*. We have tested several values for both parameters and have set the segment-size to 5000 and the resolution to 10 (Supplementary Fig.s S5) as these parameters yield a good trade-off between performance and runtime.

An additional parameter that is to be specified is the size of the considered genomic region up- and downstream of a gene. This parameter influences whether distal associations can be discovered and influences the runtime of the tool. We have conducted runtime experiments (Supplementary Fig. S6) and found that even with a window size of 0.5*MB* (excluding the size of the genes) REMs can be learned in about 10 minutes per gene. As regulatory interactions typically arise within topological associated domains [8], this is also a feasible value in practice, especially for analyses focusing only on a few distinct genes.

### Regulatory regions derived from STITCHIT perform well in gene-expression prediction

Prior to a biological evaluation of the suggested REMs, we investigate their general characteristics. Here, computed REMs using STITCHIT, the UNIFIEDPEAKS approach, the GENEHANCER database, and using the aggregation of individual DHS sites for the datasets originating from Blueprint, ENCODE and Roadmap are investigated more closely.

As illustrated in Fig. 2a both STITCHIT and UNIFIEDPEAKS identify more candidate regions per gene than GENEHANCER. Simultaneously, the regions retrieved by STITCHIT and UNIFIEDPEAKS are shorter than those extracted from GENEHANCER (Fig. 2b). The same observation is made using Pearson correlation as a measure to filter candidate regions (Supplementary Fig. S7). This suggests that although STITCHIT predicts more individual segments, the total genomic space covered by those must not be larger than that of UNIFIEDPEAKS regions. As shown in Supplementary Table S7, the UNIFIEDPEAKS regions indeed cover a larger fraction of the genome than STITCHIT and GENEHANCER regions.

**Figure 2:**
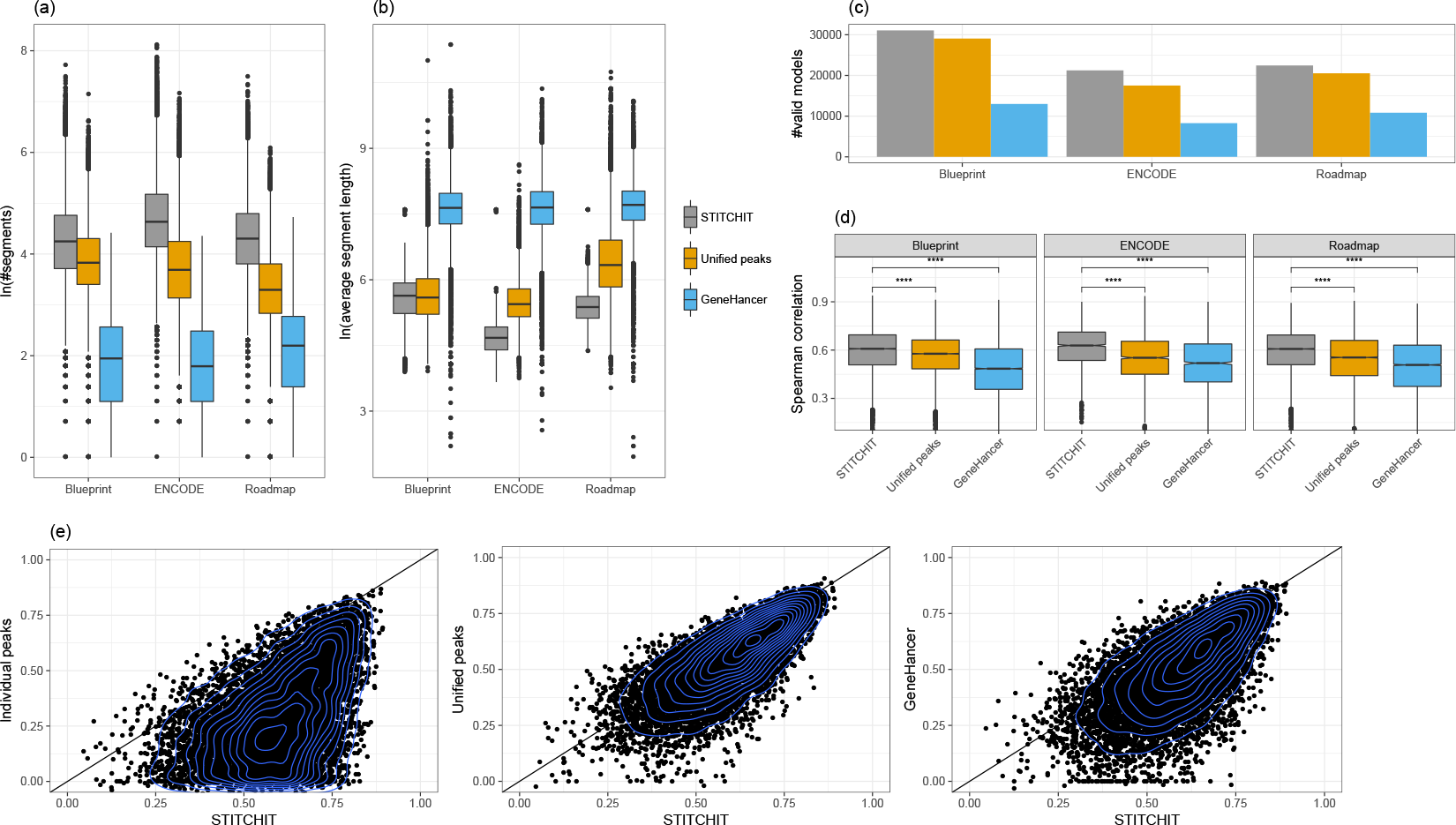
(a) The natural logarithm of the number of segments selected by STITCHIT, UNIFIEDPEAKS, and GENEHANCER is shown for each dataset respectively, whereas in (b), the average length of the selected segments is depicted. The number of learned models is shown in (c), separately per consortia and method. (d) Boxplots showing Spearman correlation between predicted and measured gene-expression using linear regression with elastic net penalty considering all regions identified by STITCHIT, the UNIFIEDPEAKS approach, GENEHANCER, and individual peak aggregation respectively for Blueprint, ENCODE, and Roadmap data. Within STITCHIT, UNIFIEDPEAKS, and GENEHANCER Spearman correlation was used for the initial filtering of candidate regions. Within each consortia, the same set of genes is displayed to allow comparability (Blueprint: 11140, ENCODE: 2057, Roadmap: 9102). As indicated by a two-sided t-test, STITCHIT regions achieve the best model performance (****: *p* ≤ 0.0001). The estimated values for the variances are: 0.018, 0.017,0.029, 0.038 (Blueprint), 0.018, 0.024, 0.032, 0.041 (Roadmap), 0.016, 0.021,0.026, 0.048 (ENCODE), for STITCHIT, UNIFIEDPEAKS, GENEHANCER and the peak aggregation respectively. (e) Scatter plots comparing the performance of STITCHIT (x-axis) against the individual peak aggregation, UNIFIEDPEAKS and GENEHANCER regions (y-axis) on Roadmap data. Each plots shows 9102 genes. A single dot represents the performance of gene-expression models for a distinct gene.

Figure 2c depicts the number of genes for which a model could be learned per consortia and linkage method. STITCHIT and UNIFIEDPEAKS segments lead to more statistically significant models than GENEHANCER segments, while STITCHIT has a slight quantitative advantages compared to UNIFIEDPEAKS.

In Fig. 2d, the Spearman correlation of elastic net models predicting gene-expression from the DNase1-seq signal within the identified REMs is depicted (c.f. Supplementary Fig. S8a for other measures). To allow for comparability, we only show model performance for genes that are covered by each tested method. In Supplementary Fig. S8a, we also show the performance for the window based peak aggregation, labeled as *Individual peaks*. There, we show for each gene only the best performing model based on either the 5kb, 50kb, or the *geneBody* window. Across all datasets, we observe that models based on STITCHIT regions achieve a significantly better correlation (*p* ≤ 0.0001) than models based on the other approaches. This is independent from the correlation measure used for the initial filtering of REMs within STITCHIT, UNIFIEDPEAKS, and GENEHANCER. In a gene-to-gene comparison, as shown in Fig. 2e for Roadmap data (c.f. Supplementary Fig. S8b-d for all datasets), STITCHIT shows a favorable performance.

In case of Blueprint data, we observe that the absolute performance difference between STITCHIT and UNIFIEDPEAKS is less pronounced than for the other two datasets. This is reflected as well by the number of selected regions and their length. The median length of selected regions is more similar for Blueprint data between STITCHIT and UNIFIEDPEAKS than for ENCODE and Roadmap data (Fig. 2b). At the same time, their average length is almost identical, in contrast to the segments identified in the other two datasets (Fig. 2a). In terms of total genomic coverage, the UNIFIEDPEAKS approach covers about 1.65 times of the space covered by STITCHIT on Roadmap data, whereas the difference on Blueprint data is much less (1.02) (Supplementary Table S7).

Further, our results clearly indicate that the supervised generation of REMs outperforms the unsupervised selection considerably, as different window sizes used with the unsupervised approach can not generalize well across different genes (Supplementary Fig. S9).

As illustrated in Supplementary Fig. S8a, using Spearman correlation for the internal filtering leads to a better model performance and was thus used for all remaining experiments in the manuscript.

### STITCHIT learns more putative regulatory regions than related approaches

To better understand the nature of STITCHIT REMs, we have conducted several statistical analyses. While not only the number and the length of REMs is different between datasets (Fig. 2a,b, Supplementary Fig. S11a), we also find that the overlap in terms of genes for which a model could be learned, is generally less than 50% between two datasets (Supplementary Fig. S10), independent from the method used for the computation. Specifically for STITCHIT, only 12.7% (4477) of all gene-specific models are shared between Blueprint, Roadmap, and ENCODE. Roughly 23% (8214) of all genes could be exclusively modeled using Blueprint data, about 2% (6917) with Roadmap, and 6% (2175) with ENCODE.

To gain a better understanding of the characteristics of the suggested regions learned with STITCHIT, we overlapped them with the ERB. Interestingly, although the absolute numbers differ, the relative trend among the different datasets is similar: most STITCHIT regions do not overlap an annotated region (Fig. 3b-d), thus they are labelled as *Unknown*. Only in about a quarter of all cases an overlap is found with a state annotated as Promoter, Promoter flanking region, TF binding site, Enhancer, or Open chromatin. The questions arises whether the remaining regions are simply noise or whether they reflect REMs that have not been annotated so far. To investigate whether these unknown REMs are performing regulatory functions, we assessed the H3K27ac signal in four randomly chosen Blueprint samples within the top 10, 000 STITCHIT REMs (c.f. Methods), within a randomly shuffled set of the same size, and within the top 10, 000 regions per class contained in the ERB. As indicated in Fig. 3e the strongest H3K27ac signal occurs within *Promoter* and *Promoter Flanking Regions*. Importantly, the signal of the randomly distributed regions is the lowest. The signal of the *Unknown* regions is similar to that of *TF binding sites* and *Open Chromatin* suggesting that these regions do have a regulatory effect.

**Figure 3:**
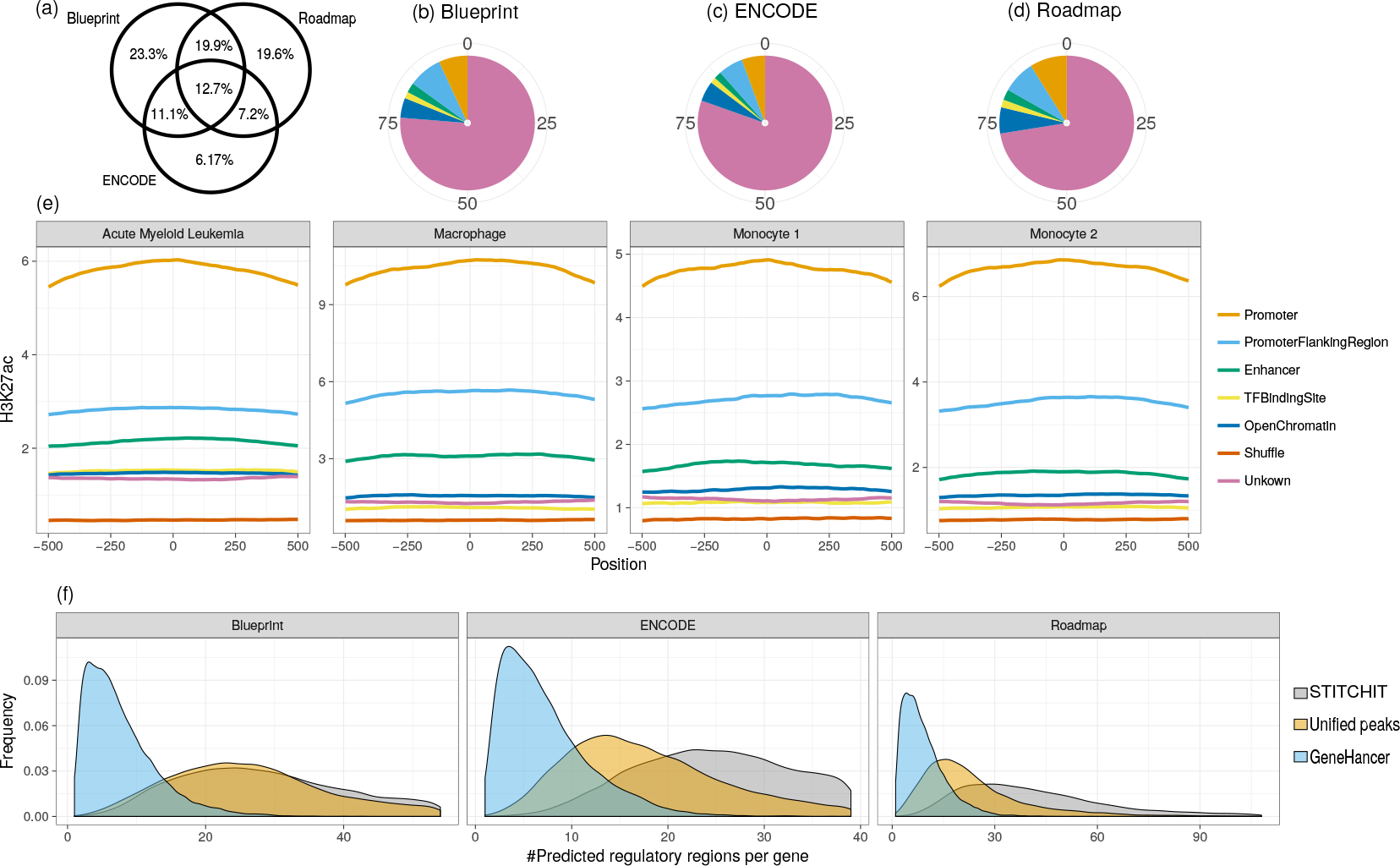
(a) The Venn diagram illustrates the overlap of genes for which a model could be learned using STITCHIT on either one of the datasets used. (b-d) The distribution of a mapping of STITCHIT sites to the ERB is shown. (e) H3K27ac signal shown for randomly shuffled regions, as well as STITCHIT regions split according to the categories obtained from intersecting all STITCHIT segments with the ERB. H3K27ac signal is shown for four Blueprint samples in a window of 1kb centered in the middle of the putative REMs. STITCHIT regions overlapping *Promoter* or *Promoter Flanking Regions* show the highest H3K27ca signal, while the signal in randomly determined regions is the lowest. The largest portion of regions, labeled as *Unknown* have a similar signal intensity as sites labeled as a *TF binding site* or *Open Chromatin*. (f) The density plots delineates the number of predicted REMs per gene, shown separately for the used datasets and tested methods.

Additionally, the trends between the datasets are varying, e.g. we find most associations with STITCHIT on the Roadmap dataset, whereas in case of the UNIFIEDPEAKS approach, we find most associations on Blueprint data.

In Supplementary Fig. S11b, we sketch the distribution of STITCHIT regions around a gene. We find that most STITCHIT regions are located in intragenic positions, that is within the genebody of their respective target gene. Importantly, we do not observe an enrichment in identified sites upstream of the TSS’s of the considered genes, indicating that we do not focus on promoter specific features. The histograms of Fig. 3f illustrate how the number of associated REMs are distributed among their target genes. As expected, the distribution for GENEHANCER is different from that of UNIFIEDPEAKS and STITCHIT. While the latter two reach the maximum, depending on the dataset, between 20-30 REMs per gene, GENEHANCER reaches the optimum at 1-4 predicted sites per gene. Our results also indicate that STITCHIT tends to find more sites per gene than the UNIFIEDPEAKS approach.

### Validation using external data

As STITCHIT can be used to learn interactions to distant sites, we compared the learned interactions to ChIA-PET data for K562 and MCF-7 cells as well as to Promoter-Capture Hi-C data for GM12878 cells. On Blueprint and Roadmap data, about one third of all possible interactions overlap with predictions by STITCHIT and UNIFIEDPEAKS, on ENCODE data about one sixth of all ChIA-PET contacts are retrieved (Fig. 4a). While GENEHANCER constantly finds the smallest overlap, STITCHIT segments overlap with more chromosomal contacts using Blueprint and Roadmap data, while the UNIFIEDPEAKS approach finds more on ENCODE data. There is no clear advantage for any method on the Promoter-Capture data (Supplementary Fig. S12d).

**Figure 4:**
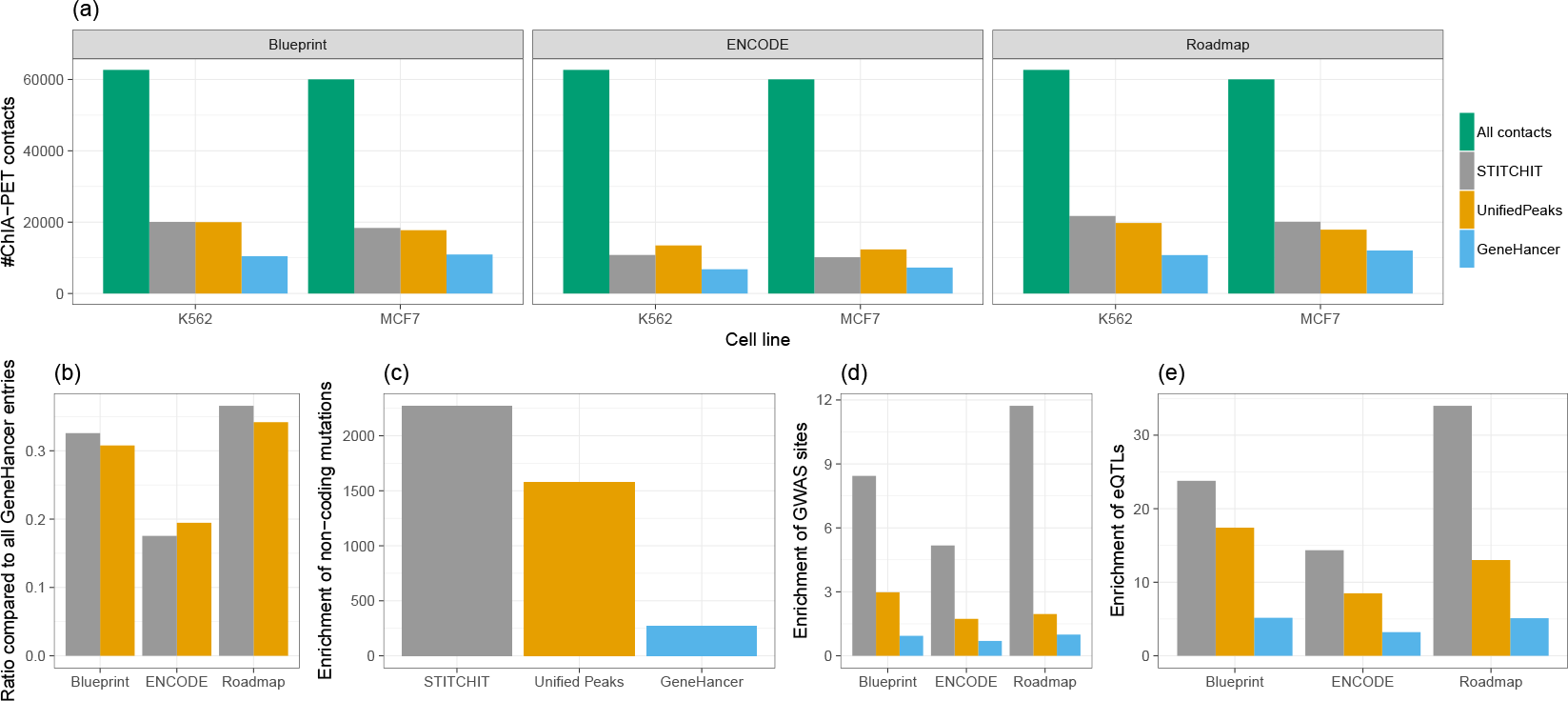
(a) The bar plot indicates how many ChIA-PET contacts are matching the associations of REMs to their target genes across all three datasets and linkage approaches for the K562 and MCF-7 cell lines. With the exception of the ENCODE dataset, STITCHIT retrieves more interactions than both UNIFIEDPEAKS and GENEHANCER. (b) Here, the ratio of recovered entries from the entire GENEHANCER database is shown for STITCHIT and UNIFIEDPEAKS. While UNIFIEDPEAKS retrieves slightly more entries than STITCHIT on ENCODE data, STITCHIT retrieves more known sites on Blueprint and Roadmap data. (c-e) Length normalized enrichment of non-coding mutations (c), of GWAS sites (d) and eQTLs (e). With the exception of (c), which considers only Blueprint data, all other analysis are performed on all datasets and indicate that STITCHIT regions achieve the best score.

Another approach to show the reliability of our predictions is to assess the amount of recovered interactions from GENEHANCER. As shown in Fig. 4b about 32 – 36% of GENEHANCER REMs are retrieved in case of Blueprint and Roadmap respectively and about 18% using ENCODE data. In the latter case the UNIFIEDPEAKS approach finds marginally more overlapping regions, whereas STITCHIT finds more known associations with Blueprint and Roadmap datasets.

The COSMIC database is a collection of known somatic mutations in cancer. Especially mutations occurring in the non-coding part of the genome overlapping REMs might be influencing TF binding in those regions and thus lead to a change in gene-expression that is contributing to the progression of cancer. By overlapping our predictions with all non-coding mutations stored in COSMIC, we suggest to which gene the non-coding somatic mutation can be linked (Supplementary Table S11). In total, we find 1, 006, 848 associations between 883, 111 somatic mutations and 22, 588 distinct genes. STITCHIT is especially well suited for this task, as its overall enrichment of regulatory sites is higher compared to UNIFIEDPEAKS and GENEHANCER regions (Fig. 4c) while at the same time also more mutations can be linked (Supplementary Fig. S12b). Interestingly randomly selected regions overlap more mutations than predicted REMs. This suggests that the sequence in the predicted REMs is conserved and thus likely to be functionally relevant (Supplementary Fig. S12b).

Similarly, we overlapped GWAS sites from the EMBL-EBI GWAS catalog with our predictions suggesting to which gene(s) GWAS hits might be associated (Supplementary Table S11). Overall we find 4697 associations with Blueprint, 888 with ENCODE, and 4588 with Roadmap data covering 3394, 753, and 3366 genes respectively using STITCHIT. Similar to above, STITCHIT yields a better enrichment score than the other two methods (Fig. 4d). Compared to a random setting, all actual regions yield significantly more associations and obtain a significantly better enrichment score (Supplementary Fig. S12a).

Expression quantitative trait loci (eQTLs) are distinct genomic loci that are linked to the expression of genes. We obtained eQTL data from the ExSNP database and overlayed it with our predictions by computing how many eQTL links are correctly overlapping our predictions. As shown in Fig. 4e, STITCHIT regions obtain the best enrichment score for the identification of eQTL overlaps supporting the accuracy of STITCHIT predictions and are also enriched in a randomization experiment (Supplementary Fig. S12c).

### CRISPR-Cas9 validated enhancers for ERBB2 are rediscovered using STITCHIT

*Klann et al.* [26] analysed regulatory elements for the *ERBB2* loci in human SKBR3 breast cancer cells. They have validated the regulatory effect of several candidate DHS sites using CRISPR-Cas9 experiments leading to a set comprising 11 validated regions influencing the expression of *ERBB2* (Supplementary Table S2). We obtained interactions for *ERBB2* using STITCHIT, UNIFIEDPEAKS, and GENEHANCER. Next, we pooled the interactions learned across the three datasets by intersecting them with all potential SKBR3 DHSs (see Methods).

As depicted in Fig. 5, STITCHIT finds 9 of the 11 regions, the UNIFIEDPEAKS approach retrieves 7 regions and GENEHANCER retrieves 6 regions. As an additional support of the predictions, we found that ChIA-Pet data (4DGenome) is linking REMs around *GRB7* to *ERBB2* [50].

**Figure 5:**
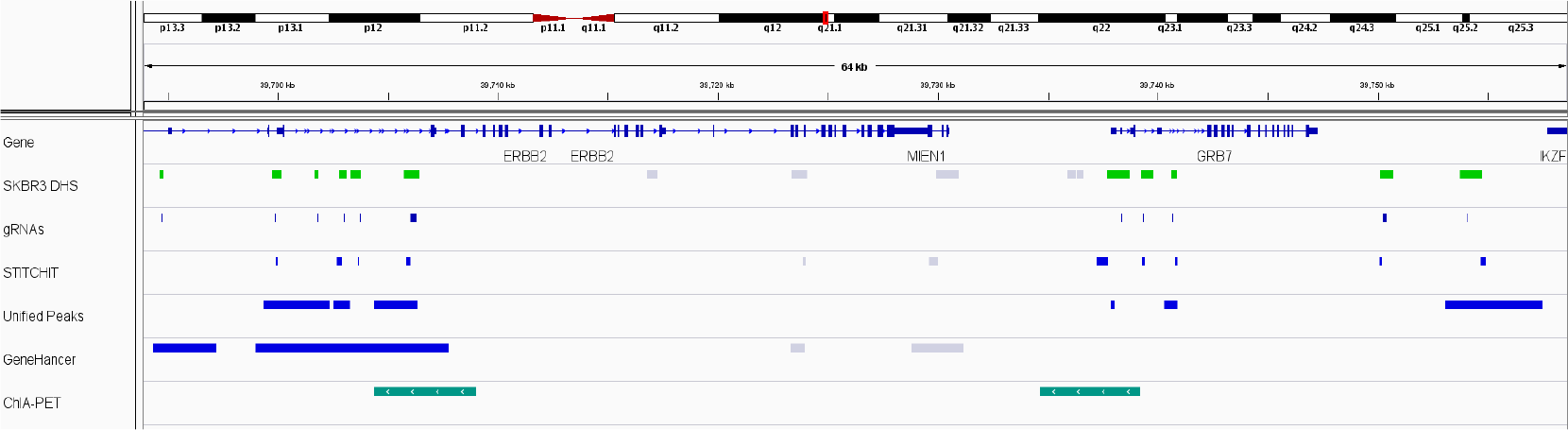
This genome-browser visualization depicts in the gene track the genomic locus of *ERBB2*. The second track shows DHS sites identified in SKBR3 cells and sites highlighted in green have been validated using CRISPR-Cas9 experiments [26], indicated by the gRNA binding sites shown in the track below. The track labeled ChIA-PET shows an interaction obtained from the 4DGenome database. The STITCHIT track contains all STITCHIT regions identified across the three datasets. A blue color indicates that the region overlaps a validated DHS site. The tracks for UNIFIEDPEAKS and GENEHANCER are generated analogously. STITCHIT retrieved 9 sites, UNIFIEDPEAKS identified 7, and GENEHANCER 6.

As shown in Fig. 5, STITCHIT regions are very sharp and are rather a subset of the validated DHSs, especially for those within the gene-body of *ERBB2*, while UNIFIEDPEAKS and GENEHANCER segments are very broad, not providing a good resolution on the regulatory landscape of the chromatin around *ERBB2*. For instance, 6 validated DHS sites are covered by only 2 GENEHANCER segments, rendering the GENEHANCER segments less useful in an exploratory analysis of gene-expression, because the actual sites of importance are not pinpointed precisely. The UNIFIEDPEAKS approach is performing better in terms of this ratio, as only in one instance an identified site overlaps 2 validated DHS, but the mean length of the UNIFIEDPEAKS regions (1449bp) is still longer than the mean length of STITCHIT regions (214bp) and longer than the validated SKBR3 DHSs (562bp). Especially in down-stream applications such as TF-binding predictions or foot-print calling, the more fine grained resolution of STITCHIT is beneficial and can avoid false-positive calls.

### Partitioning of large regulatory elements using STITCHIT

As shown above, the UNIFIEDPEAKS approach produces longer candidate regions than STITCHIT. We define a *split event* as the occurrence of a peak that is divided into several regions by STITCHIT with the additional constraint that the new sub-regions should not be linked to the same gene as the original peak. As depicted in Fig. 6a such *split events* do occur frequently. Note, that for illustration purposes, split events of degree > 10 and smaller than < 2 are not displayed.

**Figure 6:**
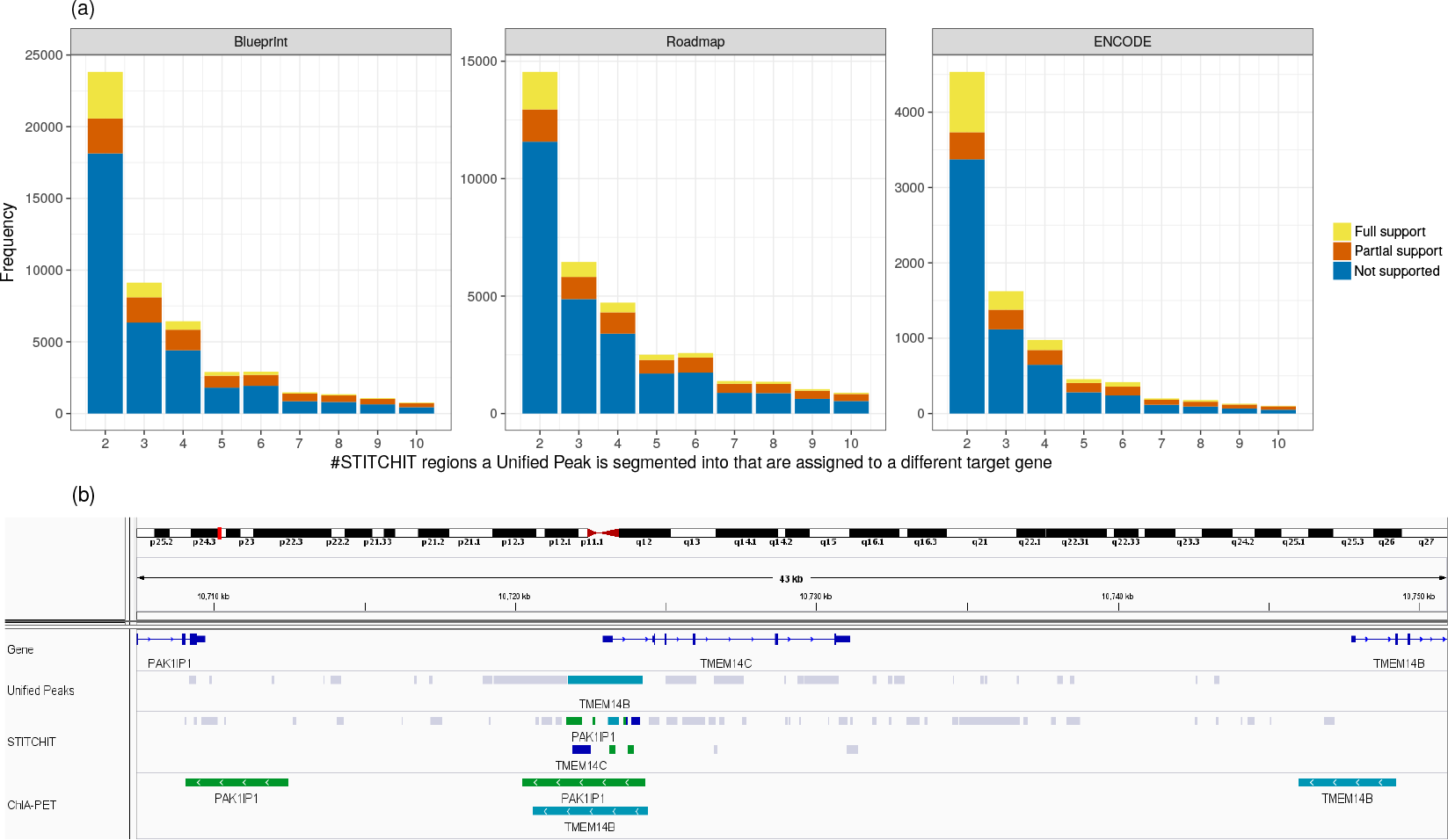
(a) The bar plots indicate on the x-axis the magnitude of a *Split event*, that is the number of differently linked STITCHIT segments a peak is split into. The y-axis holds the frequency for the individual counts. The color code indicates whether STITCHIT associations are fully supported by conformation data, partially supported or not supported at all. (b) Example for a *split event* at the *TMEM14C* locus. At the promoter of *TMEM14C*, a peak that is linked to *TMEM14B* is split into several STITCHIT segments. These are associated to PAK1P1, *TMEM14C* itself, and *TMEM14B*. All STITCHIT associations shown here are supported by ChIA-PET data.

An example is provided in Fig. 6b. Here, a peak is linked exclusively to *TMEM14B* by the UNIFIEDPEAKS method. The peak itself is located around the promoter of *TMEM14C* and covers a total genomxic range of 2497*bp*. STITCHIT divides that peak into segments linked to PAK1P1, to *TMEM14C* itself, and to *TMEM14B*. ChIA-PET data obtained from K562 cells supports the long range interactions to *PAK1P1* and *TMEM14B*. This example, together with the analysis presented in Fig. 6a underlines the ability of STITCHIT to precisely pinpoint regions of regulatory potential and suggests the application of segmenting large REMs, into more refined segments and to reveal their regulatory interactions.

### Exploratory analysis of the regulatory landscape of *EGR1*

To better understand the functional advantage of STITCHIT over UNIFIEDPEAKS, we have investigated the regulatory landscape of *EGR1* in more detail. For *EGR1*, the Spearman correlation achieved by the UNIFIEDPEAKS REMs in gene-expression modelling is 0.55, while STITCHIT regions achieve a correlation of 0.72. Here, we test whether this difference in model performance is also reflected by an improved interpretability of the identified regions regarding the regulation of *EGR1*. In Fig. 7a, we show the identified candidate regions ranked according to the absolute value of the regression coefficients per site (Supplementary Table S8). A striking difference between STITCHIT and UNIFIEDPEAKS is that the latter identifies one large segment (U1: 8970bp) covering 2842bp upstream of *EGR1*, the entire *EGR1* gene as well as 2304bp downstream of *EGR1* TTS. This segment is split up into two regions using STITCHIT: a region downstream of *EGR1* TTS (S1), and into a region within the first exon of *EGR1* (S2). As shown by the DNase1-seq signal tracks in Fig. 7a, STITCHIT region S1 and S2 do overlap DNase1-seq signal in sample *C0010KB*, in which *EGR1* is expressed, whereas they lack signal in *C005VG11*, where *EGR1* is not expressed. It is likely that this difference between STITCHIT and UNIFIEDPEAKS is the main reason for the observed performance difference.

**Figure 7:**
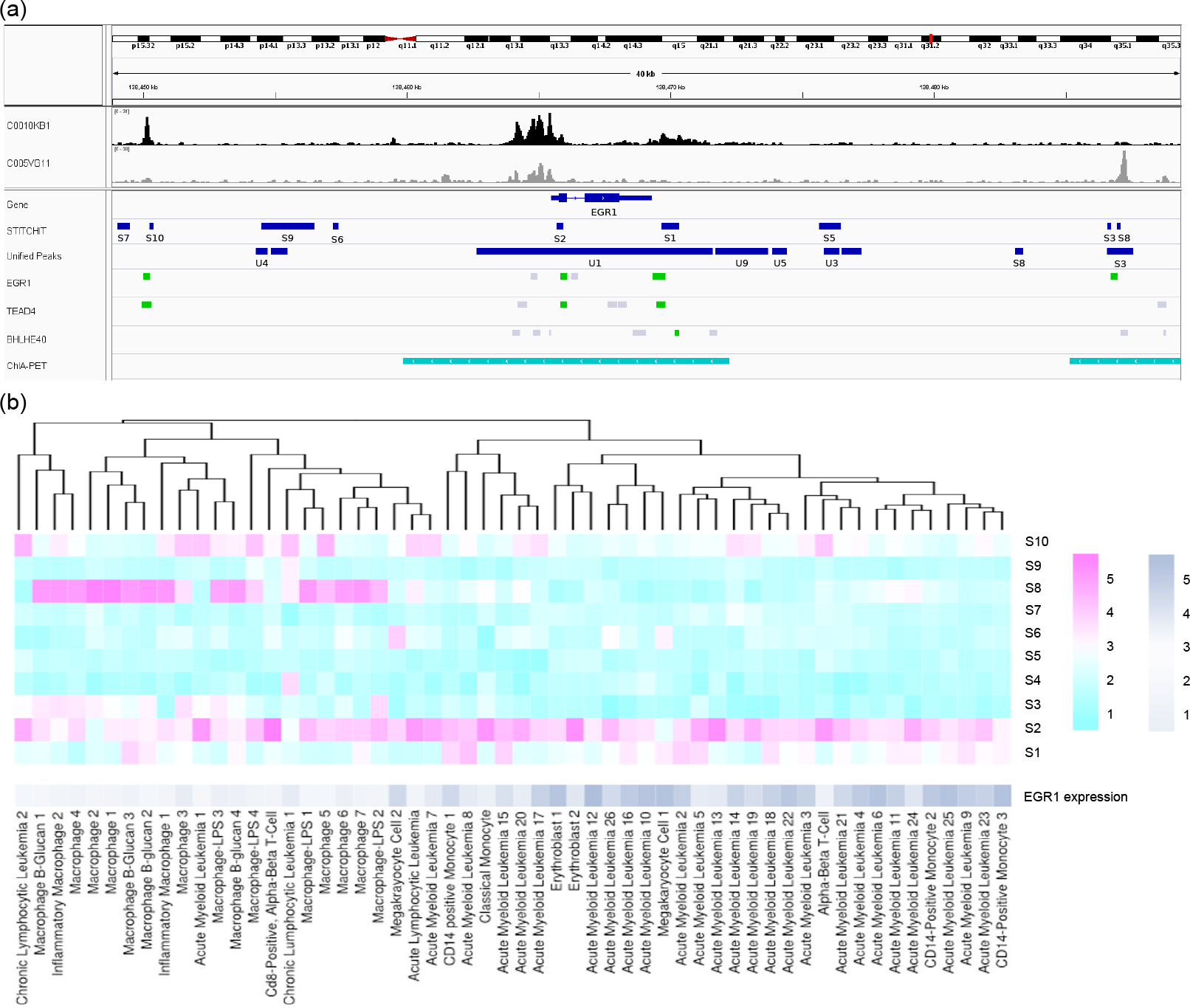
(a) Genome browser tracks describing the regulation of *EGR1*. Track *C0010KB1 (black)* exemplifies the DNase1-seq signal for a sample where *EGR1* is expressed, whereas track *C005VG11 (gray)* illustrates the case where *EGR1* is not expressed. For the last three tracks, a green segment indicates that the respective TF ChIP-seq peak overlaps with a STITCHIT region. (b) Heat map that is clustered according to the DNase1-seq signal in the candidate REMs *S*1…*S*10 identified by STITCHIT, the gene-expression of *EGR1* is not used for the clustering itself and shown for illustration purposes only. The data has been log transformed with a pseudo-count of 1. Two major clusters can be observed corresponding to samples where *EGR1* is expressed and to those samples where *EGR1* is not expressed. The heatmap shows the *log*_2_ of read counts for DNase1-seq, and *log*_2_ of TPM for gene-expression, respectively.

Another interesting association can be observed for S3 and S8, which also overlap a segment identified with UNIFIEDPEAKS (U2). S3 has the strongest negative regression coefficient identified by STITCHIT for *EGR1* and indeed this region (as well as S8) shows signal in *C005VG11* but not in *C0010KB*, supporting the role of the regions as an active repressor of *EGR1*. The link of S3 to *EGR1* is further supported by ChIA-PET data.

While these examples provide insights on the level of individual samples, we have considered the DNase1-seq signal within all identified STITCHIT regions and used it to cluster the Blueprint samples (Fig. 7b). Using only the signal within the candidate regulatory sites, an almost perfect clustering into samples according to *EGR1* expression levels could be obtained. The clustering can be used to assess the cell-type specificity the suggested regions.

We further hypothesized that regions with the strong reg. coefficient should be functionally active, e.g. containing TF binding sites. We used Fimo [17] to predict TF binding in REM S1 (Supplementary Table S9) and REM S3 (Supplementary Table S10), the regions with the strongest positive and negative association predicted by STITCHIT, respectively. For the top hits (ranked by Fimo q-value), we checked ENCODE for TF ChIP-seq data. Indeed, we found a peak of TEAD4 and BHLHE40 in S1, which are ranked as first and fourth hit by FIMO. For the second and third TF, namely SP8 and BCL6, as well as for most TFs binding in S3 no ChIP-seq data was available at ENCODE. However, EGR1 ChIP-seq data, which was predicted to bind in S3, is available. Surprisingly, EGR1 does not only bind to S3, but also to S1, S2 and S10, a region uniquely identified using STITCHIT that is about 15*kb* upstream of the EGR1 gene. Overall, the ChIP-seq analysis not only suggests that the regions identified with STITCHIT are functionally relevant, it also suggests a potential self regulatory role of EGR1 by binding REMs associated to its own gene.

### STITCHIT suggests genes and REMs related to Doxorubicin resistance

We use a recently published genome-wide doxorubicin CRISPR-Cas9 resistance screen [56] to show another possible application of STITCHIT, which is the association of experimentally highlighted regions to their most likely target genes. As stated above, the CRISPR-Cas9 screen led to 332 non-coding target sites of 226 validated gRNA sequences. In total, we found 111 putative REMs overlapping with a non-coding gRNA binding site, which could be linked to 100 different genes using STITCHIT (Supplementary Table S6). Importantly, STITCHIT obtains significantly more REMs overlapping the gRNA binding sites than randomly selected regions (Supplementary Fig. S13). Two genes targeted by gRNAs via regulatory element interactions have been previously reported to be associated with doxorubicin resistance: *MMP9* and *ATF3* (p-values 0.03 and 0.08, respectively). Doxorubicin is a chemotherapy medication used to treat different forms of cancer. It induces double-strand DNA breaks and triggers DNA damage associated cell cycle arrest and apoptosis pathways, for example via MAPK/ERK pathway [51, 5]. In myocytes, an increase of Myocardial Metallo Proteinases (MMPs) expression through increasing ROS formation induced by doxorubicin was observed [49]. At position chr20:45993823, overlaps between a gRNA hit and STITCHIT predictions assume a regulatory site of MMP9 (Supplementary Table 6), being in agreement with the finding of Spallarossa et al [49]. ATF3 is a TF known to be involved in cellular stress response and is enriched in cells exposed to stress signals [20]. Under doxorubicin treatment, it has been reported that ATF3 affects cell death and cell cycle progression, however it is unclear whether the factor acts as a negative or positive regulator [35, 37]. Nobori et al. claim that ATF3 plays a pivotal role as transcriptional regulator in the process of doxorubicin-induced cytotoxicity via an ERK-dependent pathway. STITCHIT identified a REM of *ATF3* in proximity of position chr1:212621320, a gRNA target site. Both *MMP3* and *ATF3* are also supported by GENEHANCER data as well as ChIA-PET interactions.

We observed that in 33% of all cases, intronic or exonic gRNA-target regions of a gene function as REMs for upstream or downstream located genes, as is the case for *APOL5* and *FANCA* introns, associated to *RBFOX2* and *ZNF276*, respectively (Supplementary Fig. S14a,b). Moreover, STITCHIT proposes single REMs being linked to multiple genes, demonstrating that one gRNA can lead to multiple correlated interrogations. For instance, the gRNA target site chr3:147409323 was linked to altered gene-expression of *ZIC1* as well as *ZIC4* (Supplementary Fig. S14c). A REM targeting the genomic location chr3:147407359-147412176 has also been identified by GENEHANCER (GRCh38/hg38, GH03J147407). In an extreme example, STITCHIT indicated one gRNA target site chr9:92116400 to be linked to four pseudogenes simultaneously (MTATP6P29, LINC00475, AL354751.3 and AL354751.1, Supplementary Fig. S14d). In sum, the combination of STITCHIT and CRISPR-Cas9 screens has the potential to prioritise non-coding gRNA hits in known and unknown REMs.

## Discussion

Here, we presented STITCHIT, a novel method to define and link REMs to their target genes. We compared STITCHIT against three related strategies: (1) an unsupervised linkage of DHS sites [42], (2) considering known regulatory elements from the GENEHANCER database [11], and (3) combining sample specific DHSs in large datasets, referred to as the UNIFIEDPEAKS approach. The latter is conceptually similar to peak aggregation strategies used by *Hait et al.* [21] or *Shooshtari et al.* [47]. In *Hait et al.* non-promoter DHS sites with a signal enrichment of ≥ 1RPKM in at least 30 samples are considered as a candidate REM. By default, each gene is linked to the 10 closest DHSs. In *Shooshtari et al.* replicate information is used to compute an overlap between DHSs. The overlap information is used in a clustering algorithm to group individual peaks into clusters. Cluster borders are defined by the extreme positions of the included DHSs. The UNIFIEDPEAKS approach is following the idea of merging the peak signal, but due to the sample specific overlap with the signal, tissue specific information is maintained as far as possible. By relying on the signal of an epigenetic signature, instead of peak calls, STITCHIT does not only omit the non-trivial peak calling step, but it is also able to pick up even gene-specific enrichment’s in the signal, which would be either omitted, or not accurately resolved by peak callers.

In gene-expression prediction experiments, STITCHIT regions achieve a better agreement between predicted and measured gene-expression than related approaches (Fig. 2a, Supplementary Fig. S8a). However, the performance of predictive models alone does not proof that the identified regions truly play a role in gene regulation. We stress that STITCHIT associations do not imply causation. Thus, we can not distinguish whether the accessibility of certain regions is driving expression of a gene, or whether it is a consequence of that gene being expressed. Also indirect associations, which could for be caused by co-regulation of genes, can not be avoided. Therefore, it is especially important to characterize the predicted REMs further.

To do so, we intersected them with the ERB (Fig. 3c-e). The ERB segments the genome according to DNase1-seq and HM data for 18 different cell-types, mostly cell-lines. Due to the limited data and biological diversity, as stated by the authors of the build themselves, the ERB is not exhaustive [59]. Thus, it it is not surprising that the largest portion of STITCHIT REMs does not overlap with an annotated regulatory region. To show that these non-overlapping REMs are still relevant, we assessed the H3K27ac signal within those sites (Fig. 4a). Compared to randomly sampled regions the HM signal is higher across the STITCHIT REMs supporting that they are of regulatory importance. On top of that, STITCHIT obtains a better recall value for retrieving known regulatory elements from the GENEHANCER database than the UNIFIEDPEAKS approach, another indication for the reliability of STITCHIT predictions. Furthermore, up to 25% of the predicted interactions have been experimentally determined using chromatin conformation data. Importantly, the experimental data is also validating a large portion of *split events*. This highlights another potential application of STITCHIT, the segmentation of super enhancers, large genomic regions with enhancer function.

Throughout this study, we observed that especially on large heterogeneous datasets, such as the Roadmap dataset, the peak-independent generation of REMs shows clear advantages over the peak-based strategies. This is especially obvious comparing our results obtained for Blueprint and Roadmap data. While the Blueprint dataset is composed of primary cells related to the hematopoietic lineage, the Roadmap dataset is more diverse and also comprised of tissue samples. On the more homogeneous Blueprint data, STITCHIT and UNIFIEDPEAKS identify almost the same number of segments with similar length (Fig. 2b,c). In contrast to that, on Roadmap data, STITCHIT selects more, but shorter REMs than UNIFIEDPEAKS (Fig. 2b,c, Fig. 3a). This difference is also reflected by the performance of the gene-expression models. The overall difference between STITCHIT and UNIFIEDPEAKS is more pronounced for Roadmap, than for Blueprint data (Fig. 2a, Supplementary Fig. S8). The most likely explanation for this behavior is that due to the high variance in the Roadmap data, merging peaks introduces a loss of specificity, by removing the information of the exact genomic location of accessible chromatin (Supplementary Fig. S1). On the less variable Blueprint dataset, this seems to be less of an issue. STITCHIT is able to resolve the sample and tissue specific variance, therefore obtaining better results on Roadmap data compared to the UNIFIEDPEAKS method. However, we note that STITCHIT is not able to outperform the UNIFIEDPEAKS approach on ENCODE data in terms of recall from the GENEHANCER database as well as in the overlap with ChIA-PET data, which is likely to be due to the low number of samples contained in the ENCODE dataset. Compared to Blueprint and Roadmap data, this might also explain why much fewer REMs have been predicted in total (Fig. 3a). This shows how closely the analyzed data is linked to model quality and reliability.

## Conclusions

Our novel method STITCHIT solves the combined task of identifying potential REMs, and linking them to their putative target genes at the same time. This is achieved by combining epigenetics and gene-expression data to identify a set of potential REMs considering the signal of the epigenetics data at hand, instead of pre-selected sites of enrichment. Hence, the peak calling step can be omitted. Subsequently, STITCHIT regions are refined using a two-level learning approach and a confidence score for each REM is computed. Our modeling approach allows a distinct REM to influence multiple genes. In this work, uniformly processed DNase1-seq and RNA-seq data from IHEC is used, however our method is conceptually not limited to DNase1-seq data as a carrier of epigenetic information, but also works with ATAC-seq, FAIRE-seq or ChIP-seq data.

We have compared STITCHIT against related strategies that are based on the integration of peaks or on known REMs, as stored in the GENEHANCER database [11], and show that STITCHIT is not only able to learn more sites with regulatory potential than the other methods, while achieving a superior explanatory power of gene-expression. It also performed well in various validation experiments.

Furthermore, we illustrate how STITCHIT can be used in an exploratory manner to elucidate the regulation of a distinct gene exemplary for *EGR1*, and we delineate how STITCHIT can facilitate the interpretation of REMs predicted with CRISPR-Cas9 in the non-coding part of the genome. STITCHIT is efficiently implemented in C++ and freely available on github: www.github.com/schulzlab/STITCHIT.

We believe that STITCHIT paves the way for a seamless integration of the wealth of epigenetics data being produced and allows an easy-to-use analysis of transcriptional regulation on the gene-level.

## Methods

### Preprocessing

Paired DNase1-seq and RNA-seq data was downloaded from the ENCODE data portal for 41 ENCODE and 110 Roadmap samples. Upon granted access, we obtained 56 paired DNase1-seq and RNA-seq samples from Blueprint. An overview is provided in Table 1 on sample numbers and tissue/cell-type diversity. Supplementary Table S1 lists all data accession numbers. Paired samples are required as they are expected to have a better correlation between chromatin structure and gene-expression, as both samples originate from the same donor. Details on data processing as well as used command calls are provided in Supplementary Sec. 1.

**Table 1:** Overview on the data used in this study.

|  | **Blueprint** | **Roadmap** | **ENCODE** |
| --- | --- | --- | --- |
| #paired samples | 56 | 110 | 41 |
| #different cell-types | 13 | 33 | 25 |
| primary cells only | Yes | No | No |

Further, we obtained H3K27ac data in *wig* format from the Blueprint data portal for four samples (C0011IH1, S00C0JH1, S00XUNH1, C0010KH1, see Supplementary Table S1). Also, we downloaded REMs contained in the GENEHANCER database from the GeneLoc website [40].

### Overall workflow and conceptual idea

Conceptually, we pursue the idea to identify regions in large genomic intervals around a gene of interest that can be associated to the genes expression variation across many samples. To identify these regions, we utilize paired epigenetics and gene-expression data. The STITCHIT algorithm uses the actual signal of the epigenetics data to highlight segments of the data showing signal variation that can be used to separate samples according to the target genes expression. Thus, the peak-calling step can be omitted and the two tasks of identifying regulatory sites and their linkage to targets are solved simultaneously. To refine the list of putative REMs identified by STITCHIT, we apply a two-level learning approach that is detailed below. This allows us to judge the explanatory power of the found regions for gene-expression and to obtain a p-value for the significance of each identified region. The workflow of the proposed methodology is depicted in Fig. 1(a), details are provided in Supplementary Fig. S2.

### The STITCHIT algorithm

In the following, we are given a dataset *D*_*g*_ with *m* rows, corresponding to the samples, and *n* columns representing the epigenetic signal at base pair resolution around the target gene *g*. Further, to each row, we assign a class label, indicating whether the corresponding sample is associated with a high, medium or low expression value (*C* = 0, 1, 2). Note that also a two level classification was used here (*C* = 0, 1), depending on the results of the POE method [14] (c.f. Supplementary Section 3). The algorithm is implemented such that any number of distinct class labels, not exceeding the number of samples, (|*C*| ≤ *m*) can be used. With *C*_*k*_ we relate to all rows to which we assigned class label *k* ∈ *C*.

A segment *s* has a start point *i* and an end point *j*, where 1 ≤ *i* ≤ *j* ≤ *n*. We call *S*_*g*_ a segmentation of *D*_*g*_, if it contains a set of non-overlapping segments that covers the whole range from 1 to *n*. The two trivial cases would be a segmentation consisting of only a single segment with start point *i* = 1 and end point *j* = *n* or the segmentation containing *n* segments, where each segment only contains a single column, *i.e.* a single base. The former would contain no information about the class labels, while the latter would consist of a large set of noisy segments which result in bad features for the learning step that is based on the segmentation. Our goal is to provide a small set of robust features for the learning step. We achieve this by joining adjacent base pairs to segments, such that the variance between the epigenetic signals of base pairs that are contained in a segment is low, w.r.t. the class label. The optimal segmentation according to the score we define below, finds a trade-off between the number of segments and the variance.

To score a segmentation, we propose an information theoretic score based on the Minimum Description Length (MDL) principle [19]. MDL is a practical instantiation to Kolmogorov complexity [27] and thus belongs to the class of compression-based scores. Formally, given a model class 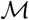, MDL identifies the best model 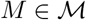 for data *D* as the one minimizing

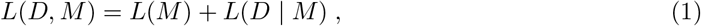

where *L*(*M*) is the length in bits of the description of the model *M*, and *L*(*D* | *M*) is the length in bits of the description of the data *D* given *M*. This is known as two-part, or crude MDL. In essence, we try to find the simplest model that can explain the data well. We follow the convention that all logarithms are base two, since the length of the encoding relates to bits, and define 0 log 0 = 0. In this work, we use MDL to balance our segmentation between having too few segments and running at risk of missing structure in the data and finding too many segments, which contain spurious information and make the post-processing infeasible.

From now on, we consider the model class of segmentations 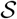 from which we want to find the optimal segmentation, 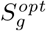 that is

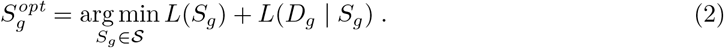

In particular, we encode a segmentation *S*_*g*_ as follows

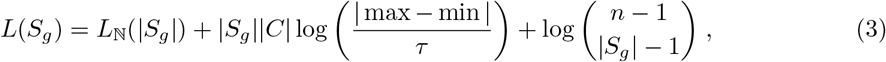

where |*S*_*g*_| denotes the number of segments, *L*_N_ is the universal prior for integer numbers [19], |*C*| is the number of class labels and *τ* ≤ 1 is the data resolution.

First, we encode the number of segments, then for each segment per category the associated mean value by assuming it lies between the minimum and the maximum value in the data and last the complexity to select |*S*_*g*_| segments from possible *n* segments.

To encode the data given a segmentation, we simply sum over the costs per segment

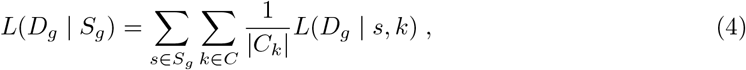

where |*C*_*k*_| corresponds to the number of rows associated with class label *k*.

To encode the costs for a specific segment and the data associated with class *k*, we encode the error assuming a Gaussian distribution. Using 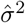 as the sample variance over the data corresponding to segment *s* and class label *k*, we get (compare [19])

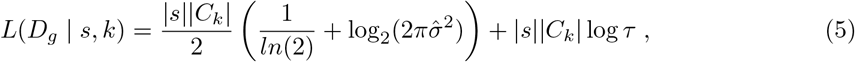

with |*s*| being the length of the segment.

To find the optimal segmentation 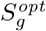, we use dynamic programming [2]. In essence, we start with a segmentation containing only a single segment. Then we iteratively compute the best segmentation containing *i* segments based on the best segmentation containing *i* − 1 segments for *i* ∈ {2, …, *n*}. Lastly, we select 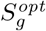 among the optimal segmentations for each possible number of segments. The runtime complexity of this algorithm is 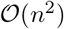. By selecting a minimum segment size of *β* and partitioning the search space into *l* chunks, we can run each chunk in parallel and the total runtime complexity reduces to 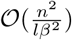. In our experiments, we use *β* = 10 and set *l* to 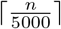, which makes the algorithm feasible to be applied on large genomic intervals. Here, we have considered 25*kb* upstream of a genes’ Transcription Start Site (TSS) and 25*kb* downstream of a genes’ Transcription Termination Site (TTS).

An example is provided in Supplementary Sec. 2.

### Selection of candidate regulatory elements

Those segments that are associated to the observed expression changes need to be extracted from 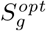. Thus, for all segments 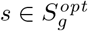 we compute both Pearson and Spearman correlation between the epigenetic signal in *s* across all samples *m* and the continuous expression values of the targetgene *g*. We select all segments that achieve either a Pearson or Spearman correlation with a significance threshold of *p* ≤ 0.05. We apply the same filtering to the alternative methods introduced below.

### Refinement of selected regions using two-level learning

STITCHIT provides for all selected segments *s* ∈ *S*_*opt*_ a matrix *X* holding the epigenetic signal within these regions. The *m* rows of *X* denote the samples, the *n* columns refer to the regions selected by STITCHIT. To further refine the suggested regions for a distinct gene *g*, we first train a linear model using elastic net regularization, as implemented in the glmnet R-package [12]. Here, we are utilizing the DNase1-seq signal within candidate REMs (*X*) to predict the expression of *g*, stored in *y*. The grouping effect results in a sparse regression coefficient vector. However, correlated features prevalent in this application, will be maintained. This is achieved by combining both the Ridge and the Lasso regularizers:

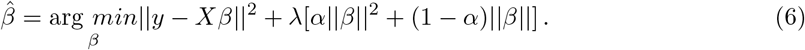

Here, *β* represents the feature coefficient vector, 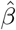 the estimated regression coefficients, and *λ* controls the total amount of regularization. Both the input matrix *X* and the response vector *y* are log-transformed, with a pseudo-count of 1, centered and normalized. The parameter *α*, which is optimized in a grid search from 0.0 to 1.0 with a step-size of 0.01 controls the trade-off between Ridge and Lasso penalty.

As previously performed by Schmidt et al [44], model performance is assessed in terms of Pearson and Spearman correlation as well as using the Mean Squared Error (MSE) between predicted and measured gene-expression. The test is performed on a hold-out test dataset in a ten-fold outer Monte Carlo cross-validation procedure where 80% of the data are randomly selected as training data and 20% as test data. The parameter *λ* is fitted in a six-fold inner cross-validation using the *cv.glmnet* procedure. The parameters’ final value is determined according to the minimum cross-validation error which is computed as the average MSE on the inner folds *(lambda.min)*.

Significance of the correlation between predicted and measured gene-expression is corrected using the Benjamini-Yekutieli correction [3], which is designed to account for dependency between the tests [21]. Only models with a q-value ≤ 0.05 are considered for interpretation of the selected regions. For those models, we refer to all features with a median non-zero regression coefficient across the outer folds by *X*_*NZ*_.

In a second learning step, as performed by *Hait et al* [21], we train an Ordinary Least Squares model (OLS) on the pre-selected features *X*_*NZ*_ predicting *y* and report the regression coefficients *β*_*OLS*_ as well as the *p*-values per feature for downstream analysis:

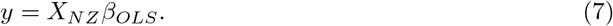

The OLS model allows for a simple comparison of regression coefficients *β*_*OLS*_ across genes, as there is no bias introduced by the regularization, and provides a straight forward way to compare individual regions. All regions and model coefficients used for interpretation and validation are obtained from the OLS models (Supplementary Fig. 3).

### Alternative approaches to identify and to link REMs to genes

We compare the REMs identified with STITCHIT to those obtained with three alternative approaches (Supplementary Fig. S4): (1) an unsupervised, window based aggregation of DHSs per gene and per sample, (2) taking the union of DHSs across all samples (UNIFIEDPEAKS), and (3) considering known REMs from the GENEHANCER database. For approaches (1) and (2), we have used the DHSs called with JAMM (see Methods). Command line arguments along with further details on how to produce the respective scores are provided in Supplementary Section 4. We applied exactly the same two-level learning paradigm for approaches (2) and (3) as described above for the regions identified with STITCHIT. The unsupervised linkage (1) is not considered for interpretation purposes.

### Unsupervised integration of peaks per sample

Similar to work by others [44, 16], we determine for each gene *g* in each sample *i* considering a predefined window 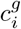, how many DHS sites are located within this window *c*^*g*^, how long are the accessible regions 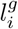, and we aggregate the signal intensity within the selected DHSs 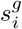. The contribution of each DHS *p* is also weighted by its distance *dist*(*p, g*) to the TSS of gene *g* following an exponential decay. Details are provided in Supplementary Sec. 4.

### Unified peaks

Here, we generate consortia specific aggregations of all DHSs called with JAMM. Overlapping sites are merged using the *bedtools merge* command. Thereby, we obtain a set of regions representing all accessible sites within one consortia. Using the *bigwig* files generated with DEEPTools and the *libBigWig* library (https://zenodo.org/record/45278), we compute the DNase1-seq signal within the merged peaks for each sample. Next, we test for all candidate peaks within a distinct window *w*, here *w* = 25*kb* upstream of a genes TSS and downstream of its TTS, whether there is a significant correlation (*p* ≤ 0.05) between the DNase1-seq signal within the peak and the expression of the gene. All merged peaks passing this test 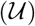 are considered for the two-level learning described above. We refer to this as the UNIFIEDPEAKS approach.

This approach is conceptually similar to the peak aggregation approaches pursued by Hait et al [21] and Shooshtari et al [47].

### GeneHancer

For all REMs obtained from the GENEHANCER database, we calculate the sample specific DNase1-seq signal within each region for each gene, using the *libBigWig* library. Note that a window or distance cut-off is not required here since each region is already assigned to its putative target gene. Considering that the GENEHANCER database is comprised of REMs originating from many different sources identified with a plethora of assays and molecular signatures, we perform the same correlation based test as above to identify a subset 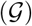 of regions with sufficient correlation between the DNase1-seq signal and the gene-expression of the respective target gene.

### Implementation & Usability

We have implemented the STITCHIT algorithm, the UNIFIEDPEAKS approach, and a linking using previously defined regions (e.g. from GENEHANCER) using C++. Each linkage method is available as a separate executable in our repository. The code can be easily build using CMAKE (version ≥ 3.1) and requires a C++11 compiler supporting *openmp* for parallel execution of STITCHIT. We have thoroughly tested STITCHIT using googletest.

### Code Availability

All scripts except for the unsupervised peak linkage, which is available at www.github.com/schulzlab/TEPIC [43], are available at www.github.com/schulzlab/STITCHIT.

### Validation of putative regulatory regions

#### Regulatory build overlap

We used the terms: Open chromatin, Promoter, Promoter Flanking Region, TF binding site and Enhancer from the Ensembl Regulatory Build (ERB) [59] (release 86), to compare predicted REMs to an established regulatory annotation of the genome.

#### Overlap with H3K27ac data

We selected the top 10, 000 STITCHIT REMs, ranked by their OLS p-values. Also, we obtained 10, 000 random regions of similar size using the BEDTools *shuffle* command excluding the original positions to obtain a set of randomly distributed sites. Additionally, we obtained the top 10, 000 regions according to their label determined by the Regulatory build overlap. Next, we obtained the H3K27ac signal for four Blueprint samples (see Data) in 1*kb* windows centered in the middle of the candidate REMs and visualized the data in R.

#### Overlap with GENEHANCER

Using BEDTools *intersect* we computed the overlap between all candidate regulatory sites identified with STITCHIT and the two-level learning with all unique entries contained in the GENEHANCER database that are within the searched 25*kb* search window and downstream of each gene (193, 298 distinct regions). The same is done for regions based on the UNIFIEDPEAKS approach, thereby assessing how many known REMs from GENEHANCER can be recovered.

#### Overlap with non-coding mutations from the COSMIC database

The COSMIC database is a vast collection of somatic mutations occurring in cancer. We assembled a collection 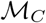 containing non-coding mutations, extracted from the file *CosmicNCV.tsv.gz*. Using 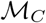, we compute a length normalized score *e*_*C*_ describing the enrichment of mutations in the REMs as

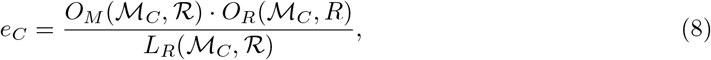

where 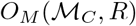 is the number of mutations in 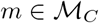 overlapping a candidate REM 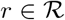, 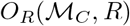 is the number of regions 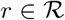 overlapping a mutation 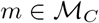, and 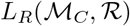 is the total genomic space covered by all regions 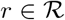 overlapping a mutation 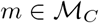. Normalizing by 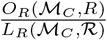 is necessary to account for the length difference between STITCHIT, UNIFIEDPEAKS, and GENEHANCER segments. This normalization factor, which we call resolution, is large, if 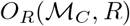 is big, that is there are many overlapping REMs, and 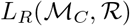 is small, that is the covered genomic space is small. The resolution is small, if 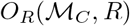 is small, that is there are only a few overlapping REMs, and 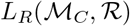 is big, that is the covered genomic space is large. Thus, the normalization adjusts the number of retrieved mutations such that if two methods identify the same number of mutations 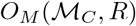, the method with a better resolution, i.e. there are many distinct REMs covering only a small part of the genome, is preferred.

Here, the score *e* is computed for all REMs suggested by all methods as well as for ten randomly shuffled region sets containing the same number of regions as the original sets, respectively. The COSMIC analysis was only performed on Blueprint data due to the large number of included acute myeloid leukemia samples.

### GWAS hits

We compiled a collection 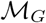 comprising all GWAS sites contained in the EMBL-EBI GWAS Catalog [32]. Using 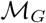, we compute a length normalized score *e*_*G*_ denoting the enrichment of GWAS hits in candidate regulatory sites as above:

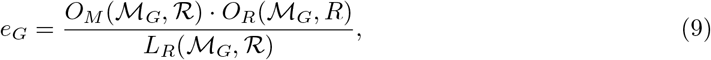

where 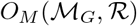 is the number of mutations in 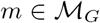 overlapping a candidate REM 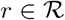 and 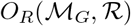 refers to the number of regions 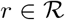 overlapping overlap a GWAS hit 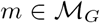.

### eQTL analysis

We obtained all eQTLS 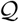 contained in the ExSNP database ([58]), which we mapped to hg38 using dbSNP [46]. To assess how many of those eQTLs overlap regulatory sites that are assigned to the same target gene as the eQTL, we compared the gene-locus assignment from all 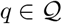 with our predictions in terms of a length-normalized enrichment score *e*_*Q*_:

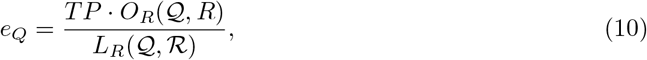

where *TP* refers to true positives, i.e. eQTLs 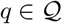 that overlap a suggested REM that is linked to the same gene as *q* itself, 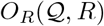 refers to the number of regions 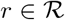 that overlap any eQTL site 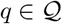 and 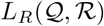 denotes the entire genomic space covered by overlapping REMs.

### ChIA-Pet & Capture Hi-C data

ChIA-Pet data for K562 and MCF-7 was downloaded from the 4DGenome database [50] and lifted to hg38 using the UCSC liftover tool. Capture Hi-C data for GM12878 was obtained from *Mifsud et al.* [34] and also lifted to hg38. The conformation data allows us to calculate how many contacts captured by the ChIA-Pet or Promoter Capture data are matching to the associations inferred by the approaches tested in this study. To match chromatin interaction data to our suggested REMs, we consider the entire gene-body of the linked gene as the second coordinates. We count a REM as contained in the conformation datasets if either the gene or the coordinate of the associated REM overlaps one coordinate of the verified interaction and the second coordinate of the interaction site overlaps the remaining coordinate of the association. Interactions that could not be detected by any of the tests, due to an exceeding genomic distance to the target gene or due to the absence of any DNase1-seq signal in the cell line related to the sample, are excluded from consideration.

### Splitting peaks through STITCHIT

From the overlap between UNIFIEDPEAKS 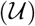 and STITCHIT 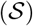 regions it can be computed into how many STITCHIT segments *s* a peak 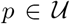 is split into. We refer to the instance that *p* is divided into several segments *s* as a *split event*. The *degree of a split event* denotes the number of STITCHIT segments *s* a peak 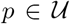 is segmented into. Within this counting procedure we also impose that any *s* overlapping *p* needs to be linked to a different gene *g* than *p*, while any STITCHIT segment *s* can be assigned to the same target gene *g*′ as long as *g*′ ≠ *g*. In addition, we quantify how many *split events* are supported by conformation data. To this end, for each *split event*, we assess how many STITCHIT segments overlap a matching genomic contact obtained from ChIA-Pet or Capture Hi-C data. If all STITCHIT regions are supported, we call a split *fully supported*, if not all but at least one region is supported we call it *partially supported*.

### Validation using Crispr-Cas9 data for ERBB2

*Klann et al.* validated several REMs for ERBB2 using CRISPR-Cas 9 experiments [26]. We assembled a list of these validated regions (c.f. Supplementary Table S2) and generated a bed file containing all validated DHS sites in SKBR3 breast cancer cells using the original DHS calls from *Klann et al*, obtained from the GEO [GSE96876] [26].

We have calculated the overlap between all DHS sites identified in SKBR3 cells and predicted REMS using BEDTools across all datasets. If more than one predicted region overlapped a DHS site, only the most significant one is kept in the intersection. Supplementary Tables S3, S4, and S5 provide a list of the overlapping STITCHIT, UNIFIEDPEAKS, and GENEHANCER regions respectively. Additionally, we have used CHIA-Pet data from 4DGenome to illustrate the interaction between downstream enhancer regions with ERBB2, visualized in IGV [39].

### Analysis of Doxorubicin resistance in hTERT-RPE1 cells using CRISPR experiments

Here, we use a genome-wide CRISPR perturbation library consisting of partially randomized de-generated oligonucleotides (5’-NNDNNNNNHNNNNHDHNVVR-3’) with flanking 3Cs homology regions which was created using ssDNA of template-plasmids and site-specific mutagenesis targeting coding and non-coding regions of the human genome in hTERT-RPE1 cells from ATCC (CRL-4000) [56]. In total, 226 unique gRNAs could be mapped to the coding and non-coding part of the genome, resulting in 332 unique genomic target sites. In order to link putative regulatory sites detected by the gRNAs to genes, the 332 distinct genomic targets sites were extended by a window of 100bp up and downstream of the gRNA binding site. The extended windows are intersected with STITCHIT regions. All predicted non-coding interactions as well as additional ChIA-Pet and GeneHancer support are shown in Supplementary Table S6.

## Declarations

### Ethics approval and consent to participate

Not applicable.

### Consent for publication

Not applicable.

### Availability of data and materials

Raw data (Supplementary Table S1) can be downloaded from the ENOCDE data portal for both ENCODE and Roadmap data. To gain access to raw data files from Blueprint, a data access application needs to be submitted. Files generated within this study are available at Zenodo (https://zenodo.org/record/2547384#.XIK0x-RYZ14). The genome annotation file from GenCode [22] as well as the candidate REMs from the GENEHANCER database are included in the STITCHIT repository at www.github.com/schulzlab/STITCHIT.

### Author’s contributions

AM developed the MDL based segmentation algorithm together with JV. FS conducted all computational experiments presented in this study, and extended and parallelized the implementation to serve all presented use-cases. FS was advised by JG and MHS. NB assisted FS and MHS with the validation analysis. MHS supervised and designed the study. FS and AM wrote the manuscript. All authors commented on and reviewed the manuscript.

## Acknowledgements

We thank Martin Vingron, Verena Heinrich, Anna Ramisch, and Tobias Zehnder from the Max Planck Institute for Molecular Genetics, Berlin, Germany for helpful comments and discussions, Markus List, TU Munich, for support with processing the DNase1-seq data and the ENCODE, Roadmap, and Blueprint consortia for providing their data. We also acknowledge the DEEP consortium for a critical discussion of the main idea of this manuscript.

## Additional Files

### Additional file 1 — Supplementary Material

Supplementary Section 1 contains details on data IDs and data processing. Supplementary Section 2 holds a more detailed description of the STITCHIT algorithm. In Section 3, details on the POE method for gene-expression discretization are provided. Details on related methods to link regulatory elements to genes are shown in Section 4. Additional Figures and Tables are listed in Supplementary Section 5.

### Additional file 2 — Supplementary Excel Sheet

The excel sheet contains information on the intersection between STITCHIT and the COSMIC database as well as STITCHIT and GWAS hits.

