## Supplementary Material for "Integrative analysis of epigenetics data identifies gene-specific regulatory elements"

### Supplementary Section 1 Data and Preprocessing

#### Accession numbers

Supplementary Table S1 lists the File IDs, Tissue/Cell type, Data type and Internal Sample ID of the raw data files obtained from ENCODE[1], Blueprint[2] and Roadmap [3]. Note that access to Blueprint and DEEP raw data files is restricted and requires an application to the Data Access Committee of the respective consortia.

| Consortium | File ID | Tissue/Cell type | Data type | Internal Sample ID |
| --- | --- | --- | --- | --- |
| ENCODE | [ENCFF000DUC](https://www.encodeproject.org/files/ENCFF000DUC/) | Endothelial cell of umbilical vein | RNA-seq | E_112ENC_124GGK_774AAA_719AAA_743IPG |
| ENCODE | [ENCFF000DUH](https://www.encodeproject.org/files/ENCFF000DUH/) | Endothelial cell of umbilical vein | RNA-seq |  |
| ENCODE | [ENCFF000DUD](https://www.encodeproject.org/files/ENCFF000DUD/) | Endothelial cell of umbilical vein | RNA-seq |  |
| ENCODE | [ENCFF000DUI](https://www.encodeproject.org/files/ENCFF000DUI/) | Endothelial cell of umbilical vein | RNA-seq |  |
| ENCODE | [ENCFF000DUE](https://www.encodeproject.org/files/ENCFF000DUE/) | Endothelial cell of umbilical vein | RNA-seq |  |
| ENCODE | [ENCFF000DUJ](https://www.encodeproject.org/files/ENCFF000DUJ/) | Endothelial cell of umbilical vein | RNA-seq |  |
| ENCODE | [ENCFF000DUF](https://www.encodeproject.org/files/ENCFF000DUF/) | Endothelial cell of umbilical vein | RNA-seq |  |
| ENCODE | [ENCFF000DUK](https://www.encodeproject.org/files/ENCFF000DUK/) | Endothelial cell of umbilical vein | RNA-seq |  |
| ENCODE | [ENCFF000DUG](https://www.encodeproject.org/files/ENCFF000DUG/) | Endothelial cell of umbilical vein | RNA-seq |  |
| ENCODE | [ENCFF000DUL](https://www.encodeproject.org/files/ENCFF000DUL/) | Endothelial cell of umbilical vein | RNA-seq |  |
| ENCODE | [ENCFF000IDF](https://www.encodeproject.org/files/ENCFF000IDF/) | keratinocyte | RNA-seq | E_589ENC_591ENC_818IBE_567ENC_565ENC_569ENC_564ENC_563ENC_586ENC |
| ENCODE | ENCFF000IDH | keratinocyte | RNA-seq |  |
| ENCODE | [ENCFF000IDC](https://www.encodeproject.org/files/ENCFF000IDC/) | keratinocyte | RNA-seq |  |
| ENCODE | [ENCFF000IDI](https://www.encodeproject.org/files/ENCFF000IDI/) | keratinocyte | RNA-seq |  |
| ENCODE | [ENCFF000HXI](https://www.encodeproject.org/files/ENCFF000HXI/) | keratinocyte | RNA-seq |  |
| ENCODE | [ENCFF000HXN](https://www.encodeproject.org/files/ENCFF000HXN/) | keratinocyte | RNA-seq |  |
| ENCODE | [ENCFF000HXH](https://www.encodeproject.org/files/ENCFF000HXH/) | keratinocyte | RNA-seq |  |
| ENCODE | [ENCFF000HZB](https://www.encodeproject.org/files/ENCFF000HZB/) | keratinocyte | RNA-seq |  |
| ENCODE | [ENCFF000HXG](https://www.encodeproject.org/files/ENCFF000HXG/) | keratinocyte | RNA-seq |  |
| ENCODE | [ENCFF000HZM](https://www.encodeproject.org/files/ENCFF000HZM/) | keratinocyte | RNA-seq |  |
| ENCODE | [ENCFF000EEA](https://www.encodeproject.org/files/ENCFF000EEA/) | keratinocyte | RNA-seq |  |
| ENCODE | [ENCFF000EEF](https://www.encodeproject.org/files/ENCFF000EEF/) | keratinocyte | RNA-seq |  |
| ENCODE | [ENCFF000EEB](https://www.encodeproject.org/files/ENCFF000EEB/) | keratinocyte | RNA-seq |  |
| ENCODE | [ENCFF000EEG](https://www.encodeproject.org/files/ENCFF000EEG/) | keratinocyte | RNA-seq |  |
| ENCODE | [ENCFF000EEC](https://www.encodeproject.org/files/ENCFF000EEC/) | keratinocyte | RNA-seq |  |
| ENCODE | [ENCFF000EEH](https://www.encodeproject.org/files/ENCFF000EEH/) | keratinocyte | RNA-seq |  |
| ENCODE | [ENCFF000EED](https://www.encodeproject.org/files/ENCFF000EED/) | keratinocyte | RNA-seq |  |
| ENCODE | [ENCFF000EEI](https://www.encodeproject.org/files/ENCFF000EEI/) | keratinocyte | RNA-seq |  |
| ENCODE | [ENCFF000EEE](https://www.encodeproject.org/files/ENCFF000EEE/) | keratinocyte | RNA-seq |  |
| ENCODE | [ENCFF000EEJ](https://www.encodeproject.org/files/ENCFF000EEJ/) | keratinocyte | RNA-seq |  |
| ENCODE | [ENCFF000ENZ](https://www.encodeproject.org/files/ENCFF000ENZ/) | fibroblast of lung | RNA-seq | E_004ENC_008ENC |
| ENCODE | [ENCFF000EOA](https://www.encodeproject.org/files/ENCFF000EOA/) | fibroblast of lung | RNA-seq |  |
| ENCODE | [ENCFF000ENY](https://www.encodeproject.org/files/ENCFF000ENY/) | fibroblast of lung | RNA-seq |  |
| ENCODE | [ENCFF000EON](https://www.encodeproject.org/files/ENCFF000EON/) | fibroblast of lung | RNA-seq |  |
| ENCODE | [ENCFF000IJZ](https://www.encodeproject.org/files/ENCFF000IJZ/) | fibroblast of lung | RNA-seq | E_340AAA_936GPP_612ENC_613ENC |
| ENCODE | [ENCFF000IKH](https://www.encodeproject.org/files/ENCFF000IKH/) | fibroblast of lung | RNA-seq |  |
| ENCODE | [ENCFF000IKA](https://www.encodeproject.org/files/ENCFF000IKA/) | fibroblast of lung | RNA-seq |  |
| ENCODE | [ENCFF000IKM](https://www.encodeproject.org/files/ENCFF000IKM/) | fibroblast of lung | RNA-seq |  |
| ENCODE | [ENCFF000HDD](https://www.encodeproject.org/files/ENCFF000HDD/) | IMR-90 | RNA-seq | E_460CLV_369ENC_372ENC_370ENC_373ENC_368ENC_371ENC |
| ENCODE | [ENCFF000HDF](https://www.encodeproject.org/files/ENCFF000HDF/) | IMR-89 | RNA-seq |  |
| ENCODE | [ENCFF000HDE](https://www.encodeproject.org/files/ENCFF000HDE/) | IMR-88 | RNA-seq |  |
| ENCODE | [ENCFF000HDH](https://www.encodeproject.org/files/ENCFF000HDH/) | IMR-87 | RNA-seq |  |
| ENCODE | [ENCFF000HAZ](https://www.encodeproject.org/files/ENCFF000HAZ/) | IMR-86 | RNA-seq |  |
| ENCODE | [ENCFF000HBG](https://www.encodeproject.org/files/ENCFF000HBG/) | IMR-85 | RNA-seq |  |
| ENCODE | [ENCFF000HBA](https://www.encodeproject.org/files/ENCFF000HBA/) | IMR-84 | RNA-seq |  |
| ENCODE | [ENCFF000HBI](https://www.encodeproject.org/files/ENCFF000HBI/) | IMR-83 | RNA-seq |  |
| ENCODE | [ENCFF000HCB](https://www.encodeproject.org/files/ENCFF000HCB/) | IMR-82 | RNA-seq |  |
| ENCODE | [ENCFF000HCC](https://www.encodeproject.org/files/ENCFF000HCC/) | IMR-81 | RNA-seq |  |
| ENCODE | [ENCFF000HCI](https://www.encodeproject.org/files/ENCFF000HCI/) | IMR-80 | RNA-seq |  |
| ENCODE | [ENCFF000HCJ](https://www.encodeproject.org/files/ENCFF000HCJ/) | IMR-79 | RNA-seq |  |
| ENCODE | [ENCFF000EQK](https://www.encodeproject.org/files/ENCFF000EQK/) | foreskin fibroblast | RNA-seq | E_074ENC_911DVL_753AAA_754AAA |
| ENCODE | [ENCFF000EQT](https://www.encodeproject.org/files/ENCFF000EQT/) | foreskin fibroblast | RNA-seq |  |
| ENCODE | [ENCFF000EQS](https://www.encodeproject.org/files/ENCFF000EQS/) | foreskin fibroblast | RNA-seq |  |
| ENCODE | [ENCFF000ESG](https://www.encodeproject.org/files/ENCFF000ESG/) | foreskin fibroblast | RNA-seq |  |
| ENCODE | [ENCFF000ESM](https://www.encodeproject.org/files/ENCFF000ESM/) | B cell | RNA-seq | E_484ENC |
| ENCODE | [ENCFF000ESS](https://www.encodeproject.org/files/ENCFF000ESS/) | B cell | RNA-seq |  |
| ENCODE | [ENCFF000ESN](https://www.encodeproject.org/files/ENCFF000ESN/) | B cell | RNA-seq |  |
| ENCODE | [ENCFF000EST](https://www.encodeproject.org/files/ENCFF000EST/) | B cell | RNA-seq |  |
| ENCODE | [ENCFF000ESO](https://www.encodeproject.org/files/ENCFF000ESO/) | B cell | RNA-seq |  |
| ENCODE | [ENCFF000ESU](https://www.encodeproject.org/files/ENCFF000ESU/) | B cell | RNA-seq |  |
| ENCODE | [ENCFF000ESP](https://www.encodeproject.org/files/ENCFF000ESP/) | B cell | RNA-seq | E_852WTL_483ENC |
| ENCODE | [ENCFF000ESV](https://www.encodeproject.org/files/ENCFF000ESV/) | B cell | RNA-seq |  |
| ENCODE | [ENCFF000ESQ](https://www.encodeproject.org/files/ENCFF000ESQ/) | B cell | RNA-seq |  |
| ENCODE | [ENCFF000ESW](https://www.encodeproject.org/files/ENCFF000ESW/) | B cell | RNA-seq |  |
| ENCODE | [ENCFF000ESR](https://www.encodeproject.org/files/ENCFF000ESR/) | B cell | RNA-seq |  |
| ENCODE | [ENCFF000ESY](https://www.encodeproject.org/files/ENCFF000ESY/) | B cell | RNA-seq |  |
| ENCODE | [ENCFF000HUX](https://www.encodeproject.org/files/ENCFF000HUX/) | CD14-positive monocyte | RNA-seq | E_477CHZ_628ENC_626ENC |
| ENCODE | [ENCFF000HVD](https://www.encodeproject.org/files/ENCFF000HVD/) | CD14-positive monocyte | RNA-seq |  |
| ENCODE | [ENCFF000HUY](https://www.encodeproject.org/files/ENCFF000HUY/) | CD14-positive monocyte | RNA-seq |  |
| ENCODE | [ENCFF000HVE](https://www.encodeproject.org/files/ENCFF000HVE/) | CD14-positive monocyte | RNA-seq |  |
| ENCODE | [ENCFF000HUZ](https://www.encodeproject.org/files/ENCFF000HUZ/) | CD14-positive monocyte | RNA-seq |  |
| ENCODE | [ENCFF000HVF](https://www.encodeproject.org/files/ENCFF000HVF/) | CD14-positive monocyte | RNA-seq |  |
| ENCODE | [ENCFF000HUU](https://www.encodeproject.org/files/ENCFF000HUU/) | CD14-positive monocyte | RNA-seq | E_629ENC_627ENC |
| ENCODE | [ENCFF000HVA](https://www.encodeproject.org/files/ENCFF000HVA/) | CD14-positive monocyte | RNA-seq |  |
| ENCODE | [ENCFF000HUV](https://www.encodeproject.org/files/ENCFF000HUV/) | CD14-positive monocyte | RNA-seq |  |
| ENCODE | [ENCFF000HVB](https://www.encodeproject.org/files/ENCFF000HVB/) | CD14-positive monocyte | RNA-seq |  |
| ENCODE | [ENCFF000HUW](https://www.encodeproject.org/files/ENCFF000HUW/) | CD14-positive monocyte | RNA-seq |  |
| ENCODE | [ENCFF000HVC](https://www.encodeproject.org/files/ENCFF000HVC/) | CD14-positive monocyte | RNA-seq |  |
| ENCODE | [ENCFF002DML](https://www.encodeproject.org/files/ENCFF002DML/) | skeletal muscle myoblast | RNA-seq | E_328AAA |
| ENCODE | [ENCFF002DMM](https://www.encodeproject.org/files/ENCFF002DMM/) | skeletal muscle myoblast | RNA-seq |  |
| ENCODE | [ENCFF002DMH](https://www.encodeproject.org/files/ENCFF002DMH/) | skeletal muscle myoblast | RNA-seq | E_665DRD |
| ENCODE | [ENCFF002DMI](https://www.encodeproject.org/files/ENCFF002DMI/) | skeletal muscle myoblast | RNA-seq |  |
| ENCODE | [ENCFF000EUS](https://www.encodeproject.org/files/ENCFF000EUS/) | Hematopoietic multipotent progenitor cell | RNA-seq | E_485ENC |
| ENCODE | [ENCFF000EUR](https://www.encodeproject.org/files/ENCFF000EUR/) | Hematopoietic multipotent progenitor cell | RNA-seq |  |
| ENCODE | [ENCFF002DMN](https://www.encodeproject.org/files/ENCFF002DMN/) | fibroblast of arm | RNA-seq | E_367AAA |
| ENCODE | [ENCFF002DMO](https://www.encodeproject.org/files/ENCFF002DMO/) | fibroblast of arm | RNA-seq |  |
| ENCODE | [ENCFF002DHP](https://www.encodeproject.org/files/ENCFF002DHP/) | fibroblast of arm | RNA-seq | E_372AAA |
| ENCODE | [ENCFF002DHQ](https://www.encodeproject.org/files/ENCFF002DHQ/) | fibroblast of arm | RNA-seq |  |
| ENCODE | [ENCFF424DEO](https://www.encodeproject.org/files/ENCFF424DEO/) | astrocyte | RNA-seq | E_021ENC |
| ENCODE | [ENCFF969BOA](https://www.encodeproject.org/files/ENCFF969BOA/) | astrocyte | RNA-seq |  |
| ENCODE | [ENCFF393GQZ](https://www.encodeproject.org/files/ENCFF393GQZ/) | astrocyte | RNA-seq | E_0052WQU |
| ENCODE | [ENCFF253MPW](https://www.encodeproject.org/files/ENCFF253MPW/) | astrocyte | RNA-seq |  |
| ENCODE | [ENCFF976PUJ](https://www.encodeproject.org/files/ENCFF976PUJ/) | adrenal gland | RNA-seq | E_371OZD_227VDO |
| ENCODE | [ENCFF576RAC](https://www.encodeproject.org/files/ENCFF576RAC/) | adrenal gland | RNA-seq |  |
| ENCODE | [ENCFF911BTP](https://www.encodeproject.org/files/ENCFF911BTP/) | adrenal gland | RNA-seq |  |
| ENCODE | [ENCFF904VHM](https://www.encodeproject.org/files/ENCFF904VHM/) | adrenal gland | RNA-seq |  |
| ENCODE | [ENCFF423RWS](https://www.encodeproject.org/files/ENCFF423RWS/) | adrenal gland | RNA-seq | E_423TBO_548ULT |
| ENCODE | [ENCFF145QCK](https://www.encodeproject.org/files/ENCFF145QCK/) | adrenal gland | RNA-seq |  |
| ENCODE | [ENCFF884HRS](https://www.encodeproject.org/files/ENCFF884HRS/) | adrenal gland | RNA-seq | E_942WBM_724UYV |
| ENCODE | [ENCFF095SRY](https://www.encodeproject.org/files/ENCFF095SRY/) | adrenal gland | RNA-seq |  |
| ENCODE | [ENCFF855MDA](https://www.encodeproject.org/files/ENCFF855MDA/) | adrenal gland | RNA-seq |  |
| ENCODE | [ENCFF223ZRA](https://www.encodeproject.org/files/ENCFF223ZRA/) | adrenal gland | RNA-seq |  |
| ENCODE | [ENCFF751LMQ](https://www.encodeproject.org/files/ENCFF751LMQ/) | thyroid gland | RNA-seq | E_658GLE_376ASB |
| ENCODE | [ENCFF188LMP](https://www.encodeproject.org/files/ENCFF188LMP/) | thyroid gland | RNA-seq |  |
| ENCODE | [ENCFF151GUG](https://www.encodeproject.org/files/ENCFF151GUG/) | thyroid gland | RNA-seq |  |
| ENCODE | [ENCFF628TMU](https://www.encodeproject.org/files/ENCFF628TMU/) | thyroid gland | RNA-seq |  |
| ENCODE | [ENCFF072VKD](https://www.encodeproject.org/files/ENCFF072VKD/) | thyroid gland | RNA-seq | E_624WYQ_702NUN |
| ENCODE | [ENCFF484BLA](https://www.encodeproject.org/files/ENCFF484BLA/) | thyroid gland | RNA-seq |  |
| ENCODE | [ENCFF086TFZ](https://www.encodeproject.org/files/ENCFF086TFZ/) | thyroid gland | RNA-seq |  |
| ENCODE | [ENCFF351OAS](https://www.encodeproject.org/files/ENCFF351OAS/) | thyroid gland | RNA-seq |  |
| ENCODE | [ENCFF411UIT](https://www.encodeproject.org/files/ENCFF411UIT/) | transverse colon | RNA-seq | E_174MRL_767MGS |
| ENCODE | [ENCFF992NAN](https://www.encodeproject.org/files/ENCFF992NAN/) | transverse colon | RNA-seq |  |
| ENCODE | [ENCFF411WXY](https://www.encodeproject.org/files/ENCFF411WXY/) | transverse colon | RNA-seq | E_409AIP_174MRL |
| ENCODE | [ENCFF543BVT](https://www.encodeproject.org/files/ENCFF543BVT/) | transverse colon | RNA-seq |  |
| ENCODE | [ENCFF122HNW](https://www.encodeproject.org/files/ENCFF122HNW/) | transverse colon | RNA-seq | E_767POE_974EYF |
| ENCODE | [ENCFF069KBE](https://www.encodeproject.org/files/ENCFF069KBE/) | transverse colon | RNA-seq |  |
| ENCODE | [ENCFF139SHO](https://www.encodeproject.org/files/ENCFF139SHO/) | gastrocnemius medialis | RNA-seq | E_921WWM_472LWI |
| ENCODE | [ENCFF825PCO](https://www.encodeproject.org/files/ENCFF825PCO/) | gastrocnemius medialis | RNA-seq |  |
| ENCODE | [ENCFF468EIW](https://www.encodeproject.org/files/ENCFF468EIW/) | uterus | RNA-seq | E_020UWI_887WTT |
| ENCODE | [ENCFF093MHL](https://www.encodeproject.org/files/ENCFF093MHL/) | uterus | RNA-seq |  |
| ENCODE | [ENCFF208XMO](https://www.encodeproject.org/files/ENCFF208XMO/) | uterus | RNA-seq | E_310EYM_210YNQ |
| ENCODE | [ENCFF141YVJ](https://www.encodeproject.org/files/ENCFF141YVJ/) | uterus | RNA-seq |  |
| ENCODE | [ENCFF017FUT](https://www.encodeproject.org/files/ENCFF017FUT/) | ovary | RNA-seq | E_279OPH_711CPB |
| ENCODE | [ENCFF856QRO](https://www.encodeproject.org/files/ENCFF856QRO/) | ovary | RNA-seq |  |
| ENCODE | [ENCFF140UYT](https://www.encodeproject.org/files/ENCFF140UYT/) | testis | RNA-seq | E_197JOA_315DHM |
| ENCODE | [ENCFF794EAB](https://www.encodeproject.org/files/ENCFF794EAB/) | testis | RNA-seq |  |
| ENCODE | [ENCFF959BUF](https://www.encodeproject.org/files/ENCFF959BUF/) | testis | RNA-seq |  |
| ENCODE | [ENCFF256AHC](https://www.encodeproject.org/files/ENCFF256AHC/) | testis | RNA-seq |  |
| ENCODE | [ENCFF714UWF](https://www.encodeproject.org/files/ENCFF714UWF/) | prostate gland | RNA-seq | E_524RBS_291PVB |
| ENCODE | [ENCFF638FRV](https://www.encodeproject.org/files/ENCFF638FRV/) | prostate gland | RNA-seq |  |
| ENCODE | [ENCFF644CJJ](https://www.encodeproject.org/files/ENCFF644CJJ/) | bipolar spindle neuron | RNA-seq | E_369AAA |
| ENCODE | [ENCFF774ZGB](https://www.encodeproject.org/files/ENCFF774ZGB/) | bipolar spindle neuron | RNA-seq |  |
| ENCODE | [ENCFF040TFC](https://www.encodeproject.org/files/ENCFF040TFC/) | bipolar spindle neuron | RNA-seq | E_374AAA |
| ENCODE | [ENCFF743MBB](https://www.encodeproject.org/files/ENCFF743MBB/) | bipolar spindle neuron | RNA-seq |  |
| ENCODE | [ENCFF245VTB](https://www.encodeproject.org/files/ENCFF245VTB/) | hepatocyte | RNA-seq | E_077RUJ |
| ENCODE | [ENCFF369QXD](https://www.encodeproject.org/files/ENCFF369QXD/) | hepatocyte | RNA-seq |  |
| ENCODE | [ENCFF653EZE](https://www.encodeproject.org/files/ENCFF653EZE/) | hepatocyte | RNA-seq | E_520VFV |
| ENCODE | [ENCFF644RGE](https://www.encodeproject.org/files/ENCFF644RGE/) | hepatocyte | RNA-seq |  |
| ENCODE | [ENCFF927MXX](https://www.encodeproject.org/files/ENCFF927MXX/) | myotube | RNA-seq | E_526EMC |
| ENCODE | [ENCFF011VTJ](https://www.encodeproject.org/files/ENCFF011VTJ/) | myotube | RNA-seq |  |
| ENCODE | [ENCFF999QZD](https://www.encodeproject.org/files/ENCFF999QZD/) | myotube | RNA-seq | E_236AFP_140MNV |
| ENCODE | [ENCFF023LZD](https://www.encodeproject.org/files/ENCFF023LZD/) | myotube | RNA-seq |  |
| ENCODE | [ENCFF119TIN](https://www.encodeproject.org/files/ENCFF119TIN/) | LHCN-M2 | RNA-seq | E_234AAA |
| ENCODE | [ENCFF494PBN](https://www.encodeproject.org/files/ENCFF494PBN/) | LHCN-M2 | RNA-seq |  |
| ENCODE | [ENCFF700EUG](https://www.encodeproject.org/files/ENCFF700EUG/) | LHCN-M2 | RNA-seq | E_869RCC_231OMQ |
| ENCODE | [ENCFF193NKV](https://www.encodeproject.org/files/ENCFF193NKV/) | LHCN-M2 | RNA-seq |  |
| ENCODE | [ENCFF939FVE](https://www.encodeproject.org/files/ENCFF939FVE/) | neural progenitor cell | RNA-seq | E_018TPT |
| ENCODE | [ENCFF201WLO](https://www.encodeproject.org/files/ENCFF201WLO/) | neural progenitor cell | RNA-seq |  |
| ENCODE | [ENCFF996KMK](https://www.encodeproject.org/files/ENCFF996KMK/) | neural progenitor cell | RNA-seq | E_044KWE |
| ENCODE | [ENCFF726PAY](https://www.encodeproject.org/files/ENCFF726PAY/) | neural progenitor cell | RNA-seq |  |
| ENCODE | [ENCFF000DGR](https://www.encodeproject.org/files/ENCFF000DGR/) | H1-hESC | RNA-seq | E_111ENC_780AAA_716AAA_051SJH_731AAA_734AAA_733AAA_732AAA |
| ENCODE | [ENCFF000DGZ](https://www.encodeproject.org/files/ENCFF000DGZ/) | H1-hESC | RNA-seq |  |
| ENCODE | [ENCFF000DGS](https://www.encodeproject.org/files/ENCFF000DGS/) | H1-hESC | RNA-seq |  |
| ENCODE | [ENCFF000DHA](https://www.encodeproject.org/files/ENCFF000DHA/) | H1-hESC | RNA-seq |  |
| ENCODE | [ENCFF000DGT](https://www.encodeproject.org/files/ENCFF000DGT/) | H1-hESC | RNA-seq |  |
| ENCODE | [ENCFF000DHB](https://www.encodeproject.org/files/ENCFF000DHB/) | H1-hESC | RNA-seq |  |
| ENCODE | [ENCFF000DGU](https://www.encodeproject.org/files/ENCFF000DGU/) | H1-hESC | RNA-seq |  |
| ENCODE | [ENCFF000DHC](https://www.encodeproject.org/files/ENCFF000DHC/) | H1-hESC | RNA-seq |  |
| ENCODE | [ENCFF000DGV](https://www.encodeproject.org/files/ENCFF000DGV/) | H1-hESC | RNA-seq |  |
| ENCODE | [ENCFF000DHD](https://www.encodeproject.org/files/ENCFF000DHD/) | H1-hESC | RNA-seq |  |
| ENCODE | [ENCFF000DGW](https://www.encodeproject.org/files/ENCFF000DGW/) | H1-hESC | RNA-seq |  |
| ENCODE | [ENCFF000DHE](https://www.encodeproject.org/files/ENCFF000DHE/) | H1-hESC | RNA-seq |  |
| ENCODE | [ENCFF000DGX](https://www.encodeproject.org/files/ENCFF000DGX/) | H1-hESC | RNA-seq |  |
| ENCODE | [ENCFF000DHF](https://www.encodeproject.org/files/ENCFF000DHF/) | H1-hESC | RNA-seq |  |
| ENCODE | [ENCFF000DGY](https://www.encodeproject.org/files/ENCFF000DGY/) | H1-hESC | RNA-seq |  |
| ENCODE | [ENCFF000DHG](https://www.encodeproject.org/files/ENCFF000DHG/) | H1-hESC | RNA-seq |  |
| ENCODE | [ENCFF000FEU](https://www.encodeproject.org/files/ENCFF000FEU/) | H1-hESC | RNA-seq |  |
| ENCODE | [ENCFF000FFF](https://www.encodeproject.org/files/ENCFF000FFF/) | H1-hESC | RNA-seq |  |
| ENCODE | [ENCFF000FET](https://www.encodeproject.org/files/ENCFF000FET/) | H1-hESC | RNA-seq |  |
| ENCODE | [ENCFF000FFG](https://www.encodeproject.org/files/ENCFF000FFG/) | H1-hESC | RNA-seq |  |
| ENCODE | [ENCFF000FJF](https://www.encodeproject.org/files/ENCFF000FJF/) | H1-hESC | RNA-seq |  |
| ENCODE | [ENCFF000FJG](https://www.encodeproject.org/files/ENCFF000FJG/) | H1-hESC | RNA-seq |  |
| ENCODE | [ENCFF000FHG](https://www.encodeproject.org/files/ENCFF000FHG/) | H1-hESC | RNA-seq |  |
| ENCODE | [ENCFF000FHH](https://www.encodeproject.org/files/ENCFF000FHH/) | H1-hESC | RNA-seq |  |
| ENCODE | [ENCFF002DMP](https://www.encodeproject.org/files/ENCFF002DMP/) | H7-hESC | RNA-seq | E_293AAA |
| ENCODE | [ENCFF002DMQ](https://www.encodeproject.org/files/ENCFF002DMQ/) | H7-hESC | RNA-seq |  |
| ENCODE | [ENCFF002DMJ](https://www.encodeproject.org/files/ENCFF002DMJ/) | H7-hESC | RNA-seq | E_297CQV_291AAA_624XJG |
| ENCODE | [ENCFF002DMK](https://www.encodeproject.org/files/ENCFF002DMK/) | H7-hESC | RNA-seq |  |
| ENCODE | [ENCFF001DOL](https://www.encodeproject.org/files/ENCFF001DOL/) | Endothelial cell of umbilical vein | DNase1-seq | E_112ENC_124GGK_774AAA_719AAA_743IPG |
| ENCODE | [ENCFF000STF](https://www.encodeproject.org/files/ENCFF000STF/) | Endothelial cell of umbilical vein | DNase1-seq |  |
| ENCODE | [ENCFF000STI](https://www.encodeproject.org/files/ENCFF000STI/) | Endothelial cell of umbilical vein | DNase1-seq |  |
| ENCODE | [ENCFF001ECW](https://www.encodeproject.org/files/ENCFF001ECW/) | keratinocyte | DNase1-seq | E_589ENC_591ENC_818IBE_567ENC_565ENC_569ENC_564ENC_563ENC_586ENC |
| ENCODE | [ENCFF001ECX](https://www.encodeproject.org/files/ENCFF001ECX/) | keratinocyte | DNase1-seq |  |
| ENCODE | [ENCFF000TBT](https://www.encodeproject.org/files/ENCFF000TBT/) | keratinocyte | DNase1-seq |  |
| ENCODE | [ENCFF000TBS](https://www.encodeproject.org/files/ENCFF000TBS/) | keratinocyte | DNase1-seq |  |
| ENCODE | [ENCFF001CMS](https://www.encodeproject.org/files/ENCFF001CMS/) | fibroblast of lung | DNase1-seq | E_004ENC_008ENC |
| ENCODE | [ENCFF001CMR](https://www.encodeproject.org/files/ENCFF001CMR/) | fibroblast of lung | DNase1-seq |  |
| ENCODE | [ENCFF001EDJ](https://www.encodeproject.org/files/ENCFF001EDJ/) | fibroblast of lung | DNase1-seq | E_340AAA_936GPP_612ENC_613ENC |
| ENCODE | [ENCFF001COU](https://www.encodeproject.org/files/ENCFF001COU/) | foreskin fibroblast | DNase1-seq | E_074ENC_911DVL_753AAA_754AAA |
| ENCODE | [ENCFF348QHI](https://www.encodeproject.org/files/ENCFF348QHI/) | foreskin fibroblast | DNase1-seq |  |
| ENCODE | [ENCFF480CMA](https://www.encodeproject.org/files/ENCFF480CMA/) | foreskin fibroblast | DNase1-seq |  |
| ENCODE | [ENCFF393PQT](https://www.encodeproject.org/files/ENCFF393PQT/) | foreskin fibroblast | DNase1-seq |  |
| ENCODE | [ENCFF001CQG](https://www.encodeproject.org/files/ENCFF001CQG/) | B cell | DNase1-seq | E_852WTL_483ENC |
| ENCODE | [ENCFF001CQF](https://www.encodeproject.org/files/ENCFF001CQF/) | B cell | DNase1-seq |  |
| ENCODE | [ENCFF001DZG](https://www.encodeproject.org/files/ENCFF001DZG/) | CD14-positive monocyte | DNase1-seq | E_477CHZ_628ENC_626ENC |
| ENCODE | [ENCFF001DZH](https://www.encodeproject.org/files/ENCFF001DZH/) | CD14-positive monocyte | DNase1-seq |  |
| ENCODE | [ENCFF000TBL](https://www.encodeproject.org/files/ENCFF000TBL/) | CD14-positive monocyte | DNase1-seq |  |
| ENCODE | [ENCFF875UUR](https://www.encodeproject.org/files/ENCFF875UUR/) | skeletal muscle myoblast | DNase1-seq | E_328AAA |
| ENCODE | [ENCFF465ATZ](https://www.encodeproject.org/files/ENCFF465ATZ/) | skeletal muscle myoblast | DNase1-seq |  |
| ENCODE | [ENCFF689CAW](https://www.encodeproject.org/files/ENCFF689CAW/) | skeletal muscle myoblast | DNase1-seq | E_665DRD |
| ENCODE | [ENCFF218GIV](https://www.encodeproject.org/files/ENCFF218GIV/) | skeletal muscle myoblast | DNase1-seq |  |
| ENCODE | [ENCFF001CQN](https://www.encodeproject.org/files/ENCFF001CQN/) | Hematopoietic multipotent progenitor cell | DNase1-seq | E_485ENC |
| ENCODE | [ENCFF001CQO](https://www.encodeproject.org/files/ENCFF001CQO/) | Hematopoietic multipotent progenitor cell | DNase1-seq |  |
| ENCODE | [ENCFF001CQP](https://www.encodeproject.org/files/ENCFF001CQP/) | Hematopoietic multipotent progenitor cell | DNase1-seq |  |
| ENCODE | [ENCFF405CPB](https://www.encodeproject.org/files/ENCFF405CPB/) | fibroblast of arm | DNase1-seq | E_367AAA |
| ENCODE | [ENCFF846EWZ](https://www.encodeproject.org/files/ENCFF846EWZ/) | fibroblast of arm | DNase1-seq |  |
| ENCODE | [ENCFF521XIS](https://www.encodeproject.org/files/ENCFF521XIS/) | fibroblast of arm | DNase1-seq |  |
| ENCODE | [ENCFF494CZA](https://www.encodeproject.org/files/ENCFF494CZA/) | fibroblast of arm | DNase1-seq |  |
| ENCODE | [ENCFF618XNK](https://www.encodeproject.org/files/ENCFF618XNK/) | fibroblast of arm | DNase1-seq |  |
| ENCODE | [ENCFF673BIU](https://www.encodeproject.org/files/ENCFF673BIU/) | fibroblast of arm | DNase1-seq |  |
| ENCODE | [ENCFF313TQJ](https://www.encodeproject.org/files/ENCFF313TQJ/) | fibroblast of arm | DNase1-seq | E_372AAA |
| ENCODE | [ENCFF413FVI](https://www.encodeproject.org/files/ENCFF413FVI/) | fibroblast of arm | DNase1-seq |  |
| ENCODE | [ENCFF400MSN](https://www.encodeproject.org/files/ENCFF400MSN/) | fibroblast of arm | DNase1-seq |  |
| ENCODE | [ENCFF349AHE](https://www.encodeproject.org/files/ENCFF349AHE/) | fibroblast of arm | DNase1-seq |  |
| ENCODE | [ENCFF001CEM](https://www.encodeproject.org/files/ENCFF001CEM/) | astrocyte | DNase1-seq | E_021ENC |
| ENCODE | [ENCFF001EBK](https://www.encodeproject.org/files/ENCFF001EBK/) | astrocyte | DNase1-seq | E_0052WQU |
| ENCODE | [ENCFF031MXQ](https://www.encodeproject.org/files/ENCFF031MXQ/) | adrenal gland | DNase1-seq | E_371OZD_227VDO |
| ENCODE | [ENCFF195BAO](https://www.encodeproject.org/files/ENCFF195BAO/) | adrenal gland | DNase1-seq |  |
| ENCODE | [ENCFF963BRG](https://www.encodeproject.org/files/ENCFF963BRG/) | adrenal gland | DNase1-seq |  |
| ENCODE | [ENCFF994JJX](https://www.encodeproject.org/files/ENCFF994JJX/) | adrenal gland | DNase1-seq |  |
| ENCODE | [ENCFF036OXC](https://www.encodeproject.org/files/ENCFF036OXC/) | adrenal gland | DNase1-seq | E_423TBO_548ULT |
| ENCODE | [ENCFF411QZU](https://www.encodeproject.org/files/ENCFF411QZU/) | adrenal gland | DNase1-seq |  |
| ENCODE | [ENCFF629RYZ](https://www.encodeproject.org/files/ENCFF629RYZ/) | adrenal gland | DNase1-seq |  |
| ENCODE | [ENCFF232YBS](https://www.encodeproject.org/files/ENCFF232YBS/) | adrenal gland | DNase1-seq |  |
| ENCODE | [ENCFF391QPN](https://www.encodeproject.org/files/ENCFF391QPN/) | adrenal gland | DNase1-seq |  |
| ENCODE | [ENCFF244XPY](https://www.encodeproject.org/files/ENCFF244XPY/) | adrenal gland | DNase1-seq |  |
| ENCODE | [ENCFF504HQG](https://www.encodeproject.org/files/ENCFF504HQG/) | adrenal gland | DNase1-seq |  |
| ENCODE | [ENCFF466GBQ](https://www.encodeproject.org/files/ENCFF466GBQ/) | adrenal gland | DNase1-seq |  |
| ENCODE | [ENCFF119STK](https://www.encodeproject.org/files/ENCFF119STK/) | adrenal gland | DNase1-seq | E_942WBM_724UYV |
| ENCODE | [ENCFF950OGZ](https://www.encodeproject.org/files/ENCFF950OGZ/) | adrenal gland | DNase1-seq |  |
| ENCODE | [ENCFF787LTC](https://www.encodeproject.org/files/ENCFF787LTC/) | adrenal gland | DNase1-seq |  |
| ENCODE | [ENCFF460GRP](https://www.encodeproject.org/files/ENCFF460GRP/) | adrenal gland | DNase1-seq |  |
| ENCODE | [ENCFF224AJP](https://www.encodeproject.org/files/ENCFF224AJP/) | thyroid gland | DNase1-seq | E_658GLE_376ASB |
| ENCODE | [ENCFF868WDJ](https://www.encodeproject.org/files/ENCFF868WDJ/) | thyroid gland | DNase1-seq |  |
| ENCODE | [ENCFF431RGV](https://www.encodeproject.org/files/ENCFF431RGV/) | thyroid gland | DNase1-seq |  |
| ENCODE | [ENCFF289FGW](https://www.encodeproject.org/files/ENCFF289FGW/) | thyroid gland | DNase1-seq |  |
| ENCODE | [ENCFF428IKC](https://www.encodeproject.org/files/ENCFF428IKC/) | thyroid gland | DNase1-seq |  |
| ENCODE | [ENCFF439OJT](https://www.encodeproject.org/files/ENCFF439OJT/) | thyroid gland | DNase1-seq |  |
| ENCODE | [ENCFF761ZSC](https://www.encodeproject.org/files/ENCFF761ZSC/) | thyroid gland | DNase1-seq |  |
| ENCODE | [ENCFF482QJN](https://www.encodeproject.org/files/ENCFF482QJN/) | thyroid gland | DNase1-seq |  |
| ENCODE | [ENCFF089TZR](https://www.encodeproject.org/files/ENCFF089TZR/) | thyroid gland | DNase1-seq | E_624WYQ_702NUN |
| ENCODE | [ENCFF961TSD](https://www.encodeproject.org/files/ENCFF961TSD/) | thyroid gland | DNase1-seq |  |
| ENCODE | [ENCFF142IUW](https://www.encodeproject.org/files/ENCFF142IUW/) | thyroid gland | DNase1-seq |  |
| ENCODE | [ENCFF475RYH](https://www.encodeproject.org/files/ENCFF475RYH/) | thyroid gland | DNase1-seq |  |
| ENCODE | [ENCFF679WCJ](https://www.encodeproject.org/files/ENCFF679WCJ/) | thyroid gland | DNase1-seq |  |
| ENCODE | [ENCFF264RJN](https://www.encodeproject.org/files/ENCFF264RJN/) | thyroid gland | DNase1-seq |  |
| ENCODE | [ENCFF882HXM](https://www.encodeproject.org/files/ENCFF882HXM/) | thyroid gland | DNase1-seq |  |
| ENCODE | [ENCFF332VYY](https://www.encodeproject.org/files/ENCFF332VYY/) | thyroid gland | DNase1-seq |  |
| ENCODE | [ENCFF476DQM](https://www.encodeproject.org/files/ENCFF476DQM/) | thyroid gland | DNase1-seq |  |
| ENCODE | [ENCFF680IKR](https://www.encodeproject.org/files/ENCFF680IKR/) | thyroid gland | DNase1-seq |  |
| ENCODE | [ENCFF721IWB](https://www.encodeproject.org/files/ENCFF721IWB/) | transverse colon | DNase1-seq | E_174MRL_767MGS |
| ENCODE | [ENCFF005PIM](https://www.encodeproject.org/files/ENCFF005PIM/) | transverse colon | DNase1-seq |  |
| ENCODE | [ENCFF745TGS](https://www.encodeproject.org/files/ENCFF745TGS/) | transverse colon | DNase1-seq |  |
| ENCODE | [ENCFF341DNU](https://www.encodeproject.org/files/ENCFF341DNU/) | transverse colon | DNase1-seq |  |
| ENCODE | [ENCFF523STC](https://www.encodeproject.org/files/ENCFF523STC/) | transverse colon | DNase1-seq |  |
| ENCODE | [ENCFF392JVD](https://www.encodeproject.org/files/ENCFF392JVD/) | transverse colon | DNase1-seq |  |
| ENCODE | [ENCFF017KXG](https://www.encodeproject.org/files/ENCFF017KXG/) | transverse colon | DNase1-seq | E_409AIP_668MJW |
| ENCODE | [ENCFF981ADW](https://www.encodeproject.org/files/ENCFF981ADW/) | transverse colon | DNase1-seq |  |
| ENCODE | [ENCFF907JCI](https://www.encodeproject.org/files/ENCFF907JCI/) | transverse colon | DNase1-seq |  |
| ENCODE | [ENCFF212DTF](https://www.encodeproject.org/files/ENCFF212DTF/) | transverse colon | DNase1-seq |  |
| ENCODE | [ENCFF600OSX](https://www.encodeproject.org/files/ENCFF600OSX/) | transverse colon | DNase1-seq |  |
| ENCODE | [ENCFF753LNI](https://www.encodeproject.org/files/ENCFF753LNI/) | transverse colon | DNase1-seq |  |
| ENCODE | [ENCFF123JGV](https://www.encodeproject.org/files/ENCFF123JGV/) | transverse colon | DNase1-seq | E_767POE_974EYF |
| ENCODE | [ENCFF228LAZ](https://www.encodeproject.org/files/ENCFF228LAZ/) | transverse colon | DNase1-seq |  |
| ENCODE | [ENCFF789LVT](https://www.encodeproject.org/files/ENCFF789LVT/) | transverse colon | DNase1-seq |  |
| ENCODE | [ENCFF163OOS](https://www.encodeproject.org/files/ENCFF163OOS/) | transverse colon | DNase1-seq |  |
| ENCODE | [ENCFF803BOW](https://www.encodeproject.org/files/ENCFF803BOW/) | transverse colon | DNase1-seq |  |
| ENCODE | [ENCFF181HWO](https://www.encodeproject.org/files/ENCFF181HWO/) | transverse colon | DNase1-seq |  |
| ENCODE | [ENCFF830AJL](https://www.encodeproject.org/files/ENCFF830AJL/) | transverse colon | DNase1-seq |  |
| ENCODE | [ENCFF750DCS](https://www.encodeproject.org/files/ENCFF750DCS/) | transverse colon | DNase1-seq |  |
| ENCODE | [ENCFF402LYH](https://www.encodeproject.org/files/ENCFF402LYH/) | gastrocnemius medialis | DNase1-seq | E_921WWM_472LWI |
| ENCODE | [ENCFF310HZA](https://www.encodeproject.org/files/ENCFF310HZA/) | gastrocnemius medialis | DNase1-seq |  |
| ENCODE | [ENCFF511KTN](https://www.encodeproject.org/files/ENCFF511KTN/) | gastrocnemius medialis | DNase1-seq |  |
| ENCODE | [ENCFF430QUU](https://www.encodeproject.org/files/ENCFF430QUU/) | gastrocnemius medialis | DNase1-seq |  |
| ENCODE | [ENCFF435DRI](https://www.encodeproject.org/files/ENCFF435DRI/) | gastrocnemius medialis | DNase1-seq |  |
| ENCODE | [ENCFF549DQK](https://www.encodeproject.org/files/ENCFF549DQK/) | gastrocnemius medialis | DNase1-seq |  |
| ENCODE | [ENCFF536GTU](https://www.encodeproject.org/files/ENCFF536GTU/) | gastrocnemius medialis | DNase1-seq |  |
| ENCODE | [ENCFF644GNP](https://www.encodeproject.org/files/ENCFF644GNP/) | gastrocnemius medialis | DNase1-seq |  |
| ENCODE | [ENCFF147VEZ](https://www.encodeproject.org/files/ENCFF147VEZ/) | uterus | DNase1-seq | E_020UWI_887WTT |
| ENCODE | [ENCFF215CCC](https://www.encodeproject.org/files/ENCFF215CCC/) | uterus | DNase1-seq |  |
| ENCODE | [ENCFF426IQZ](https://www.encodeproject.org/files/ENCFF426IQZ/) | uterus | DNase1-seq |  |
| ENCODE | [ENCFF903BLH](https://www.encodeproject.org/files/ENCFF903BLH/) | uterus | DNase1-seq |  |
| ENCODE | [ENCFF503KIS](https://www.encodeproject.org/files/ENCFF503KIS/) | uterus | DNase1-seq |  |
| ENCODE | [ENCFF793TLT](https://www.encodeproject.org/files/ENCFF793TLT/) | uterus | DNase1-seq |  |
| ENCODE | [ENCFF026UVD](https://www.encodeproject.org/files/ENCFF026UVD/) | uterus | DNase1-seq | E_310EYM_210YNQ |
| ENCODE | [ENCFF972HMB](https://www.encodeproject.org/files/ENCFF972HMB/) | uterus | DNase1-seq |  |
| ENCODE | [ENCFF489OVA](https://www.encodeproject.org/files/ENCFF489OVA/) | uterus | DNase1-seq |  |
| ENCODE | [ENCFF281WBK](https://www.encodeproject.org/files/ENCFF281WBK/) | uterus | DNase1-seq |  |
| ENCODE | [ENCFF454GEF](https://www.encodeproject.org/files/ENCFF454GEF/) | uterus | DNase1-seq |  |
| ENCODE | [ENCFF663JLO](https://www.encodeproject.org/files/ENCFF663JLO/) | uterus | DNase1-seq |  |
| ENCODE | [ENCFF315GDY](https://www.encodeproject.org/files/ENCFF315GDY/) | ovary | DNase1-seq | E_279OPH_711CPB |
| ENCODE | [ENCFF593IJZ](https://www.encodeproject.org/files/ENCFF593IJZ/) | ovary | DNase1-seq |  |
| ENCODE | [ENCFF339MTK](https://www.encodeproject.org/files/ENCFF339MTK/) | ovary | DNase1-seq |  |
| ENCODE | [ENCFF648NXJ](https://www.encodeproject.org/files/ENCFF648NXJ/) | ovary | DNase1-seq |  |
| ENCODE | [ENCFF906SAW](https://www.encodeproject.org/files/ENCFF906SAW/) | ovary | DNase1-seq |  |
| ENCODE | [ENCFF649SYN](https://www.encodeproject.org/files/ENCFF649SYN/) | ovary | DNase1-seq |  |
| ENCODE | [ENCFF373PSV](https://www.encodeproject.org/files/ENCFF373PSV/) | testis | DNase1-seq | E_197JOA_315DHM |
| ENCODE | [ENCFF033QYM](https://www.encodeproject.org/files/ENCFF033QYM/) | testis | DNase1-seq |  |
| ENCODE | [ENCFF716WKO](https://www.encodeproject.org/files/ENCFF716WKO/) | testis | DNase1-seq |  |
| ENCODE | [ENCFF359CJV](https://www.encodeproject.org/files/ENCFF359CJV/) | testis | DNase1-seq |  |
| ENCODE | [ENCFF503BZM](https://www.encodeproject.org/files/ENCFF503BZM/) | testis | DNase1-seq |  |
| ENCODE | [ENCFF848NYW](https://www.encodeproject.org/files/ENCFF848NYW/) | testis | DNase1-seq |  |
| ENCODE | [ENCFF790FQW](https://www.encodeproject.org/files/ENCFF790FQW/) | testis | DNase1-seq |  |
| ENCODE | [ENCFF655XTX](https://www.encodeproject.org/files/ENCFF655XTX/) | testis | DNase1-seq |  |
| ENCODE | [ENCFF003PTJ](https://www.encodeproject.org/files/ENCFF003PTJ/) | prostate gland | DNase1-seq | E_524RBS_291PVB |
| ENCODE | [ENCFF673WML](https://www.encodeproject.org/files/ENCFF673WML/) | prostate gland | DNase1-seq |  |
| ENCODE | [ENCFF677FUH](https://www.encodeproject.org/files/ENCFF677FUH/) | prostate gland | DNase1-seq |  |
| ENCODE | [ENCFF271HAG](https://www.encodeproject.org/files/ENCFF271HAG/) | prostate gland | DNase1-seq |  |
| ENCODE | [ENCFF984PAZ](https://www.encodeproject.org/files/ENCFF984PAZ/) | prostate gland | DNase1-seq |  |
| ENCODE | [ENCFF577XUT](https://www.encodeproject.org/files/ENCFF577XUT/) | prostate gland | DNase1-seq |  |
| ENCODE | [ENCFF637XDH](https://www.encodeproject.org/files/ENCFF637XDH/) | prostate gland | DNase1-seq |  |
| ENCODE | [ENCFF953LAY](https://www.encodeproject.org/files/ENCFF953LAY/) | prostate gland | DNase1-seq |  |
| ENCODE | [ENCFF185JDV](https://www.encodeproject.org/files/ENCFF185JDV/) | bipolar spindle neuron | DNase1-seq | E_369AAA |
| ENCODE | [ENCFF856VKF](https://www.encodeproject.org/files/ENCFF856VKF/) | bipolar spindle neuron | DNase1-seq |  |
| ENCODE | [ENCFF335WYY](https://www.encodeproject.org/files/ENCFF335WYY/) | bipolar spindle neuron | DNase1-seq |  |
| ENCODE | [ENCFF404BAG](https://www.encodeproject.org/files/ENCFF404BAG/) | bipolar spindle neuron | DNase1-seq |  |
| ENCODE | [ENCFF634HDQ](https://www.encodeproject.org/files/ENCFF634HDQ/) | bipolar spindle neuron | DNase1-seq |  |
| ENCODE | [ENCFF436PPF](https://www.encodeproject.org/files/ENCFF436PPF/) | bipolar spindle neuron | DNase1-seq |  |
| ENCODE | [ENCFF447HQG](https://www.encodeproject.org/files/ENCFF447HQG/) | bipolar spindle neuron | DNase1-seq | E_374AAA |
| ENCODE | [ENCFF347BGI](https://www.encodeproject.org/files/ENCFF347BGI/) | bipolar spindle neuron | DNase1-seq |  |
| ENCODE | [ENCFF674SOV](https://www.encodeproject.org/files/ENCFF674SOV/) | bipolar spindle neuron | DNase1-seq |  |
| ENCODE | [ENCFF441FWE](https://www.encodeproject.org/files/ENCFF441FWE/) | bipolar spindle neuron | DNase1-seq |  |
| ENCODE | [ENCFF977EUX](https://www.encodeproject.org/files/ENCFF977EUX/) | bipolar spindle neuron | DNase1-seq |  |
| ENCODE | [ENCFF665YOB](https://www.encodeproject.org/files/ENCFF665YOB/) | bipolar spindle neuron | DNase1-seq |  |
| ENCODE | [ENCFF287MXM](https://www.encodeproject.org/files/ENCFF287MXM/) | hepatocyte | DNase1-seq | E_077RUJ |
| ENCODE | [ENCFF526GHY](https://www.encodeproject.org/files/ENCFF526GHY/) | hepatocyte | DNase1-seq |  |
| ENCODE | [ENCFF415CLL](https://www.encodeproject.org/files/ENCFF415CLL/) | hepatocyte | DNase1-seq |  |
| ENCODE | [ENCFF459ZWQ](https://www.encodeproject.org/files/ENCFF459ZWQ/) | hepatocyte | DNase1-seq |  |
| ENCODE | [ENCFF754VPL](https://www.encodeproject.org/files/ENCFF754VPL/) | hepatocyte | DNase1-seq |  |
| ENCODE | [ENCFF699RMR](https://www.encodeproject.org/files/ENCFF699RMR/) | hepatocyte | DNase1-seq |  |
| ENCODE | [ENCFF364MZF](https://www.encodeproject.org/files/ENCFF364MZF/) | hepatocyte | DNase1-seq | E_520VFV |
| ENCODE | [ENCFF335LPO](https://www.encodeproject.org/files/ENCFF335LPO/) | hepatocyte | DNase1-seq |  |
| ENCODE | [ENCFF001DOC](https://www.encodeproject.org/files/ENCFF001DOC/) | myotube | DNase1-seq | E_526EMC |
| ENCODE | [ENCFF001DNO](https://www.encodeproject.org/files/ENCFF001DNO/) | myotube | DNase1-seq | E_236AFP_140MNV |
| ENCODE | [ENCFF001DNN](https://www.encodeproject.org/files/ENCFF001DNN/) | myotube | DNase1-seq |  |
| ENCODE | [ENCFF001DNP](https://www.encodeproject.org/files/ENCFF001DNP/) | myotube | DNase1-seq |  |
| ENCODE | [ENCFF186RIJ](https://www.encodeproject.org/files/ENCFF186RIJ/) | LHCN-M2 | DNase1-seq | E_234AAA |
| ENCODE | [ENCFF675VWC](https://www.encodeproject.org/files/ENCFF675VWC/) | LHCN-M2 | DNase1-seq |  |
| ENCODE | [ENCFF797RJZ](https://www.encodeproject.org/files/ENCFF797RJZ/) | LHCN-M2 | DNase1-seq |  |
| ENCODE | [ENCFF249NQM](https://www.encodeproject.org/files/ENCFF249NQM/) | LHCN-M2 | DNase1-seq | E_869RCC_231OMQ |
| ENCODE | [ENCFF717GPW](https://www.encodeproject.org/files/ENCFF717GPW/) | LHCN-M2 | DNase1-seq |  |
| ENCODE | [ENCFF822UVF](https://www.encodeproject.org/files/ENCFF822UVF/) | LHCN-M2 | DNase1-seq |  |
| ENCODE | [ENCFF219BIC](https://www.encodeproject.org/files/ENCFF219BIC/) | neural progenitor cell | DNase1-seq | E_018TPT |
| ENCODE | [ENCFF344WBT](https://www.encodeproject.org/files/ENCFF344WBT/) | neural progenitor cell | DNase1-seq |  |
| ENCODE | [ENCFF484OQF](https://www.encodeproject.org/files/ENCFF484OQF/) | neural progenitor cell | DNase1-seq |  |
| ENCODE | [ENCFF412NKG](https://www.encodeproject.org/files/ENCFF412NKG/) | neural progenitor cell | DNase1-seq |  |
| ENCODE | [ENCFF198NDG](https://www.encodeproject.org/files/ENCFF198NDG/) | neural progenitor cell | DNase1-seq | E_044KWE |
| ENCODE | [ENCFF690YCI](https://www.encodeproject.org/files/ENCFF690YCI/) | neural progenitor cell | DNase1-seq |  |
| ENCODE | [ENCFF226OBW](https://www.encodeproject.org/files/ENCFF226OBW/) | neural progenitor cell | DNase1-seq |  |
| ENCODE | [ENCFF569MQH](https://www.encodeproject.org/files/ENCFF569MQH/) | neural progenitor cell | DNase1-seq |  |
| ENCODE | [ENCFF795AUA](https://www.encodeproject.org/files/ENCFF795AUA/) | neural progenitor cell | DNase1-seq |  |
| ENCODE | [ENCFF996JWF](https://www.encodeproject.org/files/ENCFF996JWF/) | neural progenitor cell | DNase1-seq |  |
| ENCODE | [ENCFF814WXD](https://www.encodeproject.org/files/ENCFF814WXD/) | neural progenitor cell | DNase1-seq |  |
| ENCODE | [ENCFF853IEJ](https://www.encodeproject.org/files/ENCFF853IEJ/) | neural progenitor cell | DNase1-seq |  |
| ENCODE | [ENCFF000SOF](https://www.encodeproject.org/files/ENCFF000SOF/) | H1-hESC | DNase1-seq | E_111ENC_780AAA_716AAA_051SJH_731AAA_734AAA_733AAA_732AAA |
| ENCODE | [ENCFF000SOH](https://www.encodeproject.org/files/ENCFF000SOH/) | H1-hESC | DNase1-seq |  |
| ENCODE | [ENCFF001CVK](https://www.encodeproject.org/files/ENCFF001CVK/) | H1-hESC | DNase1-seq |  |
| ENCODE | [ENCFF001BAJ](https://www.encodeproject.org/files/ENCFF001BAJ/) | H7-hESC | DNase1-seq | E_293AAA |
| ENCODE | [ENCFF001CXB](https://www.encodeproject.org/files/ENCFF001CXB/) | H7-hESC | DNase1-seq | E_297CQV_291AAA_624XJG |
| Blueprint | 130919_SN546_0217_B_C2GFPACXX_CTTGTA_1_read2.fastq.gz | CD8-positive, alpha-beta T cell | RNA-seq | B_C0066P12 |
| Blueprint | 130919_SN546_0217_B_C2GFPACXX_CTTGTA_1_read1.fastq.gz | CD8-positive, alpha-beta T cell | RNA-seq | B_C0066P12 |
| Blueprint | 130207_SN546_0194_A_H093AADXX_CTTGTA_1_read1.fastq.gz | CD14-positive, CD16-negative classical monocyte | RNA-seq | B_C005PS12 |
| Blueprint | 130207_SN546_0194_A_H093AADXX_CTTGTA_1_read2.fastq.gz | CD14-positive, CD16-negative classical monocyte | RNA-seq | B_C005PS12 |
| Blueprint | 150226_SN935_0006_B_C5YBTACXX_TTAGGC_4_read1.fastq.gz | Acute Lymphocytic Leukemia | RNA-seq | B_S00DFM11 |
| Blueprint | 150226_SN935_0006_B_C5YBTACXX_TTAGGC_4_read2.fastq.gz | Acute Lymphocytic Leukemia | RNA-seq | B_S00DFM11 |
| Blueprint | C3JYVACXX_lane6_5385_ACAGTG_L006_R1.fastq.gz | macrophage - T=6days LPS | RNA-seq | B_S00HSH11 |
| Blueprint | C3JYVACXX_lane6_5389_ATCACG_L006_R1.fastq.gz | macrophage - T=6days LPS | RNA-seq | B_S00JRB11 |
| Blueprint | C2MVTACXX_lane7_4641_ACAGTG_L007_R1.fastq.gz | macrophage - T=6days LPS | RNA-seq | B_S00BYT11 |
| Blueprint | C2NTEACXX_lane7_4819_ACAGTG_L007_R1.fastq.gz | macrophage - T=6days LPS | RNA-seq | B_S00CS011 |
| Blueprint | C250RACXX_lane5_3990_CTTGTA_L005_R2.fastq.gz | CD34-negative, CD41-positive, CD42-positive megakaryocyte cell | RNA-seq | B_C006NSB1 |
| Blueprint | C250RACXX_lane5_3990_CTTGTA_L005_R1.fastq.gz | CD34-negative, CD41-positive, CD42-positive megakaryocyte cell | RNA-seq | B_C006NSB1 |
| Blueprint | C250RACXX_lane7_3990_CTTGTA_L007_R2.fastq.gz | CD34-negative, CD41-positive, CD42-positive megakaryocyte cell | RNA-seq | B_C006NSB1 |
| Blueprint | C250RACXX_lane7_3990_CTTGTA_L007_R1.fastq.gz | CD34-negative, CD41-positive, CD42-positive megakaryocyte cell | RNA-seq | B_C006NSB1 |
| Blueprint | C250RACXX_lane8_3990_CTTGTA_L008_R1.fastq.gz | CD34-negative, CD41-positive, CD42-positive megakaryocyte cell | RNA-seq | B_C006NSB1 |
| Blueprint | C250RACXX_lane8_3990_CTTGTA_L008_R2.fastq.gz | CD34-negative, CD41-positive, CD42-positive megakaryocyte cell | RNA-seq | B_C006NSB1 |
| Blueprint | C250RACXX_lane6_3990_CTTGTA_L006_R2.fastq.gz | CD34-negative, CD41-positive, CD42-positive megakaryocyte cell | RNA-seq | B_C006NSB1 |
| Blueprint | C250RACXX_lane6_3990_CTTGTA_L006_R1.fastq.gz | CD34-negative, CD41-positive, CD42-positive megakaryocyte cell | RNA-seq | B_C006NSB1 |
| Blueprint | 130919_SN546_0217_B_C2GFPACXX_ACAGTG_2_read1.fastq.gz | CD34-negative, CD41-positive, CD42-positive megakaryocyte cell | RNA-seq | B_S004BT |
| Blueprint | 130919_SN546_0217_B_C2GFPACXX_ACAGTG_2_read2.fastq.gz | CD34-negative, CD41-positive, CD42-positive megakaryocyte cell | RNA-seq | B_S004BT |
| Blueprint | 140618_SN935_0195_A_C42K0ACXX_ACTGAT_3_read1.fastq.gz | CD4-positive, alpha-beta T cell | RNA-seq | B_S008H111 |
| Blueprint | 140618_SN935_0195_A_C42K0ACXX_ACTGAT_3_read2.fastq.gz | CD4-positive, alpha-beta T cell | RNA-seq | B_S008H111 |
| Blueprint | 140618_SN935_0195_A_C42K0ACXX_GCCAAT_1_read1.fastq.gz | erythroblast | RNA-seq | B_S002R512 |
| Blueprint | 140618_SN935_0195_A_C42K0ACXX_GCCAAT_1_read2.fastq.gz | erythroblast | RNA-seq | B_S002R512 |
| Blueprint | 130919_SN546_0217_B_C2GFPACXX_GTGAAA_2_read2.fastq.gz | erythroblast | RNA-seq | B_S002S312 |
| Blueprint | 130919_SN546_0217_B_C2GFPACXX_GTGAAA_2_read1.fastq.gz | erythroblast | RNA-seq | B_S002S312 |
| Blueprint | 130919_SN546_0217_B_C2GFPACXX_TTAGGC_5_read1.fastq.gz | macrophage | RNA-seq | B_S001S714 |
| Blueprint | 130919_SN546_0217_B_C2GFPACXX_TTAGGC_5_read2.fastq.gz | macrophage | RNA-seq | B_S001S714 |
| Blueprint | 130919_SN546_0217_B_C2GFPACXX_ATCACG_4_read2.fastq.gz | inflammatory macrophage | RNA-seq | B_S001MJ12 |
| Blueprint | 130919_SN546_0217_B_C2GFPACXX_ATCACG_4_read1.fastq.gz | inflammatory macrophage | RNA-seq | B_S001MJ12 |
| Blueprint | 130919_SN546_0217_B_C2GFPACXX_TGACCA_6_read1.fastq.gz | inflammatory macrophage | RNA-seq | B_S0022I14 |
| Blueprint | 130919_SN546_0217_B_C2GFPACXX_TGACCA_6_read2.fastq.gz | inflammatory macrophage | RNA-seq | B_S0022I14 |
| Blueprint | C3JYVACXX_lane6_5384_TGACCA_L006_R1.fastq.gz | macrophage - T=6days untreated | RNA-seq | B_S00HRJ11 |
| Blueprint | C2MVTACXX_lane7_4640_TGACCA_L007_R1.fastq.gz | macrophage - T=6days untreated | RNA-seq | B_S00BXV11 |
| Blueprint | C2NTEACXX_lane7_4818_TGACCA_L007_R1.fastq.gz | macrophage - T=6days untreated | RNA-seq | B_S00CR211 |
| Blueprint | C3JYVACXX_lane6_5388_CTTGTA_L006_R1.fastq.gz | macrophage - T=6days untreated | RNA-seq | B_S00JQD11 |
| Blueprint | 150429_SN935_0015_A_C5RHEACXX_GATCAG_1_read2.fastq.gz | Acute Myeloid Leukemia | RNA-seq | B_S013M311 |
| Blueprint | 150429_SN935_0015_A_C5RHEACXX_GATCAG_1_read1.fastq.gz | Acute Myeloid Leukemia | RNA-seq | B_S013M311 |
| Blueprint | 150429_SN935_0015_A_C5RHEACXX_TTAGGC_1_read1.fastq.gz | Acute Myeloid Leukemia | RNA-seq | B_S00D0F11 |
| Blueprint | 150429_SN935_0015_A_C5RHEACXX_TTAGGC_1_read2.fastq.gz | Acute Myeloid Leukemia | RNA-seq | B_S00D0F11 |
| Blueprint | 151209_SN546_0285_A_H3WLNADXX_ACAGTG_1_read1.fastq.gz | Acute Myeloid Leukemia | RNA-seq | B_S00D6311 |
| Blueprint | 151209_SN546_0285_A_H3WLNADXX_ACAGTG_1_read2.fastq.gz | Acute Myeloid Leukemia | RNA-seq | B_S00D6311 |
| Blueprint | 150429_SN935_0015_A_C5RHEACXX_GTCCGC_5_read2.fastq.gz | Acute Myeloid Leukemia | RNA-seq | B_S005EJ11 |
| Blueprint | 150429_SN935_0015_A_C5RHEACXX_GTCCGC_5_read1.fastq.gz | Acute Myeloid Leukemia | RNA-seq | B_S005EJ11 |
| Blueprint | 151209_SN546_0285_A_H3WLNADXX_GTGAAA_1_read1.fastq.gz | Acute Myeloid Leukemia | RNA-seq | B_S013QW11 |
| Blueprint | 151209_SN546_0285_A_H3WLNADXX_GTGAAA_1_read2.fastq.gz | Acute Myeloid Leukemia | RNA-seq | B_S013QW11 |
| Blueprint | 150226_SN935_0006_B_C5YBTACXX_ACTGAT_3_read2.fastq.gz | Acute Myeloid Leukemia | RNA-seq | B_S005FH11 |
| Blueprint | 150226_SN935_0006_B_C5YBTACXX_ACTGAT_3_read1.fastq.gz | Acute Myeloid Leukemia | RNA-seq | B_S005FH11 |
| Blueprint | 150429_SN935_0015_A_C5RHEACXX_TTAGGC_7_read2.fastq.gz | Acute Myeloid Leukemia | RNA-seq | B_S00XXH11 |
| Blueprint | 150429_SN935_0015_A_C5RHEACXX_TTAGGC_7_read1.fastq.gz | Acute Myeloid Leukemia | RNA-seq | B_S00XXH11 |
| Blueprint | 150226_SN935_0006_B_C5YBTACXX_GCCAAT_1_read1.fastq.gz | Acute Myeloid Leukemia | RNA-seq | B_S00CXR11 |
| Blueprint | 150226_SN935_0006_B_C5YBTACXX_GCCAAT_1_read2.fastq.gz | Acute Myeloid Leukemia | RNA-seq | B_S00CXR11 |
| Blueprint | 150603_SN935_0017_A_C73GHACXX_GGCTAC_1_read1.fastq.gz | Acute Myeloid Leukemia | RNA-seq | B_S013N111 |
| Blueprint | 150603_SN935_0017_A_C73GHACXX_GGCTAC_1_read2.fastq.gz | Acute Myeloid Leukemia | RNA-seq | B_S013N111 |
| Blueprint | 150226_SN935_0006_B_C5YBTACXX_GTCCGC_3_read2.fastq.gz | Acute Myeloid Leukemia | RNA-seq | B_S00CYP11 |
| Blueprint | 150226_SN935_0006_B_C5YBTACXX_GTCCGC_3_read1.fastq.gz | Acute Myeloid Leukemia | RNA-seq | B_S00CYP11 |
| Blueprint | 150429_SN935_0015_A_C5RHEACXX_CGATGT_4_read2.fastq.gz | Acute Myeloid Leukemia | RNA-seq | B_S00D5511 |
| Blueprint | 150429_SN935_0015_A_C5RHEACXX_CGATGT_4_read1.fastq.gz | Acute Myeloid Leukemia | RNA-seq | B_S00D5511 |
| Blueprint | 150226_SN935_0006_B_C5YBTACXX_ACAGTG_2_read2.fastq.gz | Acute Myeloid Leukemia | RNA-seq | B_S00D3911 |
| Blueprint | 150226_SN935_0006_B_C5YBTACXX_ACAGTG_2_read1.fastq.gz | Acute Myeloid Leukemia | RNA-seq | B_S00D3911 |
| Blueprint | 150429_SN935_0015_A_C5RHEACXX_TGACCA_8_read2.fastq.gz | Acute Myeloid Leukemia | RNA-seq | B_S00XUN11 |
| Blueprint | 150429_SN935_0015_A_C5RHEACXX_TGACCA_8_read1.fastq.gz | Acute Myeloid Leukemia | RNA-seq | B_S00XUN11 |
| Blueprint | 150429_SN935_0015_A_C5RHEACXX_ACTTGA_2_read2.fastq.gz | Acute Myeloid Leukemia | RNA-seq | B_S00XYF11 |
| Blueprint | 150429_SN935_0015_A_C5RHEACXX_ACTTGA_2_read1.fastq.gz | Acute Myeloid Leukemia | RNA-seq | B_S00XYF11 |
| Blueprint | 150429_SN935_0015_A_C5RHEACXX_CAGATC_3_read2.fastq.gz | Acute Myeloid Leukemia | RNA-seq | B_S013PY11 |
| Blueprint | 150429_SN935_0015_A_C5RHEACXX_CAGATC_3_read1.fastq.gz | Acute Myeloid Leukemia | RNA-seq | B_S013PY11 |
| Blueprint | 150429_SN935_0015_A_C5RHEACXX_GATCAG_7_read1.fastq.gz | Acute Myeloid Leukemia | RNA-seq | B_S00Y1311 |
| Blueprint | 150429_SN935_0015_A_C5RHEACXX_GATCAG_7_read2.fastq.gz | Acute Myeloid Leukemia | RNA-seq | B_S00Y1311 |
| Blueprint | 150429_SN935_0015_A_C5RHEACXX_TGACCA_2_read2.fastq.gz | Acute Myeloid Leukemia | RNA-seq | B_S00XWJ11 |
| Blueprint | 150429_SN935_0015_A_C5RHEACXX_TGACCA_2_read1.fastq.gz | Acute Myeloid Leukemia | RNA-seq | B_S00XWJ11 |
| Blueprint | 150603_SN935_0017_A_C73GHACXX_CAGATC_1_read1.fastq.gz | Acute Myeloid Leukemia | RNA-seq | B_S013RU11 |
| Blueprint | 150603_SN935_0017_A_C73GHACXX_CAGATC_1_read2.fastq.gz | Acute Myeloid Leukemia | RNA-seq | B_S013RU11 |
| Blueprint | 150226_SN935_0006_B_C5YBTACXX_GTGAAA_2_read2.fastq.gz | Acute Myeloid Leukemia | RNA-seq | B_S00D1D11 |
| Blueprint | 150226_SN935_0006_B_C5YBTACXX_GTGAAA_2_read1.fastq.gz | Acute Myeloid Leukemia | RNA-seq | B_S00D1D11 |
| Blueprint | 150429_SN935_0015_A_C5RHEACXX_GGCTAC_3_read2.fastq.gz | Acute Myeloid Leukemia | RNA-seq | B_S00Y0511 |
| Blueprint | 150429_SN935_0015_A_C5RHEACXX_GGCTAC_3_read1.fastq.gz | Acute Myeloid Leukemia | RNA-seq | B_S00Y0511 |
| Blueprint | 150429_SN935_0015_A_C5RHEACXX_ACTGAT_5_read2.fastq.gz | Acute Myeloid Leukemia | RNA-seq | B_S00D4711 |
| Blueprint | 150429_SN935_0015_A_C5RHEACXX_ACTGAT_5_read1.fastq.gz | Acute Myeloid Leukemia | RNA-seq | B_S00D4711 |
| Blueprint | 150603_SN935_0017_A_C73GHACXX_CGATGT_2_read1.fastq.gz | Acute Myeloid Leukemia | RNA-seq | B_S00XVL11 |
| Blueprint | 150603_SN935_0017_A_C73GHACXX_CGATGT_2_read2.fastq.gz | Acute Myeloid Leukemia | RNA-seq | B_S00XVL11 |
| Blueprint | 150429_SN935_0015_A_C5RHEACXX_GTGAAA_4_read2.fastq.gz | Acute Myeloid Leukemia | RNA-seq | B_S013SS11 |
| Blueprint | 150429_SN935_0015_A_C5RHEACXX_GTGAAA_4_read1.fastq.gz | Acute Myeloid Leukemia | RNA-seq | B_S013SS11 |
| Blueprint | 150429_SN935_0015_A_C5RHEACXX_AGTCAA_6_read1.fastq.gz | Acute Myeloid Leukemia | RNA-seq | B_00Y6U11 |
| Blueprint | 150429_SN935_0015_A_C5RHEACXX_AGTCAA_6_read2.fastq.gz | Acute Myeloid Leukemia | RNA-seq | B_00Y6U11 |
| Blueprint | 150226_SN935_0006_B_C5YBTACXX_CTTGTA_1_read1.fastq.gz | Acute Myeloid Leukemia | RNA-seq | B_00CWT11 |
| Blueprint | 150226_SN935_0006_B_C5YBTACXX_CTTGTA_1_read2.fastq.gz | Acute Myeloid Leukemia | RNA-seq | B_00CWT11 |
| Blueprint | 150429_SN935_0015_A_C5RHEACXX_ACAGTG_6_read2.fastq.gz | Acute Myeloid Leukemia | RNA-seq | B_00Y4Y11 |
| Blueprint | 150429_SN935_0015_A_C5RHEACXX_ACAGTG_6_read1.fastq.gz | Acute Myeloid Leukemia | RNA-seq | B_00Y4Y11 |
| Blueprint | 130919_SN546_0217_B_C2GFPACXX_GATCAG_5_read2.fastq.gz | macrophage | RNA-seq | B_0022I12 |
| Blueprint | 130919_SN546_0217_B_C2GFPACXX_GATCAG_5_read1.fastq.gz | macrophage | RNA-seq | B_S0022I12 |
| Blueprint | 130919_SN546_0217_B_C2GFPACXX_ACTGAT_3_read1.fastq.gz | macrophage | RNA-seq | B_C005VG11 |
| Blueprint | 130919_SN546_0217_B_C2GFPACXX_ACTGAT_3_read2.fastq.gz | macrophage | RNA-seq | B_C005VG11 |
| Blueprint | 150917_SN546_0282_B_C7C6TACXX_GTCCGC_8_read1.fastq.gz | Chronic Lymphocytic Leukemia | RNA-seq | B_S00B0N11 |
| Blueprint | 150917_SN546_0282_B_C7C6TACXX_GTCCGC_8_read2.fastq.gz | Chronic Lymphocytic Leukemia | RNA-seq | B_S00B0N11 |
| Blueprint | 150603_SN935_0017_A_C73GHACXX_GCCAAT_8_read2.fastq.gz | Chronic Lymphocytic Leukemia | RNA-seq | B_S00B2J11 |
| Blueprint | 150603_SN935_0017_A_C73GHACXX_GCCAAT_8_read1.fastq.gz | Chronic Lymphocytic Leukemia | RNA-seq | B_S00B2J11 |
| Blueprint | C2MVTACXX_lane7_4642_GCCAAT_L007_R1.fastq.gz | macrophage - T=6days B-glucan | RNA-seq | B_S00C0J11 |
| Blueprint | C3JYVACXX_lane6_5386_GCCAAT_L006_R1.fastq.gz | macrophage - T=6days B-glucan | RNA-seq | B_S00HTF11 |
| Blueprint | C3JYVACXX_lane6_5390_TTAGGC_L006_R1.fastq.gz | macrophage - T=6days B-glucan | RNA-seq | B_S00JS911 |
| Blueprint | C2NTEACXX_lane7_4820_GCCAAT_L007_R1.fastq.gz | macrophage - T=6days B-glucan | RNA-seq | B_S00CTZ11 |
| Blueprint | s_120830_6.ACAGTG.read1.fastq.gz | CD14-positive, CD16-negative classical monocyte | RNA-seq | B_C0010KB1 |
| Blueprint | s_120830_6.ACAGTG.read2.fastq.gz | CD14-positive, CD16-negative classical monocyte | RNA-seq | B_C0010KB1 |
| Blueprint | s_120830_3.TTAGGC.read1.fastq.gz | CD14-positive, CD16-negative classical monocyte | RNA-seq | B_C001UYB4 |
| Blueprint | s_120830_3.TTAGGC.read2.fastq.gz | CD14-positive, CD16-negative classical monocyte | RNA-seq | B_C001UYB4 |
| Blueprint | s_120830_5.AGTTCC.read1.fastq.gz | CD14-positive, CD16-negative classical monocyte | RNA-seq | B_C0011IB1 |
| Blueprint | s_120830_5.AGTTCC.read2.fastq.gz | CD14-positive, CD16-negative classical monocyte | RNA-seq | B_C0011IB1 |
| Blueprint | C20FWACXX_lane5_4172_ACAGTG_L005_R1.fastq.gz | CD8-positive, alpha-beta T cell | DNase1-seq | B_C0066P12 |
| Blueprint | C20BGACXX_lane6_3758_CAGATC_L006_R1.fastq.gz | CD14-positive, CD16-negative classical monocyte | DNase1-seq | B_C005PS12 |
| Blueprint | C2U81ACXX_lane8_4901_CGATGT_L008_R1.fastq.gz | Acute Lymphocytic Leukemia | DNase1-seq | B_S00DFM11 |
| Blueprint | C3JYVACXX_lane8_5397_ACAGTG_L008_R1.fastq.gz | macrophage - T=6days LPS | DNase1-seq | B_S00HSH11 |
| Blueprint | C3JYVACXX_lane7_5401_ACTTGA_L007_R1.fastq.gz | macrophage - T=6days LPS | DNase1-seq | B_S00JRB11 |
| Blueprint | C2MVTACXX_lane8_4644_TGACCA_L008_R1.fastq.gz | macrophage - T=6days LPS | DNase1-seq | B_S00BYT11 |
| Blueprint | C2N9FACXX_lane8_4644_TGACCA_L008_R1.fastq.gz | macrophage - T=6days LPS | DNase1-seq | B_S00BYT11 |
| Blueprint | C2NE2ACXX_lane8_4754_GATCAG_L008_R1.fastq.gz | macrophage - T=6days LPS | DNase1-seq | B_S00CS011 |
| Blueprint | C2NTEACXX_lane8_4754_GATCAG_L008_R1.fastq.gz | macrophage - T=6days LPS | DNase1-seq | B_S00CS011 |
| Blueprint | C20BGACXX_lane5_3685_GCCAAT_L005_R1.fastq.gz | CD34-negative, CD41-positive, CD42-positive megakaryocyte cell | DNase1-seq | B_C006NSB1 |
| Blueprint | C5TLLACXX_lane7_6736_GGCTAC_L007_R1.fastq.gz | CD34-negative, CD41-positive, CD42-positive megakaryocyte cell | DNase1-seq | B_S004BT |
| Blueprint | C5TLLACXX_lane7_6737_AGTCAA_L007_R1.fastq.gz | CD4-positive, alpha-beta T cell | DNase1-seq | B_S008H111 |
| Blueprint | C7KUGACXX_lane6_7444_CAGATC_L006_R1.fastq.gz | erythroblast | DNase1-seq | B_S002R512 |
| Blueprint | C6LR9ACXX_lane8_7443_GCCAAT_L008_R1.fastq.gz | erythroblast | DNase1-seq | B_S002S312 |
| Blueprint | C2NTUACXX_lane8_4646_GCCAAT_L008_R1.fastq.gz | macrophage | DNase1-seq | B_S001S714 |
| Blueprint | C20BGACXX_lane5_3687_CTTGTA_L005_R1.fastq.gz | inflammatory macrophage | DNase1-seq | B_S001MJ12 |
| Blueprint | C2NTUACXX_lane8_4648_CTTGTA_L008_R1.fastq.gz | inflammatory macrophage | DNase1-seq | B_S0022I14 |
| Blueprint | C3JYVACXX_lane7_5396_TGACCA_L007_R1.fastq.gz | macrophage - T=6days untreated | DNase1-seq | B_S00HRJ11 |
| Blueprint | C2N9FACXX_lane8_4643_CGATGT_L008_R1.fastq.gz | macrophage - T=6days untreated | DNase1-seq | B_S00BXV11 |
| Blueprint | C2MVTACXX_lane8_4643_CGATGT_L008_R1.fastq.gz | macrophage - T=6days untreated | DNase1-seq | B_S00BXV11 |
| Blueprint | C2NTEACXX_lane8_4753_ACTTGA_L008_R1.fastq.gz | macrophage - T=6days untreated | DNase1-seq | B_S00CR211 |
| Blueprint | C2NE2ACXX_lane8_4753_ACTTGA_L008_R1.fastq.gz | macrophage - T=6days untreated | DNase1-seq | B_S00CR211 |
| Blueprint | C3JYVACXX_lane7_5400_TTAGGC_L007_R1.fastq.gz | macrophage - T=6days untreated | DNase1-seq | B_S00JQD11 |
| Blueprint | C791JACXX_lane1_7991_AGTCAA_L001_R1.fastq.gz | Acute Myeloid Leukemia | DNase1-seq | B_S013M311 |
| Blueprint | C4Y81ACXX_lane7_5974_TAGCTT_L007_R1.fastq.gz | Acute Myeloid Leukemia | DNase1-seq | B_S00D0F11 |
| Blueprint | C5682ACXX_lane7_5965_TGACCA_L007_R1.fastq.gz | Acute Myeloid Leukemia | DNase1-seq | B_S00D6311 |
| Blueprint | C20FWACXX_lane3_4175_CTTGTA_L003_R1.fastq.gz | Acute Myeloid Leukemia | DNase1-seq | B_S005EJ11 |
| Blueprint | C7KUGACXX_lane3_7994_CCGTCC_L003_R1.fastq.gz | Acute Myeloid Leukemia | DNase1-seq | B_S013QW11 |
| Blueprint | C20BGACXX_lane6_3754_ACAGTG_L006_R1.fastq.gz | Acute Myeloid Leukemia | DNase1-seq | B_S005FH11 |
| Blueprint | C68R7ACXX_lane5_6919_TTAGGC_L005_R1.fastq.gz | Acute Myeloid Leukemia | DNase1-seq | B_S00XXH11 |
| Blueprint | C55VCACXX_lane8_5975_GGCTAC_L008_R1.fastq.gz | Acute Myeloid Leukemia | DNase1-seq | B_S00CXR11 |
| Blueprint | C791JACXX_lane1_7992_AGTTCC_L001_R1.fastq.gz | Acute Myeloid Leukemia | DNase1-seq | B_S013N111 |
| Blueprint | C5682ACXX_lane7_5971_TTAGGC_L007_R1.fastq.gz | Acute Myeloid Leukemia | DNase1-seq | B_S00CYP11 |
| Blueprint | C4Y81ACXX_lane8_5970_ATCACG_L008_R1.fastq.gz | Acute Myeloid Leukemia | DNase1-seq | B_S00D5511 |
| Blueprint | C4Y81ACXX_lane8_5972_ACTTGA_L008_R1.fastq.gz | Acute Myeloid Leukemia | DNase1-seq | B_S00D3911 |
| Blueprint | C68R7ACXX_lane5_6913_GAGTGG_L005_R1.fastq.gz | Acute Myeloid Leukemia | DNase1-seq | B_S00XUN11 |
| Blueprint | C6957ACXX_lane6_6920_ACTTGA_L006_R1.fastq.gz | Acute Myeloid Leukemia | DNase1-seq | B_S00XYF11 |
| Blueprint | C7KUGACXX_lane3_7993_ATGTCA_L003_R1.fastq.gz | Acute Myeloid Leukemia | DNase1-seq | B_S013PY11 |
| Blueprint | C68R7ACXX_lane8_6922_TAGCTT_L008_R1.fastq.gz | Acute Myeloid Leukemia | DNase1-seq | B_S00Y1311 |
| Blueprint | C68R7ACXX_lane8_6923_GGCTAC_L008_R1.fastq.gz | Acute Myeloid Leukemia | DNase1-seq | B_S00Y1311 |
| Blueprint | C6957ACXX_lane5_6917_ATCACG_L005_R1.fastq.gz | Acute Myeloid Leukemia | DNase1-seq | B_S00XWJ11 |
| Blueprint | C7DJ1ACXX_lane2_7995_GTAGAG_L002_R1.fastq.gz | Acute Myeloid Leukemia | DNase1-seq | B_S013RU11 |
| Blueprint | C5682ACXX_lane7_5966_ACAGTG_L007_R1.fastq.gz | Acute Myeloid Leukemia | DNase1-seq | B_S00D1D11 |
| Blueprint | C68R7ACXX_lane8_6921_GATCAG_L008_R1.fastq.gz | Acute Myeloid Leukemia | DNase1-seq | B_S00Y0511 |
| Blueprint | C5682ACXX_lane8_5968_CAGATC_L008_R1.fastq.gz | Acute Myeloid Leukemia | DNase1-seq | B_S00D4711 |
| Blueprint | C6957ACXX_lane3_6914_GGTAGC_L003_R1.fastq.gz | Acute Myeloid Leukemia | DNase1-seq | B_S00XVL11 |
| Blueprint | C7KUGACXX_lane3_7996_GTCCGC_L003_R1.fastq.gz | Acute Myeloid Leukemia | DNase1-seq | B_S013SS11 |
| Blueprint | C6957ACXX_lane7_6926_ACAGTG_L007_R1.fastq.gz | Acute Myeloid Leukemia | DNase1-seq | B_S00Y6U11 |
| Blueprint | C4Y81ACXX_lane7_5967_GCCAAT_L007_R1.fastq.gz | Acute Myeloid Leukemia | DNase1-seq | B_S00CWT11 |
| Blueprint | C6957ACXX_lane6_6924_CGATGT_L006_R1.fastq.gz | Acute Myeloid Leukemia | DNase1-seq | B_S00Y4Y11 |
| Blueprint | C2NTUACXX_lane8_4647_CAGATC_L008_R1.fastq.gz | macrophage | DNase1-seq | B_S0022I12 |
| Blueprint | D1YHUACXX_lane8_3684_ACAGTG_L008_R1.fastq.gz | macrophage | DNase1-seq | B_C005VG11 |
| Blueprint | C68R7ACXX_lane5_7046_GCCAAT_L005_R1.fastq.gz | Chronic Lymphocytic Leukemia | DNase1-seq | B_S00B0N11 |
| Blueprint | C5NY1ACXX_lane8_6126_ATCACG_L008_R1.fastq.gz | Chronic Lymphocytic Leukemia | DNase1-seq | B_S00B2J11 |
| Blueprint | C2MVTACXX_lane8_4645_ACAGTG_L008_R1.fastq.gz | macrophage - T=6days B-glucan | DNase1-seq | B_S00C0J11 |
| Blueprint | C3JYVACXX_lane7_5398_GCCAAT_L007_R1.fastq.gz | macrophage - T=6days B-glucan | DNase1-seq | B_S00HTF11 |
| Blueprint | C3JYVACXX_lane7_5402_GATCAG_L007_R1.fastq.gz | macrophage - T=6days B-glucan | DNase1-seq | B_S00JS911 |
| Blueprint | C2NE2ACXX_lane8_4755_TAGCTT_L008_R1.fastq.gz | macrophage - T=6days B-glucan | DNase1-seq | B_S00CTZ11 |
| Blueprint | D1FB1ACXX_lane8_3324_ACAGTG_L008_R1.fastq.gz | CD14-positive, CD16-negative classical monocyte | DNase1-seq | B_C0010KB1 |
| Blueprint | D1FB1ACXX_lane6_3326_CAGATC_L006_R1.fastq.gz | CD14-positive, CD16-negative classical monocyte | DNase1-seq | B_C001UYB4 |
| Blueprint | D1FB1ACXX_lane8_3323_TGACCA_L008_R1.fastq.gz | CD14-positive, CD16-negative classical monocyte | DNase1-seq | B_C0011IB1 |
| Roadmap | [ENCFF583FQV](https://www.encodeproject.org/files/ENCFF583FQV/) | skin fibroblast | RNA-seq | [R_ENCSR637GBV](https://www.encodeproject.org/experiments/ENCSR637GBV/) |
| Roadmap | [ENCFF249BIO](https://www.encodeproject.org/files/ENCFF249BIO/) | skin fibroblast | RNA-seq | [R_ENCSR655XQF](https://www.encodeproject.org/experiments/ENCSR655XQF/) |
| Roadmap | [ENCFF946MIY](https://www.encodeproject.org/files/ENCFF946MIY/) | skin fibroblast | RNA-seq | R_ENCSR022MON |
| Roadmap | [ENCFF533GEZ](https://www.encodeproject.org/files/ENCFF533GEZ/) | skin fibroblast | RNA-seq | R_ENCSR022MON |
| Roadmap | [ENCFF273FCE](https://www.encodeproject.org/files/ENCFF273FCE/) | skin fibroblast | RNA-seq | [R_ENCSR982VYI](https://www.encodeproject.org/experiments/ENCSR982VYI/) |
| Roadmap | [ENCFF224MVX](https://www.encodeproject.org/files/ENCFF224MVX/) | fibroblast of skin of abdomen | RNA-seq | [R_ENCSR361DRG](https://www.encodeproject.org/experiments/ENCSR361DRG/) |
| Roadmap | [ENCFF079EAZ](https://www.encodeproject.org/files/ENCFF079EAZ/) | fibroblast of skin of abdomen | RNA-seq | [R_ENCSR681ALA](https://www.encodeproject.org/experiments/ENCSR681ALA/) |
| Roadmap | [ENCFF105OLM](https://www.encodeproject.org/files/ENCFF105OLM/) | IMR-90 | RNA-seq | R_ENCSR424FAZ |
| Roadmap | [ENCFF381KYM](https://www.encodeproject.org/files/ENCFF381KYM/) | IMR-90 | RNA-seq | R_ENCSR424FAZ |
| Roadmap | [ENCFF547LGQ](https://www.encodeproject.org/files/ENCFF547LGQ/) | IMR-90 | RNA-seq | R_ENCSR424FAZ |
| Roadmap | [ENCFF094HAH](https://www.encodeproject.org/files/ENCFF094HAH/) | IMR-90 | RNA-seq | R_ENCSR424FAZ |
| Roadmap | [ENCFF714LDG](https://www.encodeproject.org/files/ENCFF714LDG/) | trophoblast cell | RNA-seq | R_ENCSR762CJN |
| Roadmap | [ENCFF629SLX](https://www.encodeproject.org/files/ENCFF629SLX/) | muscle of arm | RNA-seq | [R_ENCSR406YML](https://www.encodeproject.org/experiments/ENCSR406YML/) |
| Roadmap | [ENCFF319ZJP](https://www.encodeproject.org/files/ENCFF319ZJP/) | muscle of arm | RNA-seq | R_ENCSR364IBB |
| Roadmap | [ENCFF070JPZ](https://www.encodeproject.org/files/ENCFF070JPZ/) | muscle of arm | RNA-seq | R_ENCSR364IBB |
| Roadmap | [ENCFF269DXG](https://www.encodeproject.org/files/ENCFF269DXG/) | muscle of arm | RNA-seq | R_ENCSR317LMH |
| Roadmap | [ENCFF442QQD](https://www.encodeproject.org/files/ENCFF442QQD/) | muscle of arm | RNA-seq | R_ENCSR620ZNQ |
| Roadmap | [ENCFF520PYM](https://www.encodeproject.org/files/ENCFF520PYM/) | muscle of arm | RNA-seq | R_ENCSR305NXN |
| Roadmap | [ENCFF617CUF](https://www.encodeproject.org/files/ENCFF617CUF/) | muscle of arm | RNA-seq | R_ENCSR677MYO |
| Roadmap | [ENCFF691YZM](https://www.encodeproject.org/files/ENCFF691YZM/) | muscle of arm | RNA-seq | R_ENCSR990LHE |
| Roadmap | [ENCFF713VFN](https://www.encodeproject.org/files/ENCFF713VFN/) | stomach | RNA-seq | R_ENCSR922VBO |
| Roadmap | [ENCFF794IMD](https://www.encodeproject.org/files/ENCFF794IMD/) | stomach | RNA-seq | R_ENCSR721HDG |
| Roadmap | [ENCFF433BMT](https://www.encodeproject.org/files/ENCFF433BMT/) | stomach | RNA-seq | R_ENCSR702IGQ |
| Roadmap | [ENCFF824JKN](https://www.encodeproject.org/files/ENCFF824JKN/) | stomach | RNA-seq | R_ENCSR549DVY |
| Roadmap | [ENCFF314DZG](https://www.encodeproject.org/files/ENCFF314DZG/) | stomach | RNA-seq | R_ENCSR783BUO |
| Roadmap | [ENCFF437SZX](https://www.encodeproject.org/files/ENCFF437SZX/) | stomach | RNA-seq | R_ENCSR783BUO |
| Roadmap | [ENCFF838LGI](https://www.encodeproject.org/files/ENCFF838LGI/) | stomach | RNA-seq | R_ENCSR951NPS |
| Roadmap | [ENCFF383KNO](https://www.encodeproject.org/files/ENCFF383KNO/) | stomach | RNA-seq | R_ENCSR123ZCX |
| Roadmap | [ENCFF701DLG](https://www.encodeproject.org/files/ENCFF701DLG/) | stomach | RNA-seq | R_ENCSR774SEX |
| Roadmap | [ENCFF326BCF](https://www.encodeproject.org/files/ENCFF326BCF/) | muscle of back | RNA-seq | R_ENCSR729ZII |
| Roadmap | [ENCFF851VPL](https://www.encodeproject.org/files/ENCFF851VPL/) | muscle of back | RNA-seq | R_ENCSR806ESH |
| Roadmap | [ENCFF093TMQ](https://www.encodeproject.org/files/ENCFF093TMQ/) | muscle of back | RNA-seq | R_ENCSR995ORR |
| Roadmap | [ENCFF670EYD](https://www.encodeproject.org/files/ENCFF670EYD/) | muscle of back | RNA-seq | R_ENCSR891JVD |
| Roadmap | [ENCFF473DKB](https://www.encodeproject.org/files/ENCFF473DKB/) | muscle of back | RNA-seq | R_ENCSR652AWW |
| Roadmap | [ENCFF447GUA](https://www.encodeproject.org/files/ENCFF447GUA/) | muscle of back | RNA-seq | R_ENCSR027EJD |
| Roadmap | [ENCFF630XRG](https://www.encodeproject.org/files/ENCFF630XRG/) | muscle of back | RNA-seq | R_ENCSR576UKA |
| Roadmap | [ENCFF096QFW](https://www.encodeproject.org/files/ENCFF096QFW/) | muscle of back | RNA-seq | R_ENCSR094RGI |
| Roadmap | [ENCFF683XJX](https://www.encodeproject.org/files/ENCFF683XJX/) | muscle of back | RNA-seq | R_ENCSR239BBI |
| Roadmap | [ENCFF015SQM](https://www.encodeproject.org/files/ENCFF015SQM/) | muscle of back | RNA-seq | R_ENCSR522XTV |
| Roadmap | [ENCFF025DKC](https://www.encodeproject.org/files/ENCFF025DKC/) | small intestine | RNA-seq | R_ENCSR719HRO |
| Roadmap | [ENCFF531PQC](https://www.encodeproject.org/files/ENCFF531PQC/) | small intestine | RNA-seq | R_ENCSR621FYE |
| Roadmap | [ENCFF796HDN](https://www.encodeproject.org/files/ENCFF796HDN/) | small intestine | RNA-seq | R_ENCSR150JIX |
| Roadmap | [ENCFF974KOL](https://www.encodeproject.org/files/ENCFF974KOL/) | small intestine | RNA-seq | R_ENCSR446RKD |
| Roadmap | [ENCFF677RLO](https://www.encodeproject.org/files/ENCFF677RLO/) | small intestine | RNA-seq | R_ENCSR523EDD |
| Roadmap | [ENCFF038OLY](https://www.encodeproject.org/files/ENCFF038OLY/) | muscle of leg | RNA-seq | R_ENCSR096USV |
| Roadmap | [ENCFF591YBG](https://www.encodeproject.org/files/ENCFF591YBG/) | muscle of leg | RNA-seq | R_ENCSR860DST |
| Roadmap | [ENCFF610CFP](https://www.encodeproject.org/files/ENCFF610CFP/) | muscle of leg | RNA-seq | R_ENCSR144UVO |
| Roadmap | [ENCFF726EJA](https://www.encodeproject.org/files/ENCFF726EJA/) | muscle of leg | RNA-seq | R_ENCSR144UVO |
| Roadmap | [ENCFF969AEA](https://www.encodeproject.org/files/ENCFF969AEA/) | muscle of leg | RNA-seq | R_ENCSR545WAC |
| Roadmap | [ENCFF280FJO](https://www.encodeproject.org/files/ENCFF280FJO/) | muscle of leg | RNA-seq | R_ENCSR174ESD |
| Roadmap | [ENCFF678IQL](https://www.encodeproject.org/files/ENCFF678IQL/) | muscle of leg | RNA-seq | R_ENCSR086DZF |
| Roadmap | [ENCFF755EWN](https://www.encodeproject.org/files/ENCFF755EWN/) | muscle of leg | RNA-seq | R_ENCSR561WEX |
| Roadmap | [ENCFF823WWJ](https://www.encodeproject.org/files/ENCFF823WWJ/) | muscle of leg | RNA-seq | R_ENCSR447IJE |
| Roadmap | [ENCFF800RTW](https://www.encodeproject.org/files/ENCFF800RTW/) | large intestine | RNA-seq | R_ENCSR286KWP |
| Roadmap | [ENCFF062QDJ](https://www.encodeproject.org/files/ENCFF062QDJ/) | large intestine | RNA-seq | R_ENCSR859KGW |
| Roadmap | [ENCFF376BOA](https://www.encodeproject.org/files/ENCFF376BOA/) | large intestine | RNA-seq | R_ENCSR859KGW |
| Roadmap | [ENCFF132LZV](https://www.encodeproject.org/files/ENCFF132LZV/) | large intestine | RNA-seq | R_ENCSR777ONH |
| Roadmap | [ENCFF600PXN](https://www.encodeproject.org/files/ENCFF600PXN/) | large intestine | RNA-seq | R_ENCSR930URM |
| Roadmap | [ENCFF203FUW](https://www.encodeproject.org/files/ENCFF203FUW/) | large intestine | RNA-seq | R_ENCSR857VKL |
| Roadmap | [ENCFF472AVS](https://www.encodeproject.org/files/ENCFF472AVS/) | large intestine | RNA-seq | R_ENCSR363BVC |
| Roadmap | [ENCFF994THJ](https://www.encodeproject.org/files/ENCFF994THJ/) | left lung | RNA-seq | R_ENCSR861SOG |
| Roadmap | [ENCFF565ABG](https://www.encodeproject.org/files/ENCFF565ABG/) | left lung | RNA-seq | R_ENCSR733MWN |
| Roadmap | [ENCFF836QCP](https://www.encodeproject.org/files/ENCFF836QCP/) | left lung | RNA-seq | R_ENCSR592EZK |
| Roadmap | [ENCFF810HYB](https://www.encodeproject.org/files/ENCFF810HYB/) | left lung | RNA-seq | R_ENCSR499NEL |
| Roadmap | [ENCFF527HNS](https://www.encodeproject.org/files/ENCFF527HNS/) | left lung | RNA-seq | R_ENCSR222IGR |
| Roadmap | [ENCFF391AKU](https://www.encodeproject.org/files/ENCFF391AKU/) | left lung | RNA-seq | R_ENCSR572FXC |
| Roadmap | [ENCFF715EGE](https://www.encodeproject.org/files/ENCFF715EGE/) | kidney | RNA-seq | R_ENCSR907KDH |
| Roadmap | [ENCFF391AJE](https://www.encodeproject.org/files/ENCFF391AJE/) | kidney | RNA-seq | R_ENCSR212AMA |
| Roadmap | [ENCFF390GZH](https://www.encodeproject.org/files/ENCFF390GZH/) | kidney | RNA-seq | R_ENCSR896QPD |
| Roadmap | [ENCFF959UJB](https://www.encodeproject.org/files/ENCFF959UJB/) | kidney | RNA-seq | R_ENCSR495UXA |
| Roadmap | [ENCFF240TXD](https://www.encodeproject.org/files/ENCFF240TXD/) | right lung | RNA-seq | R_ENCSR554KBK |
| Roadmap | [ENCFF994CGB](https://www.encodeproject.org/files/ENCFF994CGB/) | right lung | RNA-seq | R_ENCSR074APH |
| Roadmap | [ENCFF299JVW](https://www.encodeproject.org/files/ENCFF299JVW/) | right lung | RNA-seq | R_ENCSR560MDQ |
| Roadmap | [ENCFF567KUC](https://www.encodeproject.org/files/ENCFF567KUC/) | right lung | RNA-seq | R_ENCSR176WMG |
| Roadmap | [ENCFF527HRM](https://www.encodeproject.org/files/ENCFF527HRM/) | right lung | RNA-seq | R_ENCSR044JAQ |
| Roadmap | [ENCFF822QPE](https://www.encodeproject.org/files/ENCFF822QPE/) | thymus | RNA-seq | R_ENCSR367QHR |
| Roadmap | [ENCFF410KEZ](https://www.encodeproject.org/files/ENCFF410KEZ/) | thymus | RNA-seq | R_ENCSR158XIJ |
| Roadmap | [ENCFF715OSA](https://www.encodeproject.org/files/ENCFF715OSA/) | thymus | RNA-seq | R_ENCSR069CMT |
| Roadmap | [ENCFF199PMN](https://www.encodeproject.org/files/ENCFF199PMN/) | thymus | RNA-seq | R_ENCSR175CNQ |
| Roadmap | [ENCFF649RZG](https://www.encodeproject.org/files/ENCFF649RZG/) | thymus | RNA-seq | R_ENCSR175CNQ |
| Roadmap | [ENCFF975AUW](https://www.encodeproject.org/files/ENCFF975AUW/) | heart | RNA-seq | R_ENCSR047LLJ |
| Roadmap | [ENCFF092JPL](https://www.encodeproject.org/files/ENCFF092JPL/) | heart | RNA-seq | R_ENCSR863BUL |
| Roadmap | [ENCFF149FEQ](https://www.encodeproject.org/files/ENCFF149FEQ/) | renal cortex interstitium | RNA-seq | R_ENCSR328PVI |
| Roadmap | [ENCFF371QWQ](https://www.encodeproject.org/files/ENCFF371QWQ/) | renal cortex interstitium | RNA-seq | R_ENCSR899SWV |
| Roadmap | [ENCFF826YSU](https://www.encodeproject.org/files/ENCFF826YSU/) | renal cortex interstitium | RNA-seq | R_ENCSR436ZKE |
| Roadmap | [ENCFF051TUA](https://www.encodeproject.org/files/ENCFF051TUA/) | renal cortex interstitium | RNA-seq | R_ENCSR436ZKE |
| Roadmap | [ENCFF479AQV](https://www.encodeproject.org/files/ENCFF479AQV/) | adrenal gland | RNA-seq | R_ENCSR335GET |
| Roadmap | [ENCFF396DVW](https://www.encodeproject.org/files/ENCFF396DVW/) | adrenal gland | RNA-seq | R_ENCSR120NEA |
| Roadmap | [ENCFF080VMX](https://www.encodeproject.org/files/ENCFF080VMX/) | adrenal gland | RNA-seq | R_ENCSR688YOZ |
| Roadmap | [ENCFF345CEU](https://www.encodeproject.org/files/ENCFF345CEU/) | adrenal gland | RNA-seq | R_ENCSR740OPV |
| Roadmap | [ENCFF953VOF](https://www.encodeproject.org/files/ENCFF953VOF/) | renal pelvis | RNA-seq | R_ENCSR424TSZ |
| Roadmap | [ENCFF993WUV](https://www.encodeproject.org/files/ENCFF993WUV/) | renal pelvis | RNA-seq | R_ENCSR204XBB |
| Roadmap | [ENCFF329YXL](https://www.encodeproject.org/files/ENCFF329YXL/) | renal pelvis | RNA-seq | R_ENCSR929KRW |
| Roadmap | [ENCFF960GCL](https://www.encodeproject.org/files/ENCFF960GCL/) | renal pelvis | RNA-seq | R_ENCSR929KRW |
| Roadmap | [ENCFF956SGC](https://www.encodeproject.org/files/ENCFF956SGC/) | left kidney | RNA-seq | R_ENCSR702IMR |
| Roadmap | [ENCFF859IDO](https://www.encodeproject.org/files/ENCFF859IDO/) | left renal cortex interstitium | RNA-seq | R_ENCSR015EMF |
| Roadmap | [ENCFF832ENU](https://www.encodeproject.org/files/ENCFF832ENU/) | left renal cortex interstitium | RNA-seq | R_ENCSR125NGM |
| Roadmap | [ENCFF139JYD](https://www.encodeproject.org/files/ENCFF139JYD/) | left renal cortex interstitium | RNA-seq | R_ENCSR125NGM |
| Roadmap | [ENCFF356DVL](https://www.encodeproject.org/files/ENCFF356DVL/) | left renal cortex interstitium | RNA-seq | R_ENCSR759WPF |
| Roadmap | [ENCFF847ODW](https://www.encodeproject.org/files/ENCFF847ODW/) | left renal cortex interstitium | RNA-seq | R_ENCSR413LXW |
| Roadmap | [ENCFF490TMA](https://www.encodeproject.org/files/ENCFF490TMA/) | left renal pelvis | RNA-seq | R_ENCSR029FTY |
| Roadmap | [ENCFF453MGJ](https://www.encodeproject.org/files/ENCFF453MGJ/) | left renal pelvis | RNA-seq | R_ENCSR321ROU |
| Roadmap | [ENCFF592GJW](https://www.encodeproject.org/files/ENCFF592GJW/) | left renal pelvis | RNA-seq | R_ENCSR410DUZ |
| Roadmap | [ENCFF493PDU](https://www.encodeproject.org/files/ENCFF493PDU/) | left renal pelvis | RNA-seq | R_ENCSR160UAZ |
| Roadmap | [ENCFF754GRT](https://www.encodeproject.org/files/ENCFF754GRT/) | right renal pelvis | RNA-seq | R_ENCSR552YAE |
| Roadmap | [ENCFF791IHU](https://www.encodeproject.org/files/ENCFF791IHU/) | right renal pelvis | RNA-seq | R_ENCSR352GCS |
| Roadmap | [ENCFF260HSY](https://www.encodeproject.org/files/ENCFF260HSY/) | right renal pelvis | RNA-seq | R_ENCSR352GCS |
| Roadmap | [ENCFF178QHC](https://www.encodeproject.org/files/ENCFF178QHC/) | right renal pelvis | RNA-seq | R_ENCSR543TQW |
| Roadmap | [ENCFF853QBA](https://www.encodeproject.org/files/ENCFF853QBA/) | right renal pelvis | RNA-seq | R_ENCSR928CEQ |
| Roadmap | [ENCFF708BFX](https://www.encodeproject.org/files/ENCFF708BFX/) | spinal cord | RNA-seq | R_ENCSR899NLW |
| Roadmap | [ENCFF725KJB](https://www.encodeproject.org/files/ENCFF725KJB/) | spinal cord | RNA-seq | R_ENCSR899NLW |
| Roadmap | [ENCFF123FVX](https://www.encodeproject.org/files/ENCFF123FVX/) | spinal cord | RNA-seq | R_ENCSR333FZW |
| Roadmap | [ENCFF659JRI](https://www.encodeproject.org/files/ENCFF659JRI/) | spinal cord | RNA-seq | R_ENCSR333FZW |
| Roadmap | [ENCFF701WAW](https://www.encodeproject.org/files/ENCFF701WAW/) | right renal cortex interstitium | RNA-seq | R_ENCSR822AOE |
| Roadmap | [ENCFF589SOA](https://www.encodeproject.org/files/ENCFF589SOA/) | right renal cortex interstitium | RNA-seq | R_ENCSR884EVS |
| Roadmap | [ENCFF711XVI](https://www.encodeproject.org/files/ENCFF711XVI/) | right renal cortex interstitium | RNA-seq | R_ENCSR400DJE |
| Roadmap | [ENCFF098SSN](https://www.encodeproject.org/files/ENCFF098SSN/) | spleen | RNA-seq | R_ENCSR265NZF |
| Roadmap | [ENCFF732SOA](https://www.encodeproject.org/files/ENCFF732SOA/) | psoas muscle | RNA-seq | R_ENCSR817TLH |
| Roadmap | [ENCFF703HYW](https://www.encodeproject.org/files/ENCFF703HYW/) | muscle of trunk | RNA-seq | R_ENCSR531RKI |
| Roadmap | [ENCFF102XVK](https://www.encodeproject.org/files/ENCFF102XVK/) | ovary | RNA-seq | R_ENCSR727VTD |
| Roadmap | [ENCFF855YTN](https://www.encodeproject.org/files/ENCFF855YTN/) | ovary | RNA-seq | R_ENCSR725TPW |
| Roadmap | [ENCFF632WEO](https://www.encodeproject.org/files/ENCFF632WEO/) | pancreas | RNA-seq | R_ENCSR629VMZ |
| Roadmap | [ENCFF075UBV](https://www.encodeproject.org/files/ENCFF075UBV/) | pancreas | RNA-seq | R_ENCSR571BML |
| Roadmap | [ENCFF276UME](https://www.encodeproject.org/files/ENCFF276UME/) | testis | RNA-seq | R_ENCSR755LFM |
| Roadmap | [ENCFF673MWT](https://www.encodeproject.org/files/ENCFF673MWT/) | forelimb muscle | RNA-seq | R_ENCSR711NGL |
| Roadmap | [ENCFF976CLZ](https://www.encodeproject.org/files/ENCFF976CLZ/) | hindlimb muscle | RNA-seq | R_ENCSR516VDS |
| Roadmap | [ENCFF990ZUE](https://www.encodeproject.org/files/ENCFF990ZUE/) | H1-hESC | RNA-seq | R_ENCSR911GQI |
| Roadmap | [ENCFF307UVF](https://www.encodeproject.org/files/ENCFF307UVF/) | H1-hESC | RNA-seq | R_ENCSR844HLP |
| Blueprint | [ENCFF426NOK](https://www.encodeproject.org/files/ENCFF426NOK/) | skin fibroblast | DNase1-seq | R_ENCBS336CDQ |
| Blueprint | [ENCFF976EWT](https://www.encodeproject.org/files/ENCFF976EWT/) | skin fibroblast | DNase1-seq | R_ENCBS336CDQ |
| Blueprint | [ENCFF638STX](https://www.encodeproject.org/files/ENCFF638STX/) | skin fibroblast | DNase1-seq | R_ENCBS336CDQ |
| Blueprint | [ENCFF761MPW](https://www.encodeproject.org/files/ENCFF761MPW/) | skin fibroblast | DNase1-seq | R_ENCBS336CDQ |
| Blueprint | [ENCFF484CBD](https://www.encodeproject.org/files/ENCFF484CBD/) | skin fibroblast | DNase1-seq | R_ENCBS336CDQ |
| Blueprint | [ENCFF700BMG](https://www.encodeproject.org/files/ENCFF700BMG/) | skin fibroblast | DNase1-seq | R_ENCBS336CDQ |
| Blueprint | [ENCFF186NLY](https://www.encodeproject.org/files/ENCFF186NLY/) | skin fibroblast | DNase1-seq | R_ENCBS336CDQ |
| Blueprint | [ENCFF645VBV](https://www.encodeproject.org/files/ENCFF645VBV/) | skin fibroblast | DNase1-seq | R_ENCBS336CDQ |
| Blueprint | [ENCFF594NMP](https://www.encodeproject.org/files/ENCFF594NMP/) | skin fibroblast | DNase1-seq | R_ENCBS336CDQ |
| Blueprint | [ENCFF865ORR](https://www.encodeproject.org/files/ENCFF865ORR/) | skin fibroblast | DNase1-seq | R_ENCBS890NFL |
| Blueprint | [ENCFF593MYJ](https://www.encodeproject.org/files/ENCFF593MYJ/) | skin fibroblast | DNase1-seq | R_ENCBS890NFL |
| Blueprint | [ENCFF419BYU](https://www.encodeproject.org/files/ENCFF419BYU/) | skin fibroblast | DNase1-seq | R_ENCBS890NFL |
| Blueprint | [ENCFF836RXB](https://www.encodeproject.org/files/ENCFF836RXB/) | skin fibroblast | DNase1-seq | R_ENCBS890NFL |
| Blueprint | [ENCFF479NNH](https://www.encodeproject.org/files/ENCFF479NNH/) | skin fibroblast | DNase1-seq | R_ENCBS180EDA |
| Blueprint | [ENCFF899QJN](https://www.encodeproject.org/files/ENCFF899QJN/) | skin fibroblast | DNase1-seq | R_ENCBS180EDA |
| Blueprint | [ENCFF707PZQ](https://www.encodeproject.org/files/ENCFF707PZQ/) | fibroblast of skin of abdomen | DNase1-seq | R_ENCBS405WVO |
| Blueprint | [ENCFF972ZVY](https://www.encodeproject.org/files/ENCFF972ZVY/) | fibroblast of skin of abdomen | DNase1-seq | R_ENCBS405WVO |
| Blueprint | [ENCFF346SMK](https://www.encodeproject.org/files/ENCFF346SMK/) | fibroblast of skin of abdomen | DNase1-seq | R_ENCBS405WVO |
| Blueprint | [ENCFF491YLY](https://www.encodeproject.org/files/ENCFF491YLY/) | fibroblast of skin of abdomen | DNase1-seq | R_ENCBS405WVO |
| Blueprint | [ENCFF657JQO](https://www.encodeproject.org/files/ENCFF657JQO/) | fibroblast of skin of abdomen | DNase1-seq | R_ENCBS405WVO |
| Blueprint | <https://www.ncbi.nlm.nih.gov/geo/query/acc.cgi?acc=GSM878636> | fibroblast of skin of abdomen | DNase1-seq | R_ENCBS599YIE |
| Blueprint | [ENCFF070YBE](https://www.encodeproject.org/files/ENCFF070YBE/) | IMR-90 | DNase1-seq | R_ENCBS754TWW_ENCBS048TNH |
| Blueprint | [ENCFF175PZA](https://www.encodeproject.org/files/ENCFF175PZA/) | IMR-90 | DNase1-seq | R_ENCBS754TWW_ENCBS048TNH |
| Blueprint | [ENCFF119LCF](https://www.encodeproject.org/files/ENCFF119LCF/) | trophoblast cell | DNase1-seq | R_ENCBS150HBC_ENCBS376RZJ |
| Blueprint | [ENCFF047VYC](https://www.encodeproject.org/files/ENCFF047VYC/) | trophoblast cell | DNase1-seq | R_ENCBS150HBC_ENCBS376RZJ |
| Blueprint | [ENCFF091DTH](https://www.encodeproject.org/files/ENCFF091DTH/) | trophoblast cell | DNase1-seq | R_ENCBS150HBC_ENCBS376RZJ |
| Blueprint | [ENCFF982YFY](https://www.encodeproject.org/files/ENCFF982YFY/) | trophoblast cell | DNase1-seq | R_ENCBS150HBC_ENCBS376RZJ |
| Blueprint | [ENCFF356LUH](https://www.encodeproject.org/files/ENCFF356LUH/) | trophoblast cell | DNase1-seq | R_ENCBS150HBC_ENCBS376RZJ |
| Blueprint | [ENCFF846KKT](https://www.encodeproject.org/files/ENCFF846KKT/) | trophoblast cell | DNase1-seq | R_ENCBS150HBC_ENCBS376RZJ |
| Blueprint | [ENCFF894MKA](https://www.encodeproject.org/files/ENCFF894MKA/) | trophoblast cell | DNase1-seq | R_ENCBS150HBC_ENCBS376RZJ |
| Blueprint | [ENCFF149EUW](https://www.encodeproject.org/files/ENCFF149EUW/) | trophoblast cell | DNase1-seq | R_ENCBS150HBC_ENCBS376RZJ |
| Blueprint | [ENCFF385MYE](https://www.encodeproject.org/files/ENCFF385MYE/) | trophoblast cell | DNase1-seq | R_ENCBS150HBC_ENCBS376RZJ |
| Blueprint | [ENCFF654RBE](https://www.encodeproject.org/files/ENCFF654RBE/) | trophoblast cell | DNase1-seq | R_ENCBS150HBC_ENCBS376RZJ |
| Blueprint | [ENCFF671NGZ](https://www.encodeproject.org/files/ENCFF671NGZ/) | trophoblast cell | DNase1-seq | R_ENCBS150HBC_ENCBS376RZJ |
| Blueprint | [ENCFF695HOH](https://www.encodeproject.org/files/ENCFF695HOH/) | trophoblast cell | DNase1-seq | R_ENCBS150HBC_ENCBS376RZJ |
| Blueprint | [ENCFF145YHJ](https://www.encodeproject.org/files/ENCFF145YHJ/) | trophoblast cell | DNase1-seq | R_ENCBS150HBC_ENCBS376RZJ |
| Blueprint | [ENCFF041BSC](https://www.encodeproject.org/files/ENCFF041BSC/) | trophoblast cell | DNase1-seq | R_ENCBS150HBC_ENCBS376RZJ |
| Blueprint | [ENCFF278YNM](https://www.encodeproject.org/files/ENCFF278YNM/) | trophoblast cell | DNase1-seq | R_ENCBS150HBC_ENCBS376RZJ |
| Blueprint | [ENCFF144BFB](https://www.encodeproject.org/files/ENCFF144BFB/) | trophoblast cell | DNase1-seq | R_ENCBS150HBC_ENCBS376RZJ |
| Blueprint | [ENCFF968JUL](https://www.encodeproject.org/files/ENCFF968JUL/) | trophoblast cell | DNase1-seq | R_ENCBS150HBC_ENCBS376RZJ |
| Blueprint | [ENCFF459PAV](https://www.encodeproject.org/files/ENCFF459PAV/) | trophoblast cell | DNase1-seq | R_ENCBS150HBC_ENCBS376RZJ |
| Blueprint | [ENCFF149ODQ](https://www.encodeproject.org/files/ENCFF149ODQ/) | muscle of arm | DNase1-seq | R_ENCBS090AGL |
| Blueprint | [ENCFF960XRX](https://www.encodeproject.org/files/ENCFF960XRX/) | muscle of arm | DNase1-seq | R_ENCBS090AGL |
| Blueprint | [ENCFF941KYM](https://www.encodeproject.org/files/ENCFF941KYM/) | muscle of arm | DNase1-seq | R_ENCBS090AGL |
| Blueprint | [ENCFF896QEW](https://www.encodeproject.org/files/ENCFF896QEW/) | muscle of arm | DNase1-seq | R_ENCBS090AGL |
| Blueprint | [ENCFF273BFB](https://www.encodeproject.org/files/ENCFF273BFB/) | muscle of arm | DNase1-seq | R_ENCBS516CJQ |
| Blueprint | [ENCFF825SDM](https://www.encodeproject.org/files/ENCFF825SDM/) | muscle of arm | DNase1-seq | R_ENCBS516CJQ |
| Blueprint | [ENCFF027HWH](https://www.encodeproject.org/files/ENCFF027HWH/) | muscle of arm | DNase1-seq | R_ENCBS054WKY |
| Blueprint | [ENCFF415GQM](https://www.encodeproject.org/files/ENCFF415GQM/) | muscle of arm | DNase1-seq | R_ENCBS586CPQ |
| Blueprint | [ENCFF817VSJ](https://www.encodeproject.org/files/ENCFF817VSJ/) | muscle of arm | DNase1-seq | R_ENCBS586CPQ |
| Blueprint | [ENCFF840APC](https://www.encodeproject.org/files/ENCFF840APC/) | muscle of arm | DNase1-seq | R_ENCBS261IIB |
| Blueprint | [ENCFF337KND](https://www.encodeproject.org/files/ENCFF337KND/) | muscle of arm | DNase1-seq | R_ENCBS180RZG |
| Blueprint | [ENCFF443DXI](https://www.encodeproject.org/files/ENCFF443DXI/) | muscle of arm | DNase1-seq | R_ENCBS180RZG |
| Blueprint | [ENCFF794SFF](https://www.encodeproject.org/files/ENCFF794SFF/) | muscle of arm | DNase1-seq | R_ENCBS892WJE |
| Blueprint | [ENCFF500PCM](https://www.encodeproject.org/files/ENCFF500PCM/) | muscle of arm | DNase1-seq | R_ENCBS892WJE |
| Blueprint | [ENCFF400PCJ](https://www.encodeproject.org/files/ENCFF400PCJ/) | muscle of arm | DNase1-seq | R_ENCBS892WJE |
| Blueprint | [ENCFF187JEI](https://www.encodeproject.org/files/ENCFF187JEI/) | muscle of arm | DNase1-seq | R_ENCBS892WJE |
| Blueprint | [ENCFF762DDD](https://www.encodeproject.org/files/ENCFF762DDD/) | muscle of arm | DNase1-seq | R_ENCBS892WJE |
| Blueprint | [ENCFF677QYQ](https://www.encodeproject.org/files/ENCFF677QYQ/) | muscle of arm | DNase1-seq | R_ENCBS892WJE |
| Blueprint | [ENCFF709WUS](https://www.encodeproject.org/files/ENCFF709WUS/) | muscle of arm | DNase1-seq | R_ENCBS892WJE |
| Blueprint | [ENCFF244EEK](https://www.encodeproject.org/files/ENCFF244EEK/) | muscle of arm | DNase1-seq | R_ENCBS892WJE |
| Blueprint | [ENCFF243CEU](https://www.encodeproject.org/files/ENCFF243CEU/) | muscle of arm | DNase1-seq | R_ENCBS892WJE |
| Blueprint | [ENCFF502JHP](https://www.encodeproject.org/files/ENCFF502JHP/) | stomach | DNase1-seq | R_ENCBS578HBL |
| Blueprint | [ENCFF044OTP](https://www.encodeproject.org/files/ENCFF044OTP/) | stomach | DNase1-seq | R_ENCBS578HBL |
| Blueprint | [ENCFF726SST](https://www.encodeproject.org/files/ENCFF726SST/) | stomach | DNase1-seq | R_ENCBS441WEO |
| Blueprint | [ENCFF629RQV](https://www.encodeproject.org/files/ENCFF629RQV/) | stomach | DNase1-seq | R_ENCBS441WEO |
| Blueprint | [ENCFF727OLN](https://www.encodeproject.org/files/ENCFF727OLN/) | stomach | DNase1-seq | R_ENCBS220QDW |
| Blueprint | [ENCFF644PKR](https://www.encodeproject.org/files/ENCFF644PKR/) | stomach | DNase1-seq | R_ENCBS220QDW |
| Blueprint | [ENCFF288CYZ](https://www.encodeproject.org/files/ENCFF288CYZ/) | stomach | DNase1-seq | R_ENCBS220QDW |
| Blueprint | [ENCFF970FYA](https://www.encodeproject.org/files/ENCFF970FYA/) | stomach | DNase1-seq | R_ENCBS246MHN |
| Blueprint | [ENCFF141WGT](https://www.encodeproject.org/files/ENCFF141WGT/) | stomach | DNase1-seq | R_ENCBS291NHF |
| Blueprint | [ENCFF667XCZ](https://www.encodeproject.org/files/ENCFF667XCZ/) | stomach | DNase1-seq | R_ENCBS291NHF |
| Blueprint | [ENCFF953NRZ](https://www.encodeproject.org/files/ENCFF953NRZ/) | stomach | DNase1-seq | R_ENCBS291NHF |
| Blueprint | [ENCFF752LNT](https://www.encodeproject.org/files/ENCFF752LNT/) | stomach | DNase1-seq | R_ENCBS878MRX |
| Blueprint | [ENCFF424NPO](https://www.encodeproject.org/files/ENCFF424NPO/) | stomach | DNase1-seq | R_ENCBS878MRX |
| Blueprint | [ENCFF883FMW](https://www.encodeproject.org/files/ENCFF883FMW/) | stomach | DNase1-seq | R_ENCBS159PIU |
| Blueprint | [ENCFF088TXE](https://www.encodeproject.org/files/ENCFF088TXE/) | stomach | DNase1-seq | R_ENCBS159PIU |
| Blueprint | [ENCFF135RGN](https://www.encodeproject.org/files/ENCFF135RGN/) | stomach | DNase1-seq | R_ENCBS159PIU |
| Blueprint | [ENCFF985UXU](https://www.encodeproject.org/files/ENCFF985UXU/) | stomach | DNase1-seq | R_ENCBS159PIU |
| Blueprint | [ENCFF718MLB](https://www.encodeproject.org/files/ENCFF718MLB/) | stomach | DNase1-seq | R_ENCBS159PIU |
| Blueprint | [ENCFF780HTB](https://www.encodeproject.org/files/ENCFF780HTB/) | stomach | DNase1-seq | R_ENCBS159PIU |
| Blueprint | [ENCFF491MFA](https://www.encodeproject.org/files/ENCFF491MFA/) | stomach | DNase1-seq | R_ENCBS159PIU |
| Blueprint | [ENCFF426THC](https://www.encodeproject.org/files/ENCFF426THC/) | stomach | DNase1-seq | R_ENCBS159PIU |
| Blueprint | [ENCFF264JZE](https://www.encodeproject.org/files/ENCFF264JZE/) | stomach | DNase1-seq | R_ENCBS716QQK |
| Blueprint | [ENCFF990XCE](https://www.encodeproject.org/files/ENCFF990XCE/) | stomach | DNase1-seq | R_ENCBS716QQK |
| Blueprint | [ENCFF380UEX](https://www.encodeproject.org/files/ENCFF380UEX/) | muscle of back | DNase1-seq | R_ENCBS384LIR |
| Blueprint | [ENCFF304VFH](https://www.encodeproject.org/files/ENCFF304VFH/) | muscle of back | DNase1-seq | R_ENCBS384LIR |
| Blueprint | [ENCFF068GRT](https://www.encodeproject.org/files/ENCFF068GRT/) | muscle of back | DNase1-seq | R_ENCBS384LIR |
| Blueprint | [ENCFF041HUV](https://www.encodeproject.org/files/ENCFF041HUV/) | muscle of back | DNase1-seq | R_ENCBS384LIR |
| Blueprint | [ENCFF678RJO](https://www.encodeproject.org/files/ENCFF678RJO/) | muscle of back | DNase1-seq | R_ENCBS384LIR |
| Blueprint | [ENCFF368HUM](https://www.encodeproject.org/files/ENCFF368HUM/) | muscle of back | DNase1-seq | R_ENCBS384LIR |
| Blueprint | [ENCFF447RQF](https://www.encodeproject.org/files/ENCFF447RQF/) | muscle of back | DNase1-seq | R_ENCBS384LIR |
| Blueprint | [ENCFF216MSZ](https://www.encodeproject.org/files/ENCFF216MSZ/) | muscle of back | DNase1-seq | R_ENCBS174IGM |
| Blueprint | [ENCFF635TUS](https://www.encodeproject.org/files/ENCFF635TUS/) | muscle of back | DNase1-seq | R_ENCBS345TTL |
| Blueprint | [ENCFF972LVN](https://www.encodeproject.org/files/ENCFF972LVN/) | muscle of back | DNase1-seq | R_ENCBS345TTL |
| Blueprint | [ENCFF943APV](https://www.encodeproject.org/files/ENCFF943APV/) | muscle of back | DNase1-seq | R_ENCBS136SDO |
| Blueprint | [ENCFF296VDO](https://www.encodeproject.org/files/ENCFF296VDO/) | muscle of back | DNase1-seq | R_ENCBS136SDO |
| Blueprint | [ENCFF248XSX](https://www.encodeproject.org/files/ENCFF248XSX/) | muscle of back | DNase1-seq | R_ENCBS645HGQ |
| Blueprint | [ENCFF527MIJ](https://www.encodeproject.org/files/ENCFF527MIJ/) | muscle of back | DNase1-seq | R_ENCBS645HGQ |
| Blueprint | [ENCFF844ELO](https://www.encodeproject.org/files/ENCFF844ELO/) | muscle of back | DNase1-seq | R_ENCBS020XIW |
| Blueprint | [ENCFF608IJP](https://www.encodeproject.org/files/ENCFF608IJP/) | muscle of back | DNase1-seq | R_ENCBS020XIW |
| Blueprint | [ENCFF225CLL](https://www.encodeproject.org/files/ENCFF225CLL/) | muscle of back | DNase1-seq | R_ENCBS897YOR |
| Blueprint | [ENCFF673KNK](https://www.encodeproject.org/files/ENCFF673KNK/) | muscle of back | DNase1-seq | R_ENCBS897YOR |
| Blueprint | [ENCFF282HOP](https://www.encodeproject.org/files/ENCFF282HOP/) | muscle of back | DNase1-seq | R_ENCBS897YOR |
| Blueprint | [ENCFF873JMK](https://www.encodeproject.org/files/ENCFF873JMK/) | muscle of back | DNase1-seq | R_ENCBS479AOA |
| Blueprint | [ENCFF961ZLA](https://www.encodeproject.org/files/ENCFF961ZLA/) | muscle of back | DNase1-seq | R_ENCBS479AOA |
| Blueprint | [ENCFF532NCU](https://www.encodeproject.org/files/ENCFF532NCU/) | muscle of back | DNase1-seq | R_ENCBS479AOA |
| Blueprint | [ENCFF347UTJ](https://www.encodeproject.org/files/ENCFF347UTJ/) | muscle of back | DNase1-seq | R_ENCBS479AOA |
| Blueprint | [ENCFF446XVP](https://www.encodeproject.org/files/ENCFF446XVP/) | muscle of back | DNase1-seq | R_ENCBS479AOA |
| Blueprint | [ENCFF914LBJ](https://www.encodeproject.org/files/ENCFF914LBJ/) | muscle of back | DNase1-seq | R_ENCBS479AOA |
| Blueprint | [ENCFF055BON](https://www.encodeproject.org/files/ENCFF055BON/) | muscle of back | DNase1-seq | R_ENCBS479AOA |
| Blueprint | [ENCFF683XJX](https://www.encodeproject.org/files/ENCFF683XJX/) | muscle of back | DNase1-seq | R_ENCBS825MQT |
| Blueprint | [ENCFF735BDG](https://www.encodeproject.org/files/ENCFF735BDG/) | muscle of back | DNase1-seq | R_ENCBS136EGD |
| Blueprint | [ENCFF904JYN](https://www.encodeproject.org/files/ENCFF904JYN/) | muscle of back | DNase1-seq | R_ENCBS136EGD |
| Blueprint | [ENCFF102YUC](https://www.encodeproject.org/files/ENCFF102YUC/) | muscle of back | DNase1-seq | R_ENCBS136EGD |
| Blueprint | [ENCFF198YRW](https://www.encodeproject.org/files/ENCFF198YRW/) | muscle of back | DNase1-seq | R_ENCBS136EGD |
| Blueprint | [ENCFF665XQC](https://www.encodeproject.org/files/ENCFF665XQC/) | small intestine | DNase1-seq | R_ENCBS853LFM |
| Blueprint | [ENCFF946FXI](https://www.encodeproject.org/files/ENCFF946FXI/) | small intestine | DNase1-seq | R_ENCBS853LFM |
| Blueprint | [ENCFF473FIC](https://www.encodeproject.org/files/ENCFF473FIC/) | small intestine | DNase1-seq | R_ENCBS853LFM |
| Blueprint | [ENCFF373XZL](https://www.encodeproject.org/files/ENCFF373XZL/) | small intestine | DNase1-seq | R_ENCBS853LFM |
| Blueprint | [ENCFF570FNC](https://www.encodeproject.org/files/ENCFF570FNC/) | small intestine | DNase1-seq | R_ENCBS615YKY |
| Blueprint | [ENCFF463POG](https://www.encodeproject.org/files/ENCFF463POG/) | small intestine | DNase1-seq | R_ENCBS529UES |
| Blueprint | [ENCFF698BGB](https://www.encodeproject.org/files/ENCFF698BGB/) | small intestine | DNase1-seq | R_ENCBS529UES |
| Blueprint | [ENCFF098RZZ](https://www.encodeproject.org/files/ENCFF098RZZ/) | small intestine | DNase1-seq | R_ENCBS529UES |
| Blueprint | [ENCFF404NLX](https://www.encodeproject.org/files/ENCFF404NLX/) | small intestine | DNase1-seq | R_ENCBS623YHX |
| Blueprint | [ENCFF670PIU](https://www.encodeproject.org/files/ENCFF670PIU/) | small intestine | DNase1-seq | R_ENCBS133LAN |
| Blueprint | [ENCFF774AET](https://www.encodeproject.org/files/ENCFF774AET/) | small intestine | DNase1-seq | R_ENCBS133LAN |
| Blueprint | [ENCFF173RST](https://www.encodeproject.org/files/ENCFF173RST/) | muscle of leg | DNase1-seq | R_ENCBS611ZBY |
| Blueprint | [ENCFF289VMW](https://www.encodeproject.org/files/ENCFF289VMW/) | muscle of leg | DNase1-seq | R_ENCBS517FUR |
| Blueprint | [ENCFF401RGR](https://www.encodeproject.org/files/ENCFF401RGR/) | muscle of leg | DNase1-seq | R_ENCBS517FUR |
| Blueprint | [ENCFF050CPH](https://www.encodeproject.org/files/ENCFF050CPH/) | muscle of leg | DNase1-seq | R_ENCBS023IXF |
| Blueprint | [ENCFF396CDS](https://www.encodeproject.org/files/ENCFF396CDS/) | muscle of leg | DNase1-seq | R_ENCBS023IXF |
| Blueprint | [ENCFF018ZYC](https://www.encodeproject.org/files/ENCFF018ZYC/) | muscle of leg | DNase1-seq | R_ENCBS947JRD |
| Blueprint | [ENCFF667LLC](https://www.encodeproject.org/files/ENCFF667LLC/) | muscle of leg | DNase1-seq | R_ENCBS947JRD |
| Blueprint | [ENCFF843QNZ](https://www.encodeproject.org/files/ENCFF843QNZ/) | muscle of leg | DNase1-seq | R_ENCBS947JRD |
| Blueprint | [ENCFF892QTV](https://www.encodeproject.org/files/ENCFF892QTV/) | muscle of leg | DNase1-seq | R_ENCBS011TVS |
| Blueprint | [ENCFF941COX](https://www.encodeproject.org/files/ENCFF941COX/) | muscle of leg | DNase1-seq | R_ENCBS011TVS |
| Blueprint | [ENCFF005KIH](https://www.encodeproject.org/files/ENCFF005KIH/) | muscle of leg | DNase1-seq | R_ENCBS011TVS |
| Blueprint | [ENCFF968NLM](https://www.encodeproject.org/files/ENCFF968NLM/) | muscle of leg | DNase1-seq | R_ENCBS011TVS |
| Blueprint | [ENCFF308AWG](https://www.encodeproject.org/files/ENCFF308AWG/) | muscle of leg | DNase1-seq | R_ENCBS011TVS |
| Blueprint | [ENCFF419QCQ](https://www.encodeproject.org/files/ENCFF419QCQ/) | muscle of leg | DNase1-seq | R_ENCBS011TVS |
| Blueprint | [ENCFF987FBY](https://www.encodeproject.org/files/ENCFF987FBY/) | muscle of leg | DNase1-seq | R_ENCBS011TVS |
| Blueprint | [ENCFF103FXL](https://www.encodeproject.org/files/ENCFF103FXL/) | muscle of leg | DNase1-seq | R_ENCBS011TVS |
| Blueprint | [ENCFF916CUH](https://www.encodeproject.org/files/ENCFF916CUH/) | muscle of leg | DNase1-seq | R_ENCBS099OIO |
| Blueprint | [ENCFF792OYK](https://www.encodeproject.org/files/ENCFF792OYK/) | muscle of leg | DNase1-seq | R_ENCBS099OIO |
| Blueprint | [ENCFF786MHH](https://www.encodeproject.org/files/ENCFF786MHH/) | muscle of leg | DNase1-seq | R_ENCBS099OIO |
| Blueprint | [ENCFF241GTO](https://www.encodeproject.org/files/ENCFF241GTO/) | muscle of leg | DNase1-seq | R_ENCBS099OIO |
| Blueprint | [ENCFF481YWO](https://www.encodeproject.org/files/ENCFF481YWO/) | muscle of leg | DNase1-seq | R_ENCBS099OIO |
| Blueprint | [ENCFF505FYZ](https://www.encodeproject.org/files/ENCFF505FYZ/) | muscle of leg | DNase1-seq | R_ENCBS099OIO |
| Blueprint | [ENCFF269DFT](https://www.encodeproject.org/files/ENCFF269DFT/) | muscle of leg | DNase1-seq | R_ENCBS099OIO |
| Blueprint | [ENCFF535RGT](https://www.encodeproject.org/files/ENCFF535RGT/) | muscle of leg | DNase1-seq | R_ENCBS099OIO |
| Blueprint | [ENCFF175SLR](https://www.encodeproject.org/files/ENCFF175SLR/) | muscle of leg | DNase1-seq | R_ENCBS099OIO |
| Blueprint | [ENCFF101OMS](https://www.encodeproject.org/files/ENCFF101OMS/) | muscle of leg | DNase1-seq | R_ENCBS099OIO |
| Blueprint | [ENCFF361UWR](https://www.encodeproject.org/files/ENCFF361UWR/) | muscle of leg | DNase1-seq | R_ENCBS143XQJ |
| Blueprint | [ENCFF742WIO](https://www.encodeproject.org/files/ENCFF742WIO/) | muscle of leg | DNase1-seq | R_ENCBS143XQJ |
| Blueprint | [ENCFF272WWY](https://www.encodeproject.org/files/ENCFF272WWY/) | muscle of leg | DNase1-seq | R_ENCBS143XQJ |
| Blueprint | [ENCFF808QSC](https://www.encodeproject.org/files/ENCFF808QSC/) | muscle of leg | DNase1-seq | R_ENCBS143XQJ |
| Blueprint | [ENCFF128KEC](https://www.encodeproject.org/files/ENCFF128KEC/) | muscle of leg | DNase1-seq | R_ENCBS984JKS |
| Blueprint | [ENCFF201FIA](https://www.encodeproject.org/files/ENCFF201FIA/) | muscle of leg | DNase1-seq | R_ENCBS984JKS |
| Blueprint | [ENCFF611HRT](https://www.encodeproject.org/files/ENCFF611HRT/) | muscle of leg | DNase1-seq | R_ENCBS984JKS |
| Blueprint | [ENCFF365KQE](https://www.encodeproject.org/files/ENCFF365KQE/) | muscle of leg | DNase1-seq | R_ENCBS984JKS |
| Blueprint | [ENCFF365NYW](https://www.encodeproject.org/files/ENCFF365NYW/) | muscle of leg | DNase1-seq | R_ENCBS984JKS |
| Blueprint | [ENCFF743SDH](https://www.encodeproject.org/files/ENCFF743SDH/) | large intestine | DNase1-seq | R_ENCBS997WGU |
| Blueprint | [ENCFF004QQU](https://www.encodeproject.org/files/ENCFF004QQU/) | large intestine | DNase1-seq | R_ENCBS997WGU |
| Blueprint | [ENCFF927LGL](https://www.encodeproject.org/files/ENCFF927LGL/) | large intestine | DNase1-seq | R_ENCBS588ZWT |
| Blueprint | [ENCFF144ZQA](https://www.encodeproject.org/files/ENCFF144ZQA/) | large intestine | DNase1-seq | R_ENCBS445IVN |
| Blueprint | [ENCFF895VUK](https://www.encodeproject.org/files/ENCFF895VUK/) | large intestine | DNase1-seq | R_ENCBS445IVN |
| Blueprint | [ENCFF181QVU](https://www.encodeproject.org/files/ENCFF181QVU/) | large intestine | DNase1-seq | R_ENCBS699KFK |
| Blueprint | [ENCFF271BCH](https://www.encodeproject.org/files/ENCFF271BCH/) | large intestine | DNase1-seq | R_ENCBS867ILV |
| Blueprint | [ENCFF122XCL](https://www.encodeproject.org/files/ENCFF122XCL/) | large intestine | DNase1-seq | R_ENCBS867ILV |
| Blueprint | [ENCFF528OCE](https://www.encodeproject.org/files/ENCFF528OCE/) | large intestine | DNase1-seq | R_ENCBS383OVQ |
| Blueprint | [ENCFF070ZPW](https://www.encodeproject.org/files/ENCFF070ZPW/) | large intestine | DNase1-seq | R_ENCBS383OVQ |
| Blueprint | [ENCFF942PFR](https://www.encodeproject.org/files/ENCFF942PFR/) | left lung | DNase1-seq | R_ENCBS078XUR |
| Blueprint | [ENCFF639RTQ](https://www.encodeproject.org/files/ENCFF639RTQ/) | left lung | DNase1-seq | R_ENCBS574MIZ |
| Blueprint | [ENCFF641TVF](https://www.encodeproject.org/files/ENCFF641TVF/) | left lung | DNase1-seq | R_ENCBS859ASH |
| Blueprint | [ENCFF666OGL](https://www.encodeproject.org/files/ENCFF666OGL/) | left lung | DNase1-seq | R_ENCBS143LJK |
| Blueprint | [ENCFF461HIP](https://www.encodeproject.org/files/ENCFF461HIP/) | left lung | DNase1-seq | R_ENCBS143LJK |
| Blueprint | [ENCFF244DOB](https://www.encodeproject.org/files/ENCFF244DOB/) | left lung | DNase1-seq | R_ENCBS143LJK |
| Blueprint | [ENCFF497CUV](https://www.encodeproject.org/files/ENCFF497CUV/) | left lung | DNase1-seq | R_ENCBS143LJK |
| Blueprint | [ENCFF551FHH](https://www.encodeproject.org/files/ENCFF551FHH/) | left lung | DNase1-seq | R_ENCBS143LJK |
| Blueprint | [ENCFF977WQN](https://www.encodeproject.org/files/ENCFF977WQN/) | left lung | DNase1-seq | R_ENCBS143LJK |
| Blueprint | [ENCFF925NBM](https://www.encodeproject.org/files/ENCFF925NBM/) | left lung | DNase1-seq | R_ENCBS143LJK |
| Blueprint | [ENCFF999PLN](https://www.encodeproject.org/files/ENCFF999PLN/) | left lung | DNase1-seq | R_ENCBS143LJK |
| Blueprint | [ENCFF135VOG](https://www.encodeproject.org/files/ENCFF135VOG/) | left lung | DNase1-seq | R_ENCBS143LJK |
| Blueprint | [ENCFF450MOY](https://www.encodeproject.org/files/ENCFF450MOY/) | left lung | DNase1-seq | R_ENCBS516MKG |
| Blueprint | [ENCFF962CEA](https://www.encodeproject.org/files/ENCFF962CEA/) | left lung | DNase1-seq | R_ENCBS516MKG |
| Blueprint | [ENCFF191FJK](https://www.encodeproject.org/files/ENCFF191FJK/) | left lung | DNase1-seq | R_ENCBS117CVU |
| Blueprint | [ENCFF145NEL](https://www.encodeproject.org/files/ENCFF145NEL/) | left lung | DNase1-seq | R_ENCBS117CVU |
| Blueprint | [ENCFF347TBD](https://www.encodeproject.org/files/ENCFF347TBD/) | left lung | DNase1-seq | R_ENCBS117CVU |
| Blueprint | [ENCFF272FPH](https://www.encodeproject.org/files/ENCFF272FPH/) | kidney | DNase1-seq | R_ENCBS478OZL |
| Blueprint | [ENCFF083NYS](https://www.encodeproject.org/files/ENCFF083NYS/) | kidney | DNase1-seq | R_ENCBS478OZL |
| Blueprint | [ENCFF916YMJ](https://www.encodeproject.org/files/ENCFF916YMJ/) | kidney | DNase1-seq | R_ENCBS263ZZU |
| Blueprint | [ENCFF799XLW](https://www.encodeproject.org/files/ENCFF799XLW/) | kidney | DNase1-seq | R_ENCBS263ZZU |
| Blueprint | [ENCFF618FCN](https://www.encodeproject.org/files/ENCFF618FCN/) | kidney | DNase1-seq | R_ENCBS263ZZU |
| Blueprint | [ENCFF970TXW](https://www.encodeproject.org/files/ENCFF970TXW/) | kidney | DNase1-seq | R_ENCBS434EOI |
| Blueprint | [ENCFF009RHR](https://www.encodeproject.org/files/ENCFF009RHR/) | kidney | DNase1-seq | R_ENCBS434EOI |
| Blueprint | [ENCFF196NVH](https://www.encodeproject.org/files/ENCFF196NVH/) | kidney | DNase1-seq | R_ENCBS434EOI |
| Blueprint | [ENCFF753ELF](https://www.encodeproject.org/files/ENCFF753ELF/) | kidney | DNase1-seq | R_ENCBS034SKE |
| Blueprint | [ENCFF286XQY](https://www.encodeproject.org/files/ENCFF286XQY/) | kidney | DNase1-seq | R_ENCBS034SKE |
| Blueprint | [ENCFF416YBE](https://www.encodeproject.org/files/ENCFF416YBE/) | right lung | DNase1-seq | R_ENCBS917VNB |
| Blueprint | [ENCFF120FQU](https://www.encodeproject.org/files/ENCFF120FQU/) | right lung | DNase1-seq | R_ENCBS917VNB |
| Blueprint | [ENCFF427YAL](https://www.encodeproject.org/files/ENCFF427YAL/) | right lung | DNase1-seq | R_ENCBS993PWO |
| Blueprint | [ENCFF346VUR](https://www.encodeproject.org/files/ENCFF346VUR/) | right lung | DNase1-seq | R_ENCBS993PWO |
| Blueprint | [ENCFF776KRS](https://www.encodeproject.org/files/ENCFF776KRS/) | right lung | DNase1-seq | R_ENCBS467PJB |
| Blueprint | [ENCFF922IPA](https://www.encodeproject.org/files/ENCFF922IPA/) | right lung | DNase1-seq | R_ENCBS467PJB |
| Blueprint | [ENCFF257CYL](https://www.encodeproject.org/files/ENCFF257CYL/) | right lung | DNase1-seq | R_ENCBS421OZO |
| Blueprint | [ENCFF514LUU](https://www.encodeproject.org/files/ENCFF514LUU/) | right lung | DNase1-seq | R_ENCBS421OZO |
| Blueprint | [ENCFF083OYH](https://www.encodeproject.org/files/ENCFF083OYH/) | right lung | DNase1-seq | R_ENCBS421OZO |
| Blueprint | [ENCFF723TEH](https://www.encodeproject.org/files/ENCFF723TEH/) | right lung | DNase1-seq | R_ENCBS122USS |
| Blueprint | [ENCFF973AFO](https://www.encodeproject.org/files/ENCFF973AFO/) | right lung | DNase1-seq | R_ENCBS122USS |
| Blueprint | [ENCFF702CZA](https://www.encodeproject.org/files/ENCFF702CZA/) | thymus | DNase1-seq | R_ENCBS484BGT |
| Blueprint | [ENCFF714BGJ](https://www.encodeproject.org/files/ENCFF714BGJ/) | thymus | DNase1-seq | R_ENCBS484BGT |
| Blueprint | [ENCFF284FZH](https://www.encodeproject.org/files/ENCFF284FZH/) | thymus | DNase1-seq | R_ENCBS948PMG |
| Blueprint | [ENCFF496OMB](https://www.encodeproject.org/files/ENCFF496OMB/) | thymus | DNase1-seq | R_ENCBS948PMG |
| Blueprint | [ENCFF901HTM](https://www.encodeproject.org/files/ENCFF901HTM/) | thymus | DNase1-seq | R_ENCBS948PMG |
| Blueprint | [ENCFF939NEZ](https://www.encodeproject.org/files/ENCFF939NEZ/) | thymus | DNase1-seq | R_ENCBS054CPR |
| Blueprint | [ENCFF518NCM](https://www.encodeproject.org/files/ENCFF518NCM/) | thymus | DNase1-seq | R_ENCBS054CPR |
| Blueprint | [ENCFF806QFV](https://www.encodeproject.org/files/ENCFF806QFV/) | thymus | DNase1-seq | R_ENCBS054CPR |
| Blueprint | [ENCFF155HAH](https://www.encodeproject.org/files/ENCFF155HAH/) | thymus | DNase1-seq | R_ENCBS198CXJ |
| Blueprint | [ENCFF681JOT](https://www.encodeproject.org/files/ENCFF681JOT/) | thymus | DNase1-seq | R_ENCBS198CXJ |
| Blueprint | [ENCFF707FJO](https://www.encodeproject.org/files/ENCFF707FJO/) | thymus | DNase1-seq | R_ENCBS198CXJ |
| Blueprint | [ENCFF870IEV](https://www.encodeproject.org/files/ENCFF870IEV/) | thymus | DNase1-seq | R_ENCBS198CXJ |
| Blueprint | [ENCFF328VXJ](https://www.encodeproject.org/files/ENCFF328VXJ/) | thymus | DNase1-seq | R_ENCBS198CXJ |
| Blueprint | [ENCFF625QAW](https://www.encodeproject.org/files/ENCFF625QAW/) | thymus | DNase1-seq | R_ENCBS198CXJ |
| Blueprint | [ENCFF278ZNT](https://www.encodeproject.org/files/ENCFF278ZNT/) | heart | DNase1-seq | R_ENCBS172XKB |
| Blueprint | [ENCFF210JWW](https://www.encodeproject.org/files/ENCFF210JWW/) | heart | DNase1-seq | R_ENCBS172XKB |
| Blueprint | [ENCFF943TNE](https://www.encodeproject.org/files/ENCFF943TNE/) | heart | DNase1-seq | R_ENCBS172XKB |
| Blueprint | [ENCFF151KDO](https://www.encodeproject.org/files/ENCFF151KDO/) | heart | DNase1-seq | R_ENCBS172XKB |
| Blueprint | [ENCFF416ZBD](https://www.encodeproject.org/files/ENCFF416ZBD/) | heart | DNase1-seq | R_ENCBS172XKB |
| Blueprint | [ENCFF106CPX](https://www.encodeproject.org/files/ENCFF106CPX/) | heart | DNase1-seq | R_ENCBS407ALA |
| Blueprint | [ENCFF923BVC](https://www.encodeproject.org/files/ENCFF923BVC/) | heart | DNase1-seq | R_ENCBS407ALA |
| Blueprint | [ENCFF007SPW](https://www.encodeproject.org/files/ENCFF007SPW/) | heart | DNase1-seq | R_ENCBS407ALA |
| Blueprint | [ENCFF311OBU](https://www.encodeproject.org/files/ENCFF311OBU/) | heart | DNase1-seq | R_ENCBS407ALA |
| Blueprint | [ENCFF452AYP](https://www.encodeproject.org/files/ENCFF452AYP/) | heart | DNase1-seq | R_ENCBS407ALA |
| Blueprint | [ENCFF450AJZ](https://www.encodeproject.org/files/ENCFF450AJZ/) | renal cortex interstitium | DNase1-seq | R_ENCBS620YJZ |
| Blueprint | [ENCFF761BFW](https://www.encodeproject.org/files/ENCFF761BFW/) | renal cortex interstitium | DNase1-seq | R_ENCBS448HVV |
| Blueprint | [ENCFF553VIY](https://www.encodeproject.org/files/ENCFF553VIY/) | renal cortex interstitium | DNase1-seq | R_ENCBS026RJE |
| Blueprint | [ENCFF971PIK](https://www.encodeproject.org/files/ENCFF971PIK/) | renal cortex interstitium | DNase1-seq | R_ENCBS026RJE |
| Blueprint | [ENCFF697SCS](https://www.encodeproject.org/files/ENCFF697SCS/) | renal cortex interstitium | DNase1-seq | R_ENCBS026RJE |
| Blueprint | [ENCFF289UKR](https://www.encodeproject.org/files/ENCFF289UKR/) | renal cortex interstitium | DNase1-seq | R_ENCBS026RJE |
| Blueprint | [ENCFF206TOI](https://www.encodeproject.org/files/ENCFF206TOI/) | adrenal gland | DNase1-seq | R_ENCBS232ONZ |
| Blueprint | [ENCFF129QFX](https://www.encodeproject.org/files/ENCFF129QFX/) | adrenal gland | DNase1-seq | R_ENCBS232ONZ |
| Blueprint | [ENCFF461SIX](https://www.encodeproject.org/files/ENCFF461SIX/) | adrenal gland | DNase1-seq | R_ENCBS660CJK |
| Blueprint | [ENCFF294INJ](https://www.encodeproject.org/files/ENCFF294INJ/) | adrenal gland | DNase1-seq | R_ENCBS660CJK |
| Blueprint | [ENCFF414WFK](https://www.encodeproject.org/files/ENCFF414WFK/) | adrenal gland | DNase1-seq | R_ENCBS200XAZ |
| Blueprint | [ENCFF149EPY](https://www.encodeproject.org/files/ENCFF149EPY/) | adrenal gland | DNase1-seq | R_ENCBS200XAZ |
| Blueprint | [ENCFF529NSR](https://www.encodeproject.org/files/ENCFF529NSR/) | adrenal gland | DNase1-seq | R_ENCBS200XAZ |
| Blueprint | [ENCFF934GEX](https://www.encodeproject.org/files/ENCFF934GEX/) | adrenal gland | DNase1-seq | R_ENCBS200XAZ |
| Blueprint | [ENCFF564QHA](https://www.encodeproject.org/files/ENCFF564QHA/) | adrenal gland | DNase1-seq | R_ENCBS200XAZ |
| Blueprint | [ENCFF702FIW](https://www.encodeproject.org/files/ENCFF702FIW/) | adrenal gland | DNase1-seq | R_ENCBS200XAZ |
| Blueprint | [ENCFF916IPH](https://www.encodeproject.org/files/ENCFF916IPH/) | adrenal gland | DNase1-seq | R_ENCBS200XAZ |
| Blueprint | [ENCFF217BAD](https://www.encodeproject.org/files/ENCFF217BAD/) | adrenal gland | DNase1-seq | R_ENCBS119WRO |
| Blueprint | [ENCFF783HCO](https://www.encodeproject.org/files/ENCFF783HCO/) | adrenal gland | DNase1-seq | R_ENCBS119WRO |
| Blueprint | [ENCFF828MHF](https://www.encodeproject.org/files/ENCFF828MHF/) | adrenal gland | DNase1-seq | R_ENCBS119WRO |
| Blueprint | [ENCFF111FYY](https://www.encodeproject.org/files/ENCFF111FYY/) | adrenal gland | DNase1-seq | R_ENCBS119WRO |
| Blueprint | [ENCFF045ZNO](https://www.encodeproject.org/files/ENCFF045ZNO/) | renal pelvis | DNase1-seq | R_ENCBS262QPN |
| Blueprint | [ENCFF785AHD](https://www.encodeproject.org/files/ENCFF785AHD/) | renal pelvis | DNase1-seq | R_ENCBS262QPN |
| Blueprint | [ENCFF568EUW](https://www.encodeproject.org/files/ENCFF568EUW/) | renal pelvis | DNase1-seq | R_ENCBS142XDW |
| Blueprint | [ENCFF779WIH](https://www.encodeproject.org/files/ENCFF779WIH/) | renal pelvis | DNase1-seq | R_ENCBS142XDW |
| Blueprint | [ENCFF287NFY](https://www.encodeproject.org/files/ENCFF287NFY/) | renal pelvis | DNase1-seq | R_ENCBS142XDW |
| Blueprint | [ENCFF678HJR](https://www.encodeproject.org/files/ENCFF678HJR/) | renal pelvis | DNase1-seq | R_ENCBS785KTZ |
| Blueprint | [ENCFF539CMH](https://www.encodeproject.org/files/ENCFF539CMH/) | renal pelvis | DNase1-seq | R_ENCBS785KTZ |
| Blueprint | [ENCFF128PNW](https://www.encodeproject.org/files/ENCFF128PNW/) | renal pelvis | DNase1-seq | R_ENCBS785KTZ |
| Blueprint | [ENCFF315MZE](https://www.encodeproject.org/files/ENCFF315MZE/) | renal pelvis | DNase1-seq | R_ENCBS785KTZ |
| Blueprint | [ENCFF048UBW](https://www.encodeproject.org/files/ENCFF048UBW/) | left kidney | DNase1-seq | R_ENCBS610XAP |
| Blueprint | [ENCFF467BCS](https://www.encodeproject.org/files/ENCFF467BCS/) | left kidney | DNase1-seq | R_ENCBS610XAP |
| Blueprint | [ENCFF629QCK](https://www.encodeproject.org/files/ENCFF629QCK/) | left renal cortex interstitium | DNase1-seq | R_ENCBS376PWL |
| Blueprint | [ENCFF776RFT](https://www.encodeproject.org/files/ENCFF776RFT/) | left renal cortex interstitium | DNase1-seq | R_ENCBS376PWL |
| Blueprint | [ENCFF711KKQ](https://www.encodeproject.org/files/ENCFF711KKQ/) | left renal cortex interstitium | DNase1-seq | R_ENCBS674SNK |
| Blueprint | [ENCFF159XPF](https://www.encodeproject.org/files/ENCFF159XPF/) | left renal cortex interstitium | DNase1-seq | R_ENCBS674SNK |
| Blueprint | [ENCFF072IHB](https://www.encodeproject.org/files/ENCFF072IHB/) | left renal cortex interstitium | DNase1-seq | R_ENCBS281CNH |
| Blueprint | [ENCFF024ZWA](https://www.encodeproject.org/files/ENCFF024ZWA/) | left renal cortex interstitium | DNase1-seq | R_ENCBS281CNH |
| Blueprint | [ENCFF155MWH](https://www.encodeproject.org/files/ENCFF155MWH/) | left renal cortex interstitium | DNase1-seq | R_ENCBS636QOC |
| Blueprint | [ENCFF874CGL](https://www.encodeproject.org/files/ENCFF874CGL/) | left renal cortex interstitium | DNase1-seq | R_ENCBS636QOC |
| Blueprint | [ENCFF083MLN](https://www.encodeproject.org/files/ENCFF083MLN/) | left renal cortex interstitium | DNase1-seq | R_ENCBS636QOC |
| Blueprint | [ENCFF785JRW](https://www.encodeproject.org/files/ENCFF785JRW/) | left renal cortex interstitium | DNase1-seq | R_ENCBS636QOC |
| Blueprint | [ENCFF991TVK](https://www.encodeproject.org/files/ENCFF991TVK/) | left renal cortex interstitium | DNase1-seq | R_ENCBS636QOC |
| Blueprint | [ENCFF417LLB](https://www.encodeproject.org/files/ENCFF417LLB/) | left renal cortex interstitium | DNase1-seq | R_ENCBS636QOC |
| Blueprint | [ENCFF584SLG](https://www.encodeproject.org/files/ENCFF584SLG/) | left renal cortex interstitium | DNase1-seq | R_ENCBS636QOC |
| Blueprint | [ENCFF954SPU](https://www.encodeproject.org/files/ENCFF954SPU/) | left renal cortex interstitium | DNase1-seq | R_ENCBS636QOC |
| Blueprint | [ENCFF423HOT](https://www.encodeproject.org/files/ENCFF423HOT/) | left renal pelvis | DNase1-seq | R_ENCBS055ULH |
| Blueprint | [ENCFF601LYG](https://www.encodeproject.org/files/ENCFF601LYG/) | left renal pelvis | DNase1-seq | R_ENCBS055ULH |
| Blueprint | [ENCFF544DPT](https://www.encodeproject.org/files/ENCFF544DPT/) | left renal pelvis | DNase1-seq | R_ENCBS055ULH |
| Blueprint | [ENCFF946SSJ](https://www.encodeproject.org/files/ENCFF946SSJ/) | left renal pelvis | DNase1-seq | R_ENCBS257BTU |
| Blueprint | [ENCFF657QAX](https://www.encodeproject.org/files/ENCFF657QAX/) | left renal pelvis | DNase1-seq | R_ENCBS257BTU |
| Blueprint | [ENCFF639WUS](https://www.encodeproject.org/files/ENCFF639WUS/) | left renal pelvis | DNase1-seq | R_ENCBS257BTU |
| Blueprint | [ENCFF431EYP](https://www.encodeproject.org/files/ENCFF431EYP/) | left renal pelvis | DNase1-seq | R_ENCBS226ZND |
| Blueprint | [ENCFF065NFR](https://www.encodeproject.org/files/ENCFF065NFR/) | left renal pelvis | DNase1-seq | R_ENCBS226ZND |
| Blueprint | [ENCFF767VEQ](https://www.encodeproject.org/files/ENCFF767VEQ/) | left renal pelvis | DNase1-seq | R_ENCBS226ZND |
| Blueprint | [ENCFF771XJT](https://www.encodeproject.org/files/ENCFF771XJT/) | left renal pelvis | DNase1-seq | R_ENCBS226ZND |
| Blueprint | [ENCFF768GNX](https://www.encodeproject.org/files/ENCFF768GNX/) | left renal pelvis | DNase1-seq | R_ENCBS754ANY |
| Blueprint | [ENCFF061DNQ](https://www.encodeproject.org/files/ENCFF061DNQ/) | left renal pelvis | DNase1-seq | R_ENCBS754ANY |
| Blueprint | [ENCFF806MUH](https://www.encodeproject.org/files/ENCFF806MUH/) | left renal pelvis | DNase1-seq | R_ENCBS754ANY |
| Blueprint | [ENCFF052JNW](https://www.encodeproject.org/files/ENCFF052JNW/) | left renal pelvis | DNase1-seq | R_ENCBS754ANY |
| Blueprint | [ENCFF909ULM](https://www.encodeproject.org/files/ENCFF909ULM/) | left renal pelvis | DNase1-seq | R_ENCBS754ANY |
| Blueprint | [ENCFF770YNO](https://www.encodeproject.org/files/ENCFF770YNO/) | left renal pelvis | DNase1-seq | R_ENCBS754ANY |
| Blueprint | [ENCFF445HLC](https://www.encodeproject.org/files/ENCFF445HLC/) | left renal pelvis | DNase1-seq | R_ENCBS754ANY |
| Blueprint | [ENCFF027YDE](https://www.encodeproject.org/files/ENCFF027YDE/) | right renal pelvis | DNase1-seq | R_ENCBS145EYH |
| Blueprint | [ENCFF423KHY](https://www.encodeproject.org/files/ENCFF423KHY/) | right renal pelvis | DNase1-seq | R_ENCBS145EYH |
| Blueprint | [ENCFF245ELJ](https://www.encodeproject.org/files/ENCFF245ELJ/) | right renal pelvis | DNase1-seq | R_ENCBS935VKR |
| Blueprint | [ENCFF863PCQ](https://www.encodeproject.org/files/ENCFF863PCQ/) | right renal pelvis | DNase1-seq | R_ENCBS935VKR |
| Blueprint | [ENCFF009UUH](https://www.encodeproject.org/files/ENCFF009UUH/) | right renal pelvis | DNase1-seq | R_ENCBS935VKR |
| Blueprint | [ENCFF530YSE](https://www.encodeproject.org/files/ENCFF530YSE/) | right renal pelvis | DNase1-seq | R_ENCBS855RFN |
| Blueprint | [ENCFF892SRC](https://www.encodeproject.org/files/ENCFF892SRC/) | right renal pelvis | DNase1-seq | R_ENCBS855RFN |
| Blueprint | [ENCFF889RBT](https://www.encodeproject.org/files/ENCFF889RBT/) | right renal pelvis | DNase1-seq | R_ENCBS855RFN |
| Blueprint | [ENCFF974YUN](https://www.encodeproject.org/files/ENCFF974YUN/) | right renal pelvis | DNase1-seq | R_ENCBS827OFK |
| Blueprint | [ENCFF795TJS](https://www.encodeproject.org/files/ENCFF795TJS/) | right renal pelvis | DNase1-seq | R_ENCBS827OFK |
| Blueprint | [ENCFF138BJK](https://www.encodeproject.org/files/ENCFF138BJK/) | right renal pelvis | DNase1-seq | R_ENCBS827OFK |
| Blueprint | [ENCFF733MIZ](https://www.encodeproject.org/files/ENCFF733MIZ/) | right renal pelvis | DNase1-seq | R_ENCBS827OFK |
| Blueprint | [ENCFF554OEC](https://www.encodeproject.org/files/ENCFF554OEC/) | right renal pelvis | DNase1-seq | R_ENCBS827OFK |
| Blueprint | [ENCFF719GVR](https://www.encodeproject.org/files/ENCFF719GVR/) | right renal pelvis | DNase1-seq | R_ENCBS827OFK |
| Blueprint | [ENCFF694DBL](https://www.encodeproject.org/files/ENCFF694DBL/) | right renal pelvis | DNase1-seq | R_ENCBS827OFK |
| Blueprint | [ENCFF260GSJ](https://www.encodeproject.org/files/ENCFF260GSJ/) | spinal cord | DNase1-seq | R_ENCBS373RUA |
| Blueprint | [ENCFF369QCM](https://www.encodeproject.org/files/ENCFF369QCM/) | spinal cord | DNase1-seq | R_ENCBS373RUA |
| Blueprint | [ENCFF222EIH](https://www.encodeproject.org/files/ENCFF222EIH/) | spinal cord | DNase1-seq | R_ENCBS300UPT |
| Blueprint | [ENCFF854UHG](https://www.encodeproject.org/files/ENCFF854UHG/) | spinal cord | DNase1-seq | R_ENCBS300UPT |
| Blueprint | [ENCFF073KRY](https://www.encodeproject.org/files/ENCFF073KRY/) | right renal cortex interstitium | DNase1-seq | R_ENCBS100DZU |
| Blueprint | [ENCFF709HXO](https://www.encodeproject.org/files/ENCFF709HXO/) | right renal cortex interstitium | DNase1-seq | R_ENCBS100DZU |
| Blueprint | [ENCFF102SFO](https://www.encodeproject.org/files/ENCFF102SFO/) | right renal cortex interstitium | DNase1-seq | R_ENCBS100DZU |
| Blueprint | [ENCFF391HSJ](https://www.encodeproject.org/files/ENCFF391HSJ/) | right renal cortex interstitium | DNase1-seq | R_ENCBS183TBX |
| Blueprint | [ENCFF855JXV](https://www.encodeproject.org/files/ENCFF855JXV/) | right renal cortex interstitium | DNase1-seq | R_ENCBS183TBX |
| Blueprint | [ENCFF934XHE](https://www.encodeproject.org/files/ENCFF934XHE/) | right renal cortex interstitium | DNase1-seq | R_ENCBS818WCN |
| Blueprint | [ENCFF455IUH](https://www.encodeproject.org/files/ENCFF455IUH/) | right renal cortex interstitium | DNase1-seq | R_ENCBS818WCN |
| Blueprint | [ENCFF863QQW](https://www.encodeproject.org/files/ENCFF863QQW/) | right renal cortex interstitium | DNase1-seq | R_ENCBS818WCN |
| Blueprint | [ENCFF948AVM](https://www.encodeproject.org/files/ENCFF948AVM/) | right renal cortex interstitium | DNase1-seq | R_ENCBS818WCN |
| Blueprint | [ENCFF882KUN](https://www.encodeproject.org/files/ENCFF882KUN/) | right renal cortex interstitium | DNase1-seq | R_ENCBS818WCN |
| Blueprint | [ENCFF705NSS](https://www.encodeproject.org/files/ENCFF705NSS/) | right renal cortex interstitium | DNase1-seq | R_ENCBS818WCN |
| Blueprint | [ENCFF838LOC](https://www.encodeproject.org/files/ENCFF838LOC/) | right renal cortex interstitium | DNase1-seq | R_ENCBS818WCN |
| Blueprint | [ENCFF422YYG](https://www.encodeproject.org/files/ENCFF422YYG/) | spleen | DNase1-seq | R_ENCBS599HUG |
| Blueprint | [ENCFF525TWK](https://www.encodeproject.org/files/ENCFF525TWK/) | psoas muscle | DNase1-seq | R_ENCBS008QPC |
| Blueprint | [ENCFF457YXQ](https://www.encodeproject.org/files/ENCFF457YXQ/) | psoas muscle | DNase1-seq | R_ENCBS008QPC |
| Blueprint | [ENCFF940GXC](https://www.encodeproject.org/files/ENCFF940GXC/) | psoas muscle | DNase1-seq | R_ENCBS008QPC |
| Blueprint | [ENCFF136KBO](https://www.encodeproject.org/files/ENCFF136KBO/) | psoas muscle | DNase1-seq | R_ENCBS008QPC |
| Blueprint | [ENCFF065TLM](https://www.encodeproject.org/files/ENCFF065TLM/) | muscle of trunk | DNase1-seq | R_ENCBS992XAC |
| Blueprint | [ENCFF979HOG](https://www.encodeproject.org/files/ENCFF979HOG/) | ovary | DNase1-seq | R_ENCBS645JEU |
| Blueprint | [ENCFF659QQK](https://www.encodeproject.org/files/ENCFF659QQK/) | ovary | DNase1-seq | R_ENCBS645JEU |
| Blueprint | [ENCFF827YHI](https://www.encodeproject.org/files/ENCFF827YHI/) | ovary | DNase1-seq | R_ENCBS341OKA |
| Blueprint | [ENCFF331JVF](https://www.encodeproject.org/files/ENCFF331JVF/) | ovary | DNase1-seq | R_ENCBS341OKA |
| Blueprint | [ENCFF780LKZ](https://www.encodeproject.org/files/ENCFF780LKZ/) | pancreas | DNase1-seq | R_ENCBS507RPJ |
| Blueprint | [ENCFF890UCG](https://www.encodeproject.org/files/ENCFF890UCG/) | pancreas | DNase1-seq | R_ENCBS507RPJ |
| Blueprint | [ENCFF985FNS](https://www.encodeproject.org/files/ENCFF985FNS/) | pancreas | DNase1-seq | R_ENCBS914JTX |
| Blueprint | [ENCFF457ONU](https://www.encodeproject.org/files/ENCFF457ONU/) | pancreas | DNase1-seq | R_ENCBS914JTX |
| Blueprint | [ENCFF109QYQ](https://www.encodeproject.org/files/ENCFF109QYQ/) | testis | DNase1-seq | R_ENCBS796DWQ |
| Blueprint | [ENCFF025KGA](https://www.encodeproject.org/files/ENCFF025KGA/) | testis | DNase1-seq | R_ENCBS796DWQ |
| Blueprint | [ENCFF249RTG](https://www.encodeproject.org/files/ENCFF249RTG/) | forelimb muscle | DNase1-seq | R_ENCBS988KQJ |
| Blueprint | [ENCFF949DLJ](https://www.encodeproject.org/files/ENCFF949DLJ/) | hindlimb muscle | DNase1-seq | R_ENCBS105LQM |
| Blueprint | [ENCFF135DMC](https://www.encodeproject.org/files/ENCFF135DMC/) | hindlimb muscle | DNase1-seq | R_ENCBS105LQM |
| Blueprint | [ENCFF256YZN](https://www.encodeproject.org/files/ENCFF256YZN/) | H1-hESC | DNase1-seq | R_ENCBS559QNR_ENCBS568FYY_ENCBS945MCY |
| Blueprint | [ENCFF217JMU](https://www.encodeproject.org/files/ENCFF217JMU/) | H1-hESC | DNase1-seq | R_ENCBS559QNR_ENCBS568FYY_ENCBS945MCY |
| Blueprint | [ENCFF390EVX](https://www.encodeproject.org/files/ENCFF390EVX/) | H1-hESC | DNase1-seq | R_ENCBS559QNR_ENCBS568FYY_ENCBS945MCY |
| Blueprint | C0011IH1.ERX197160.H3K27ac.bwa.GRCh38.20150528.bw | CD14-positive, CD16-negative classical monocyte | H3K27ac | C0011IH1 |
| Blueprint | S00C0JH1.ERX547233.H3K27ac.bwa.GRCh38.20150528.bw | macrophage_-_T_6days_B-glucan | H3K27ac | S00C0JH1 |
| Blueprint | S00XUNH1.ERX941029.H3K27ac.bwa.GRCh38.20150529.bw | Acute Myeloid Leukemia | H3K27ac | S00XUNH1 |
| Blueprint | C0010KH1.ERX197174.H3K27ac.bwa.GRCh38.20150528.bw | CD14-positive, CD16-negative classical monocyte | H3K27ac | C0010KH1 |

**Supplementary Table S1:** Data IDs of the raw data files processed in this study.

#### Data preprocessing

We conducted two main data preprocessing steps: (1) quantification of gene-expression and (2) alignment of DNase1-seq data and identification of DNase hypersensitive sites.

The essential commands used for both steps along with software versions are presented in the next two sections.

##### RNA-seq processing

Gene-expression was quantified using Salmon [5], version 0.8.2., the Gencode [6] transcript index v26, and the Gencode genome annotation v26.

For single end reads, we used the command:

***./salmon quant -i gencode.v26.transcripts.index/ -l A -r <Sample>_R1.fastq.gz -p 12 -o quants/<Sample> --seqBias --gcBias -g gencode.v26.annotation.gtf***

For paired end reads, we used the command:

***./salmon quant -i gencode.v26.transcripts.index/ -l A -1 <Sample>_R1.fastq.gz -2 <Sample>_R2.fastq.gz -p 12 -o quants/<Sample> --seqBias --gcBias -g gencode.v26.annotation.gtf***

Discretized gene-expression values were computed using the POE method, see Supplementary Sec. 4 for details.

##### DNase1-seq processing

DNase1-seq reads were aligned to the hg38 reference genome with bowtie [7], version 1.2.1.1, and samtools [8] version 1.2. For single end reads, this was done using the command call:

***./bowtie --threads 10 -S GRCh38_no_alt/GCA_000001405.15_GRCh38_no_alt_analysis_set <Sample>_R1.fastq.gz | samtools view -b -o <Sample>.bam -) 2> Sample>.bowtie_statistics.txt***

and for paired end reads using:

***./bowtie --threads 10 -S GRCh38_no_alt/GCA_000001405.15_GRCh38_no_alt_analysis_set -1 <Sample>_R1.fastq.gz -2 <Sample>_R2.fastq.gz | samtools view -b -o <Sample>.bam -) 2 > <Sample> .bowtie_statistics.txt***

In case that multiple fasta files exist for one sample, they were concatenated using the linux ***cat*** command prior to alignment, if applicable, separately per strand.

DNase Hypersensitive Sites (DHS) were identified with JAMM [9], version 1.0.7.5 and samtools version 1.2.

For single end reads, we use the commands:

***bedtools bamtobed -i <Sample>.bam > JAMM-Input/<Sample>.bed***

***bash JAMM.sh –s JAMM-Input -g hg38_chrSize.txt –o <Sample>_peaks -f 1 -p 8***

***rm –r JAMM-Input/***

For paired end reads, we run the commands:

***samtools sort -n -O bam -@10 -T Bam-Sort-Pre <Sample>.bam | samtools view -bf 0x2 - | bedtools bamtobed -bedpe -i stdin > JAMM-Input/<Sample>.bed***

***bash JAMM.sh -s JAMM-Input -g hg38_chrSize.txt -o <Sample>_peaks -f 1 -p 8 -t paired***

***rm –r JAMM-Input/***

As described in the JAMM github, we computed the enrichment of DNase1-seq signal within the peak by dividing column 7 by column 9 of the JAMM output files:

***awk '{print $1\"\t\"$2\"\t\"$3\"\t\"$7/$9}' <Sample>/peaks/filtered.peaks.narrowPeak > <Sample>/peaks/filtered.peaks.narrowPeak.adoptedCoverage***

Bigwig files were generated using the DEEPtools [10] bamCoverage program, version 2.4.2 with the command

***bamCoverage -b <Sample>.bam.sorted.bam –o <Sample>.bw -p 4 –normalizeUsingRPKM***

### Supplementary Section 2: STITCHIT algorithm

#### Running the STITCHIT algorithm

To run STITCHIT, the user needs to provide discretized (-d) and original expression data (-o), a gene annotation file (-a), a chromosome size file (-s), as well as big wig files with the epigenetic signal to consider (-b). Using the command:

**./build/core/STITCHIT –b <Consortium>/DNase_bw/ -a ../../../nobackup/References/gencode.v26.annotation.gtf -d <Consortium>/_Discretised_Complet.txt -o <Consortium>_expression.txt  -s data/hg38_chrSize.txt -w 25000 -c 12 -p 0.05 -g <geneID> -z 10 -f ../../../archive00/Segmentation_<Consortium>/ -r 500000 -t 2000**

a call to STITCHIT can be invoked. The parameter –w denotes the size of the window extension up an downstream of the gene, -c denotes the number of used CPU, -p is the significance threshold for the correlation test, -g is the parameter to denote the target gene ID, -z indicates the width of the initial binning, -f denotes the output path, -r is the maximum size of the entire search region and –t refers to the maximum size of a segment. Further details on the parameters are provided in the repository (www.github.com/Schulzlab/Stitchit).

#### General Setup of the Data Matrix processed in Steps (c) to (f) in Figure S1

The extracted the data matrix $D_{i,j}$ consists of $m$ rows corresponding to the samples and $n$ columns that correspond to the base pair positions. In addition, each sample is associated with a class label according to its expression value (see Figure S1c). With $C_{k}$ we relate to all rows that are assigned to label $k$. We initially group the base pairs into segments of size $\beta$, which is a user defined parameter that we set to set $\beta=5$ in our experiments. Setting $\beta$ to a larger value would speed up the runtime, however, since no segment can be smaller than $\beta$, increasing it could lead to missing out segments.

#### Example

The following example showcases a run of the STITCH algorithm on a data matrix consisting of six samples with six initial segments ($\beta=1$). The example tries to find a segmentation that best explains the expression difference for two classes, labelled 1 and 2. Each class has been assigned to three samples.

Data matrix $D_{i,j}$

|  | **b1** | **b2** | **b3** | **b4** | **b5** | **b6** | **exp** |
| --- | --- | --- | --- | --- | --- | --- | --- |
| **s1** | 10 | 9 | 1 | 1 | 7 | 8 | 1 |
| **s2** | 11 | 10 | 2 | 1 | 7 | 8 | 1 |
| **s3** | 10 | 8 | 1 | 0 | 9 | 8 | 1 |
| **s4** | 2 | 4 | 25 | 24 | 0 | 1 | 2 |
| **s5** | 3 | 5 | 23 | 22 | 1 | 0 | 2 |
| **s6** | 2 | 6 | 26 | 25 | 0 | 1 | 2 |

From the data matrix, we compute the sum $S$ and sum of squares of the values of each column of $D$ per class.

$$S_{k}\left[ j \right]=S\left[ j-1 \right]+\sum_{i\in C_{k}} d_{i,j}$$

$${SS}_{k}\left[ j \right]=SS\left[ j-1 \right]+\sum_{i\in C_{k}} d_{i,j}^{2}$$

Sum per class $S_{k}:$

| **S_1_1** | **S_1_2** | **S_1_3** | **S_1_4** | **S_1_5** | **S_1_6** |
| --- | --- | --- | --- | --- | --- |
| 31 | 58 | 62 | 64 | 87 | 111 |

| **S_2_1** | **S_2_2** | **S_2_3** | **S_2_4** | **S_2_5** | **S_2_6** |
| --- | --- | --- | --- | --- | --- |
| 7 | 22 | 96 | 167 | 168 | 170 |

Sum of squares per class ${SS}_{k}$

| **SS_1_1** | **SS_1_2** | **SS_1_3** | **SS_1_4** | **SS_1_5** | **SS_1_6** |
| --- | --- | --- | --- | --- | --- |
| 321 | 566 | 572 | 574 | 753 | 945 |

| **SS_2_1** | **SS_2_2** | **SS_2_3** | **SS_2_4** | **SS_2_5** | **SS_2_6** |
| --- | --- | --- | --- | --- | --- |
| 17 | 94 | 1924 | 3609 | 3610 | 3612 |

To calculate the data costs $w_{i,j}^{k}$ for each hypothetical segment from position *i* to *j*, with $i\leq j$, we need to compute the empirical standard deviation $\hat{\sigma}_{i,j}^{k}$ as well as the number of data points within a bin$n_{i,j}^{k}$. This we can do in constant time using the precomputed vectors $S_{k}$ and ${SS}_{k}$ [15], where

$\hat{\sigma}_{i,j}^{k}= \sqrt{\frac{1}{n_{i,j}^{k}} \left( \left( SS_{k}\left[ j \right]-SS_{k}[i-1] \right)-\frac{1}{n_{i,j}^{k}}\left( S_{k}\left[ j \right]-S_{k}[i-1] \right)^{2} \right)}$,

$n_{i,j}^{k}=\left( j+1-i \right)*\left| C_{k} \right|$,

where $\left| C_{k} \right|$ denotes the number of samples related to class$k$.

Data points per segment $n_{i,j}$

| **i, j** | **1** | **2** | **3** | **4** | **5** | **6** |
| --- | --- | --- | --- | --- | --- | --- |
| **1** | 3 | 6 | 9 | 12 | 15 | 18 |
| **2** | 0 | 3 | 6 | 9 | 12 | 15 |
| **3** | 0 | 0 | 3 | 6 | 9 | 12 |
| **4** | 0 | 0 | 0 | 3 | 6 | 9 |
| **5** | 0 | 0 | 0 | 0 | 3 | 6 |
| **6** | 0 | 0 | 0 | 0 | 0 | 3 |
| The matrix $n_{i,j}$ is identical for both classes. | | | | | | |

Standard deviation class 1 $\hat{\sigma}_{i,j}^{1}$

| **i, j** | **1** | **2** | **3** | **4** | **5** | **6** |
| --- | --- | --- | --- | --- | --- | --- |
| **1** | 0.471405 | 0.942809 | 4.012327 | 4.403282 | 4.069398 | 3.804237 |
| **2** | 0 | 0.816497 | 3.890873 | 3.829708 | 3.771236 | 3.627059 |
| **3** | 0 | 0 | 0.471405 | 0.57735 | 3.224137 | 3.47511 |
| **4** | 0 | 0 | 0 | 0.471405 | 3.578485 | 3.435472 |
| **5** | 0 | 0 | 0 | 0 | 0.942809 | 0.687184 |
| **6** | 0 | 0 | 0 | 0 | 0 | 0 |

Standard deviation class 2 $\hat{\sigma}_{i,j}^{2}$

| **i,j** | **1** | **2** | **3** | **4** | **5** | **6** |
| --- | --- | --- | --- | --- | --- | --- |
| **1** | 0.471405 | 1.490712 | 10 | 10.34777 | 10.73437 | 10.55789 |
| **2** | 0 | 0.816497 | 9.889669 | 9.113821 | 10.92748 | 11.02643 |
| **3** | 0 | 0 | 1.247219 | 1.34371 | 11.29186 | 11.87668 |
| **4** | 0 | 0 | 0 | 1.247219 | 11.7047 | 10.9522 |
| **5** | 0 | 0 | 0 | 0 | 0.471405 | 0.5 |
| **6** | 0 | 0 | 0 | 0 | 0 | 0.471405 |

From $\hat{\sigma}_{i,j}^{k}$ and$n_{i,j}^{k}$, we then compute the data costs $w_{i,j}^{k}$ according to the MDL encoding for Gaussian distributed data points [16]. For notational convenience, we write $s$ to denote to the segment according to the range $i,j$. Hence, we compute the data costs for segment $s$ and class label $k$ as

$$L\left( D | s,k \right)=w_{i,j}^{k}= \frac{n_{i,j}^{k}}{2}\left( \frac{1}{log(2)}+\log_{2} (2\pi\hat{\sigma}_{i,j}^{k}\hat{\sigma}_{i,j}^{k}) \right)+n_{i,j}^{k}\log_{2} \tau,$$

Where $\tau$ is the data resolution. To simplify the example, we set $\tau=1$. As we only work with integers setting $\tau=1$ is sufficient, however, since we later on also encode the mean values for each segment, which can be a floating point number, we set $\tau=0.1$ in all our experiments.

Scores for class 1 $w_{i,j}^{1}$

| **i,j** | **1** | **2** | **3** | **4** | **5** | **6** |
| --- | --- | --- | --- | --- | --- | --- |
| **1** | 5.705249 | 17.4105 | 44.92036 | 61.50349 | 75.17291 | 88.45775 |
| **2** | 0 | 8.082693 | 29.68083 | 44.31552 | 58.821 | 72.68268 |
| **3** | 0 | 0 | 5.705249 | 13.16539 | 42.08063 | 57.40525 |
| **4** | 0 | 0 | 0 | 5.705249 | 28.95637 | 42.90498 |
| **5** | 0 | 0 | 0 | 0 | 8.705249 | 14.67289 |
| **6** | 0 | 0 | 0 | 0 | 0 | 0 |

Scores for class 2 $w_{i,j}^{2}$

| **i,j** | **1** | **2** | **3** | **4** | **5** | **6** |
| --- | --- | --- | --- | --- | --- | --- |
| **1** | 5.705249 | 21.37628 | 56.77776 | 76.29552 | 96.16316 | 114.9653 |
| **2** | 0 | 8.082693 | 37.75581 | 55.57291 | 77.2392 | 96.7441 |
| **3** | 0 | 0 | 9.916281 | 20.4776 | 58.35531 | 78.68126 |
| **4** | 0 | 0 | 0 | 9.916281 | 39.21437 | 57.95874 |
| **5** | 0 | 0 | 0 | 0 | 5.705249 | 11.92027 |
| **6** | 0 | 0 | 0 | 0 | 0 | 5.705249 |

Next, we compute the total cost matrix $W$, defined as

$W_{i,j}=\sum_{l=1..k} \frac{w_{i,j}^{k}}{\left| C_{k} \right|}$,

Total cost matrix *W:*

| **i,j** | **1** | **2** | **3** | **4** | **5** | **6** |
| --- | --- | --- | --- | --- | --- | --- |
| **1** | 3.803499 | 12.92893 | 33.89937 | 45.93301 | 57.11203 | 67.80769 |
| **2** | 0 | 5.388462 | 22.47888 | 33.29614 | 45.3534 | 56.47559 |
| **3** | 0 | 0 | 5.207177 | 11.21433 | 33.47865 | 45.36217 |
| **4** | 0 | 0 | 0 | 5.207177 | 22.72358 | 33.62124 |
| **5** | 0 | 0 | 0 | 0 | 4.803499 | 8.864386 |
| **6** | 0 | 0 | 0 | 0 | 0 | 1.90175 |

Using $W$, we conduct the dynamic programming approach as described in [17] to find the optimal segmentation for each possible number of segments according to our cost function, following the recursion formula

$$c_{i,j}= \min_{k=1,\ldots,j-1} (c_{i-1,k}+W_{k+1,j})$$

| **i,j** | **1** | **2** | **3** | **4** | **5** | **6** |
| --- | --- | --- | --- | --- | --- | --- |
| **1** | 3.803499 | 12.92893 | 33.89937 | 45.93301 | 57.11203 | 67.80769 |
| **2** | 0 | 9.191961 | 18.1361 | 24.14326 | 46.40757 | 54.79739 |
| **3** | 0 | 0 | 14.39914 | 20.40629 | 28.94675 | 33.00764 |
| **4** | 0 | 0 | 0 | 19.60631 | 25.20979 | 29.27068 |
| **5** | 0 | 0 | 0 | 0 | 24.40981 | 28.4707 |
| **6** | 0 | 0 | 0 | 0 | 0 | 26.31156 |

Subsequently, we compute the costs for the models of a segmentation $S$ consisting of $|S|$ segments according to:

$$L\left( S \right)=L_{\mathbb{N}}\left( \left| S \right| \right)+\left| C \right|\left| S \right|{log}_{2}\left( \frac{|\max-\min|}{\tau} \right)+{log}_{2}\left( \begin{matrix} n-1 \\ \left| S \right|-1 \end{matrix} \right),$$

Where $\left| C \right|$ is the number of possible class labels (in this example $\left| C \right|=2$).

Costs for the binning:

| **Level 1** | 10.91945 |
| --- | --- |
| **Level 2** | 23.64225 |
| **Level 3** | 35.29254 |
| **Level 4** | 45.44401 |
| **Level 5** | 54.66348 |
| **Level 6** | 62.33327 |

Combined with the last column of the dynamic programing matrix this yields the final costs:

| **Level 1** | 78.72714 |
| --- | --- |
| **Level 2** | 78.43965 |
| **Level 3** | **68.30019** |
| **Level 4** | 74.71469 |
| **Level 5** | 83.13418 |
| **Level 6** | 88.64484 |

Here, the minimum cost is associated to the third level.

Performing backtracking in the dynamic programming matrix yields the segmentation

$$b_{1,2},b_{3,4},b_{5,6}.$$

The corresponding points in the matrix are marked in red.

| **i,j** | **1** | **2** | **3** | **4** | **5** | **6** |
| --- | --- | --- | --- | --- | --- | --- |
| **1** | 3.803499 | **12.92893** | 33.89937 | 45.93301 | 57.11203 | 67.80769 |
| **2** | 0 | 9.191961 | 18.1361 | **24.14326** | 46.40757 | 54.79739 |
| **3** | 0 | 0 | 14.39914 | 20.40629 | 28.94675 | **33.00764** |
| **4** | 0 | 0 | 0 | 19.60631 | 25.20979 | 29.27068 |
| **5** | 0 | 0 | 0 | 0 | 24.40981 | 28.4707 |
| **6** | 0 | 0 | 0 | 0 | 0 | 26.31156 |

### Supplementary Section 3: Discretize expression data

We use the Probability Of Expression (POE)[11] algorithm to discretize the gene-expression data. Due to R computability and versioning issues, we had to rewrite some parts of the code. The entire R code is shown below. Note that the implementation of POE is also not designed to treat whole genome datasets. Therefore, we consider the data in batches, i.e. a subset of genes is discretized one after another.

library("metaArray")
library("Biobase")

args=commandArgs(TRUE)

myPOE<-function (AA, NN = NULL, id = NULL, M = 2000, kap.min = 3, logdata = FALSE,
                 stepsize = 0.5, centersample = FALSE, centergene = FALSE,
                 generatestarts = TRUE, start.method = 1, startobject = R0,
                 collapse.to.two = FALSE, burnin = 200, collapse.window = 50,
                 converge.threshold = 0.01, PR = list(alpha.mm = 0, alpha.sd = 100,
                                                      mu.mm = 0, mu.sd = 100, pipos.mm = 0, pipos.sd = 100,
                                                      pineg.mm = 0, pineg.sd = 100, kap.pri.rate = 1, tausqinv.aa = 1,
                                                      tausqinv.bb = 0.1))

{
  TT <- ncol(AA)
  GG <- nrow(AA)
  tmp.poemat <- matrix(0, GG, TT)
  if (!is.null(NN) & length(NN) != TT) {
    stop("Length of NN should be the same as ncol(AA)")
  }
  if (is.na(sum(as.vector(AA))))
    stop("NA values not allowed in AA matrix. Remove rows with NA values and retry.")
  if (logdata == TRUE)
    AA <- log(AA)
  if (is.null(id) == TRUE)
    id <- as.character(1:GG)
  if (is.null(NN))
    NN <- rep(0, TT)
  if (centersample) {
    sammed <- apply(AA, 2, median)
    AA <- sweep(AA, 2, sammed)
  }                                                                                                                                                                                                                                                                                                                                                                                       
  if (centergene) {
    genmed <- apply(AA, 1, median)
    AA <- sweep(AA, 1, genmed)
  }
  if (generatestarts == TRUE) {
    alpha0 <- apply(AA, 2, mean)
    mats <- t(apply(AA, 1, sort))
    mug0 <- sigmag0 <- kappaposg0 <- kappanegg0 <- rep(NA,
                                                       GG)
    piposg0 <- pinegg0 <- rep(NA, GG)
    if (start.method == 1) {
      for (gg in 1:GG) {
        mug0[gg] <- mean(mats[gg, -c(1, 2, TT - 1, TT)])
        pinegg0[gg] <- piposg0[gg] <- 2/TT
        sigmag0[gg] <- sqrt(var(mats[gg, -c(1, 2, TT -
                                              1, TT)]))
        kappaposg0[gg] <- max(kap.min * sigmag0[gg],
                              mug0[gg] - mats[gg, 1])
        kappanegg0[gg] <- max(kap.min * sigmag0[gg],
                              mats[gg, TT] - mug0[gg])
      }
    }
    if (start.method == 2) {
      for (gg in 1:GG) {
        clu <- kmeans(mats[gg, ], 2)
        if (clu$size[1] == clu$size[2]) {
          big <- 1
          sma <- 2
        }
        else {
          big <- (1:2)[clu$size == max(clu$size)]
          sma <- (1:2)[clu$size == min(clu$size)]
        }
        mug0[gg] <- clu$centers[big]
        sigmag0[gg] <- sqrt(var(mats[gg, clu$cluster ==
                                       big]))
        if (mug0[gg] > 0) {
          pinegg0[gg] <- clu$size[sma]/TT
          piposg0[gg] <- 1/TT
        }
        else {
          piposg0[gg] <- clu$size[sma]/TT
          pinegg0[gg] <- 1/TT
        }
        kappaposg0[gg] <- max(max(mats[gg, ]) - mug0[gg],
                              kap.min * sigmag0[gg])
        kappanegg0[gg] <- max(mug0[gg] - min(mats[gg,
                                                  ]), kap.min * sigmag0[gg])
      }
    }
    if (start.method == 3) {
      for (gg in 1:GG) {
        clu <- kmeans(mats[gg, ], 3)
        posc <- (1:3)[clu$centers == max(clu$centers)]
        negc <- (1:3)[clu$centers == min(clu$centers)]
        midc <- (1:3)[clu$centers == median(clu$centers)]
        mug0[gg] <- clu$centers[midc]
        sigmag0[gg] <- sqrt(var(mats[gg, clu$cluster ==
                                       midc]))
        pinegg0[gg] <- clu$size[negc]/TT
        piposg0[gg] <- clu$size[posc]/TT
        kappaposg0[gg] <- max(max(mats[gg, ]) - mug0[gg],
                              kap.min * sigmag0[gg])
        kappanegg0[gg] <- max(mug0[gg] - min(mats[gg,
                                                  ]), kap.min * sigmag0[gg])
      }
    }
    if (start.method == 4) {
      if (sum(NN) < 3)
        stop("start.method==4 requires at least 3 normal samples")
      for (gg in 1:GG) {
        mug0[gg] <- mean(AA[gg, NN == 1])
        sigmag0[gg] <- sqrt(var(AA[gg, NN == 1]))
        clu <- kmeans(AA[gg, ], 2)
        norm <- (1:2)[abs(clu$centers - mug0[gg]) ==
                        min(abs(clu$centers - mug0[gg]))]
  expr <- 1
        if (norm == 1)
          expr <- 2
        if (clu$centers[norm] > clu$centers[expr]) {
          pinegg0[gg] <- min(clu$size[expr]/TT, 1/2)
          piposg0[gg] <- 1/TT
        }
        else {
          piposg0[gg] <- min(clu$size[expr]/TT, 1/2)
          pinegg0[gg] <- 1/TT
        }
        kappaposg0[gg] <- max(max(AA[gg, ]) - mug0[gg],
                              kap.min * sigmag0[gg])
        kappanegg0[gg] <- max(mug0[gg] - min(AA[gg, ]),
                              kap.min * sigmag0[gg])
      }
    }
    sigmagsqinv0 <- 1/sigmag0^2
    gamma0 <- (mean(sigmagsqinv0))^2/var(sigmagsqinv0)
    lambda0 <- mean(sigmagsqinv0)/var(sigmagsqinv0)
    tmpmat <- matrix(0, GG, TT)

    PP <- list(alpha = alpha0, mug = mug0, kappaposg = kappaposg0,
               kappanegg = kappanegg0, sigmag = sigmag0, piposg = piposg0,
               pinegg = pinegg0, mu = mean(mug0), tausqinv = 1/var(mug0),
               gamma = gamma0, lambda = lambda0, pil.pos.mean = mean(logit(piposg0)),
               pil.neg.mean = mean(logit(pinegg0)), pil.pos.prec = mean(piposg0) *
                 (1 - mean(piposg0)), pil.neg.prec = mean(pinegg0) *
                 (1 - mean(pinegg0)), kap.pos.rate = 1/mean(kappaposg0),
               kap.neg.rate = 1/mean(kappanegg0), poe = tmp.poemat,
               phat.pos = tmpmat, phat.neg = tmpmat, accept = 0)
  }
  if (generatestarts == FALSE) {
    mmm <- length(startobject$gamma)
    PP <- list(alpha = startobject$alpha[mmm, ], mug = startobject$mug[mmm,
                                                                       ], kappaposg = startobject$kappaposg[mmm, ], kappanegg = startobject$kappanegg[mmm,
                                                                                                                                                      ], sigmag = startobject$sigmag[mmm, ], piposg = startobject$piposg[mmm,
                                                                                                                                                                                                                         ], pinegg = startobject$pinegg[mmm, ], mu = startobject$mu[mmm],
               tausqinv = startobject$tausqinv[mmm], gamma = startobject$gamma[mmm],
               lambda = startobject$lambda[mmm], pil.pos.mean = startobject$pil.pos.mean[mmm],
               pil.pos.prec = startobject$pil.pos.prec[mmm], pil.neg.mean = startobject$pil.neg.mean[mmm],
               pil.neg.prec = startobject$pil.neg.prec[mmm], kap.pos.rate = startobject$kap.pos.rate[mmm],
               kap.neg.rate = startobject$kap.neg.rate[mmm], poe = tmp.poemat,
               phat.pos = tmpmat, phat.neg = tmpmat, accept = 0)
  }
  if (collapse.to.two) {
    cat("Collapsing to Two: ")
    cw <- collapse.window
    mm <- 0
    pidif <- 1
    pim.pos <- pim.neg <- 100
    while (pidif > converge.threshold) {
      mm <- mm + 1
      new.PR <- unlist(PR)
      new.PP <- unlist(PP)
      avgpos <- rep(0, GG * (3 * TT + 6) + TT + 11)
      res <- .C("poe_fit_2", as.matrix(AA), as.integer(NN),
                as.double(new.PR), as.double(new.PP), as.integer(GG),
                as.integer(TT), as.integer(1), avgpos = as.double(avgpos),
                PACKAGE = "metaArray")$avgpos
      PP$alpha <- res[1:TT]
      PP$mug <- res[(TT + 1):(TT + GG)]
      PP$kappaposg <- res[(TT + GG + 1):(TT + 2 * GG)]
      PP$kappanegg <- res[(TT + 2 * GG + 1):(TT + 3 * GG)]
      PP$sigmag <- res[(TT + 3 * GG + 1):(TT + 4 * GG)]
      PP$piposg <- res[(TT + 4 * GG + 1):(TT + 5 * GG)]
      PP$pinegg <- res[(TT + 5 * GG + 1):(TT + 6 * GG)]
      PP$mu <- res[TT + 6 * GG + 1]
      PP$tausqinv <- res[TT + 6 * GG + 2]
      PP$gamma <- res[TT + 6 * GG + 3]
      PP$lambda <- res[TT + 6 * GG + 4]
      PP$pil.pos.mean <- res[TT + 6 * GG + 5]
      PP$pil.neg.mean <- res[TT + 6 * GG + 6]
      PP$pil.pos.prec <- res[TT + 6 * GG + 7]
      PP$pil.neg.prec <- res[TT + 6 * GG + 8]
      PP$kap.pos.rate <- res[TT + 6 * GG + 9]
      PP$kap.neg.rate <- res[TT + 6 * GG + 10]
      PP$poe <- matrix(res[(TT + 6 * GG + 11):(GG * (TT +
                                                       6) + TT + 10)], nrow = GG)
      PP$phat.pos <- matrix(res[(GG * (TT + 6) + TT + 11):(GG *
                                                             (2 * TT + 6) + TT + 10)], nrow = GG)
      PP$phat.neg <- matrix(res[(GG * (2 * TT + 6) + TT +
                                   11):(GG * (3 * TT + 6) + TT + 10)], nrow = GG)
      PP$accept <- res[GG * (3 * TT + 6) + TT + 11]
      gc()
      pim.pos <- c(pim.pos, mean(PP$piposg))
      pim.neg <- c(pim.neg, mean(PP$pinegg))
      if (mm > (2 * cw)) {
        pidif <- 0.5 * max(abs(mean(pim.neg[(mm - cw -
                                               1):(mm)]) - mean(pim.neg[(mm - 2 * cw - 1):(mm -
                                                                                             cw)]))/(mean(pim.neg[(mm - cw - 1):(mm)]) +
                                                                                                       mean(pim.neg[(mm - 2 * cw - 1):(mm - cw)])),
                           abs(mean(pim.pos[(mm - cw - 1):(mm)]) - mean(pim.pos[(mm -
                                                                                   2 * cw - 1):(mm - cw)]))/(mean(pim.pos[(mm -
                                                                                                                             cw - 1):(mm)]) + mean(pim.pos[(mm - 2 * cw -
                                                                                                                                                              1):(mm - cw)])))
      }
    }
 for (gg in 1:GG) {
      ppplus <- mean(PP$phat.pos[gg, ])
      ppminu <- mean(PP$phat.pos[gg, ] - PP$poe[gg, ])
      pp0 <- 1 - ppminu - ppplus
      if (ppplus < 2/TT & abs(ppminu - pp0) < 6/TT) {
        PP$phat.pos[gg, ] <- abs(PP$poe[gg, ])
        PP$poe[gg, ] <- abs(PP$poe[gg, ])
        ee <- round(PP$poe[gg, ])
        PP$mug[gg] <- mean(AA[gg, ee == 0], na.rm = TRUE)
        PP$piposg[gg] <- mean(ifelse(ee == 1, 1, 0))
        PP$pinegg[gg] <- mean(ifelse(ee == -1, 1, 0))
        PP$sigmag[gg] <- sqrt(var(AA[gg, ee == 0], na.rm = TRUE))
        PP$kappanegg[gg] <- kap.min * PP$sigmag[gg]
        PP$kappaposg[gg] <- kap.min * PP$sigmag[gg]
      }
    }
    cat("Finished\\n")
  }
  cat("Gibbs Sampler\n")
  new.PR <- unlist(PR)
  new.PP <- unlist(PP)
  avgpos <- rep(0, GG * (3 * TT + 6) + TT + 11)
  cat("First\n")
  res <- .C("poe_fit", as.matrix(AA), as.integer(NN), as.double(new.PR),
            as.double(new.PP), as.integer(GG), as.integer(TT), as.integer(M),
            as.integer(burnin), avgpos = as.double(avgpos), PACKAGE = "metaArray")$avgpos
 cat("Second\n")
  alpha <- res[1:TT]
  mug <- res[(TT + 1):(TT + GG)]
  kappaposg <- res[(TT + GG + 1):(TT + 2 * GG)]
  kappanegg <- res[(TT + 2 * GG + 1):(TT + 3 * GG)]
  sigmag <- res[(TT + 3 * GG + 1):(TT + 4 * GG)]
  piposg <- res[(TT + 4 * GG + 1):(TT + 5 * GG)]
  pinegg <- res[(TT + 5 * GG + 1):(TT + 6 * GG)]
  mu <- res[TT + 6 * GG + 1]
  tausqinv <- res[TT + 6 * GG + 2]
  gamma <- res[TT + 6 * GG + 3]
  lambda <- res[TT + 6 * GG + 4]
  pil.pos.mean <- res[TT + 6 * GG + 5]
  pil.neg.mean <- res[TT + 6 * GG + 6]
  pil.pos.prec <- res[TT + 6 * GG + 7]
  pil.neg.prec <- res[TT + 6 * GG + 8]
  kap.pos.rate <- res[TT + 6 * GG + 9]
  kap.neg.rate <- res[TT + 6 * GG + 10]
  cat("Three\n")
  poe <- matrix(res[(TT + 6 * GG + 11):(GG * (TT + 6) + TT +
                                          10)], nrow = GG)
  poe <- as.data.frame(poe)
  rownames(poe) <- rownames(AA)
  colnames(poe) <- colnames(AA)
  accept <- res[GG * (3 * TT + 6) + TT + 11]
  gc()
          cat("Four")
  foo <- function(org, fit, inter = 0.01) {
    ord <- order(org)
    org.t <- org[ord]
    fit.t <- fit[ord]
    l <- length(org)
    min.fit <- min(fit.t)
    i <- 1
    while (fit.t[i] - min.fit > inter) {
      fit.t[i] <- min.fit
      i <- i + 1
    }
    i <- length(org)
    max.fit <- max(fit.t)
    while (max.fit - fit.t[i] > inter) {
      fit.t[i] <- max.fit
      i <- i - 1
    }
    fit2 <- fit
    fit2[ord] <- fit.t
    fit2
  }
  for (i in 1:GG) {
    poe[i, ] <- foo(as.numeric(AA[i, ]), as.numeric(poe[i,
                                                        ]))
  }
  return(list(alpha = alpha, mug = mug, kappaposg = kappaposg,
              kappanegg = kappanegg, sigmag = sigmag, piposg = piposg,
              pinegg = pinegg, mu = mu, tausqinv = tausqinv, gamma = gamma,
              lambda = lambda, pil.pos.mean = pil.pos.mean, pil.pos.prec = pil.pos.prec,
              pil.neg.mean = pil.neg.mean, pil.neg.prec = pil.neg.prec,
              kap.pos.rate = kap.pos.rate, kap.neg.rate = kap.neg.rate,
              poe = poe, accept = accept))
}

data<-read.table(args[1],header=T,row.names=1,stringsAsFactors = F)
medGenes<-apply(data,1,median)
data<-data[-which(medGenes == 0),]

batchSize=2000

nbatchs=dim(data)[1]/batchSize

for (i in c(1:ceiling(nbatchs))){

subset<-data[c((((i-1)*batchSize)+1):min(i*batchSize,dim(data)[1])),]

print(paste0("Discretizing batch : ",(i-1)*batchSize+1," to ",min(i*batchSize,dim(data)[1])))
poe.Result_CenterBoth_2000<-myPOE(log(as.matrix((subset))+1),M=2000,centergene = TRUE, centersample = TRUE)
write.table(round(poe.Result_CenterBoth_2000$poe,digits=0),file=paste0("Discretised_Data_Batch_",i,".txt"),quote=FALSE,row.names = TRUE, col.names=NA,sep="\t")

}

### Supplementary Section 4 Additional gene-enhancer linkage approaches

#### Unsupervised window based linkage

For each gene $g$ in each sample $i$ we compute the number of DNase Hypersensitive sites$c_{i}^{g}$, the length of DHSs $l_{i}^{g}$, as well as the signal within the DHSs $s_{i}^{g}$ according to:

$c_{s}^{g}=\sum_{p\in P_{w,g}} I\left( p \right)*\exp\frac{\left( -dist(d_{p,g} \right)}{d_{0}})$,

$$l_{s}^{g}=\sum_{p\in P_{w,g}} \left| p \right|*\exp\left( -dist(d_{p,g} \right)/d_{0}),$$

$s_{s}^{g}=\sum_{p\in P_{w,g}} s(p)*\exp\frac{\left( -dist(d_{p,g} \right)}{d_{0}}).$

Here, $I$ is the indicator function, |$p$| denotes the length of $p,$ the function $s(p)$ denotes the signal within $p.$ $P_{w,g}$ denotes the set of DHSs $p$ within a window $w$ around the TSS of gene $g.$ As shown in Figure S2, we consider three different windows $w$: (1) a window of size 5kb centered at the TSS of gene $g$, (2) a window of size 50kb centered at the TSS of gene $g$, (3) a window considering a 2.5kb upstreaming region from the TSS of gene $g$ combined with the entire gene-body of $g.$ The input matrix of the linear regression is composed of the following terms:

|  | Count DHSs | Length DHSs | Signal DHSs |
| --- | --- | --- | --- |
| Sample 1 | $c_{1}^{g}$ | $l_{1}^{g}$ | $s_{1}^{g}$ |
| ... |  |  |  |
| Sample n | $c_{n}^{g}$ | $l_{n}^{g}$ | $s_{n}^{g}$ |

We compute these quantities using the TEPIC [12] tool using the following commands

5kb:

***bash TEPIC.sh -g hg38.noPrefix.masked.fa -b <Sample>_peaks/peaks/filtered.peaks.narrowPeak.adoptedCoverage -o <Sample>_5kb -p ../PWMs/human_jaspar_hoc_kellis.PSEM -c 12 -n 4 -w 5000 -a gencode.v26.annotation.gtf -f gencode.v26.annotation.gtf -q TRUE***

50kb:

***bash TEPIC.sh -g hg38.noPrefix.masked.fa -b <Sample>_peaks/peaks/filtered.peaks.narrowPeak.adoptedCoverage -o <Sample>_50kb -p ../PWMs/human_jaspar_hoc_kellis.PSEM -c 12 -n 4 -w 50000 -a gencode.v26.annotation.gtf -f gencode.v26.annotation.gtf -q TRUE***

gene body:

***bash TEPIC.sh -g hg38.noPrefix.masked.fa -b <Sample>_peaks/peaks/filtered.peaks.narrowPeak.adoptedCoverage -o <Sample>_gB -p ../PWMs/human_jaspar_hoc_kellis.PSEM -c 12 -n 4 -w 5000 -a gencode.v26.annotation.gtf -f gencode.v26.annotation.gtf -q TRUE -y***

The workflow is depicted in Figure S2a. Details on the parameters are provided in the TEPIC repository (www.github.com/Schulzlab/TEPIC).

#### Promoters and Enhancers extracted from GeneHancer

We downloaded the collection of all regulatory elements contained in the GeneHancer database [13], obtained at 6^th^ August, 2018 . Using an adapted version of the tool presented in this work, we compute the DNase1-seq signal within each of these regions and identify relevant segments using a correlation test. This yields a per-gene input matrix of the following form:

|  | DNase1-seq signal in GeneHancer site 1 | … | DNase1-seq signal in GeneHancer site m |
| --- | --- | --- | --- |
| Sample 1 | $g_{1}^{1}$ |  | $g_{1}^{m}$ |
| ... |  |  |  |
| Sample n | $g_{n}^{1}$ |  | $g_{n}^{m}$ |

Note that this approach does not depend on any size or regions cut-offs.

We provide a script to parse the GeneHancer tsv file into a simple tab delimited format:

*chr* ***tab*** *start* ***tab*** *end* ***tab*** *ENSG-ID* ***tab*** *gene name*

The script can be used with the command:

***python rewriteGeneHancer.py <Original GeneHancer dump> ENSGIds_GeneName.txt > <Destination.txt>***

Subsequently, the candidate regions can be computed via:

***./build/core/GENEHANCER -b <Consortium>/DNase_bw/ -a /gencode.v26.annotation.gtf -o <Consortium>_expression.txt -s data/hg38_chrSize.txt -w 25000 -p 0.05 -g <geneID> -f GeneHancer_<Consortium> -r 500000 –k genehancer_database_06.07.2018_EnsembleMatch.sorted.bed***

Here, <Consortium> is a placeholder for the data source, and <geneID> is a placeholder for the target gene. Details on the parameters are provided in the repository (www.github.com/Schulzlab/Stitch).

The workflow is depicted in Figure S2b.

#### Unification of DNase hypersensitive sites

Using bedtools [14], we combined all DHSs within a window $w$ centered at the TSS of the target gene $g$into several merged sites ${MS}^{m}.$ Within these sites, we compute the average DNase1-seq signal using an adapted version of the STITCHIT algorithm and identify sites that are correlated to the expression of gene $g$. This yields a feature matrix as shown below.

|  | DNase1-seq signal in unified site 1 | … | DNase1-seq signal in unified site m |
| --- | --- | --- | --- |
| Sample 1 | $u_{1}^{1}$ |  | $u_{1}^{m}$ |
| ... |  |  |  |
| Sample n | $u_{n}^{1}$ |  | $u_{n}^{m}$ |

We used the linux cat and sort commands together with the bedtools *merge* command

***cat <Consortium>/*/peaks/filtered.peaks.narrowPeak.adoptedCoverage

sort -s -k1,1 -k2,2n  Merged_<Consortium>.bed > Merged_<Consortium>.sorted.bed

bedtools merge -i Merged_<Consortium>.sorted.bed > Merged_<Consortium>.sorted.merged.bed***

to generate the merged DHS sites and subsequently generated the feature matrices using:

***./build/core/UNIFIED_PEAKS -b <Consortium>/DNase_wig_Normalized/ -a gencode.v26.annotation.gtf -o <Consortium>_expression.txt -s data/hg38_chrSize.txt -w 25000 -p 0.05 -g <geneID>  -f UnifiedPeaks_<Consortium> -r 500000 -k Merged_<Consortium>_Peaks.sorted.merged.bed***

The workflow is depicted in Figure S2c. Details on the parameters are provided in the repository ([www.github.com/Schulzlab/Stitchit](http://www.github.com/Schulzlab/Stitch)).

Supplementary Section 5: Supplementary Figures & Tables

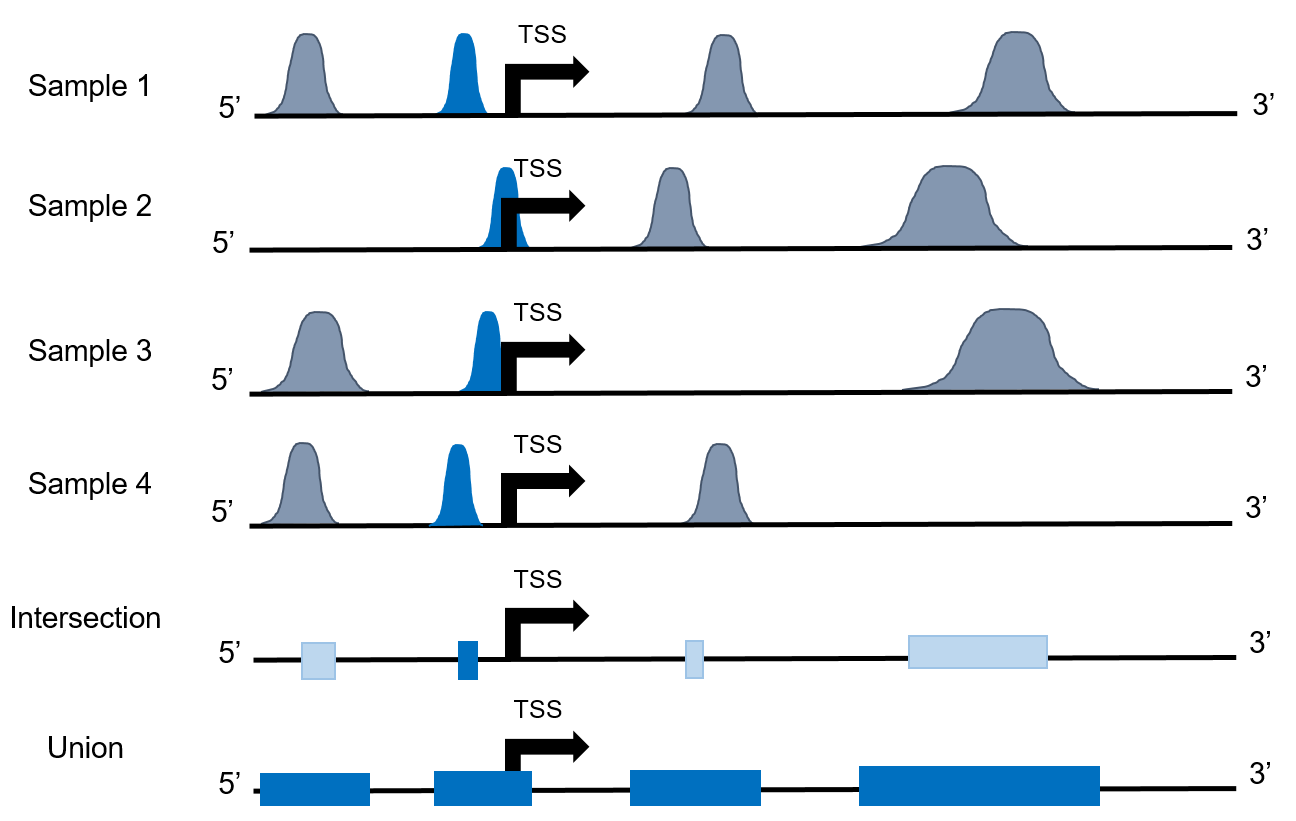

**Supplementary Figure S1:** This Figure illustrates two common ways to integrate DHS sites across multiple samples. In the intersection, only the accessible portion of the genome across all samples is available. In the union case, all overlapping DHS sites are merged. The latter case preserves a large portion of the accessible chromatin, but it loses the possible varying position of the DHSs in the data. The intersection on the other hand might be too conservative.

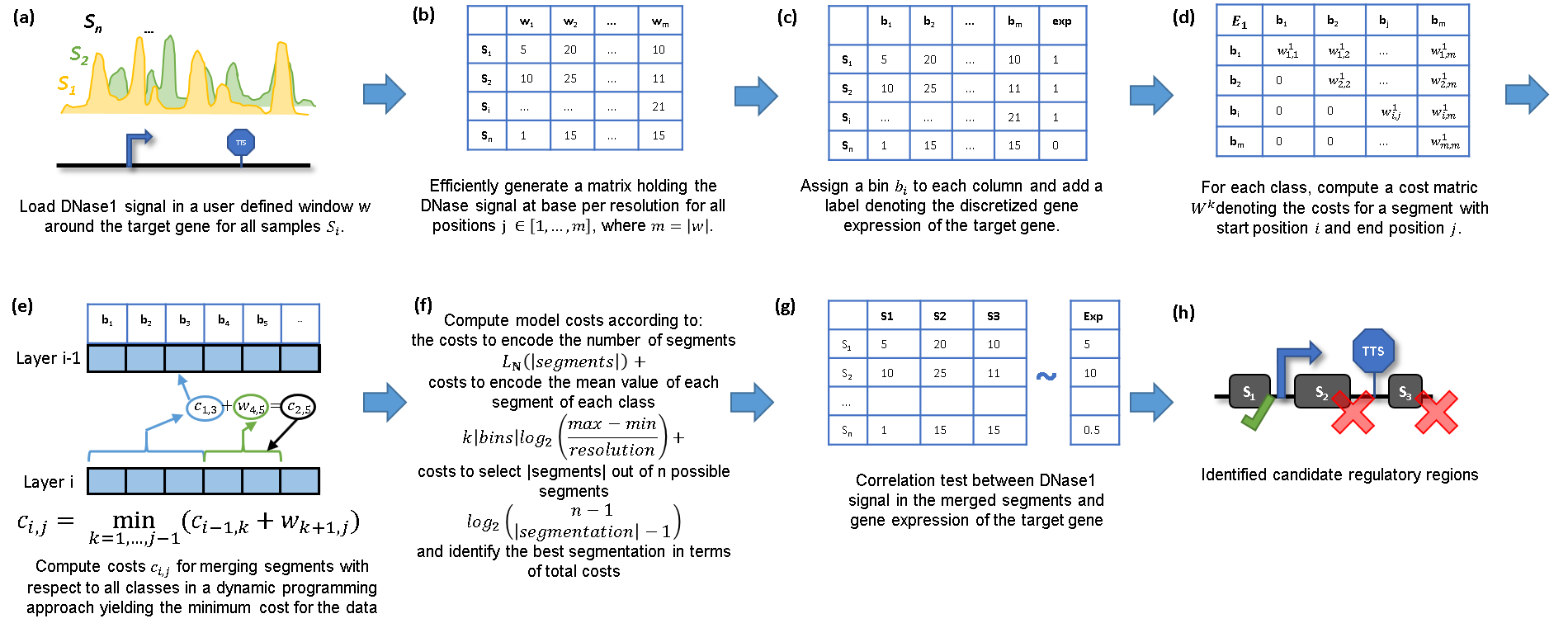

**Supplementary Figure S2:** Schematic overview on the STITCHIT algorithm. (a) Upon data generation we load DNase1-seq and gene-expression data per gene across for all available samples. (b) Using the C-library *libBigWig*, we efficiently generate a data matrix holding the DNase1-seq data at base-pair resolution. (c) We divide the data into x distinct classes referring to the discrete expression value of the target gene across the respective samples. (d) For each expression class, we compute a cost matrix that denotes the costs for generating every possible binning. (e) The optimal solution with the minimum cost is computed in a dynamic programming approach, yielding the costs of the data given a certain model (*L(D|M)*). (f) Following the MDL principle, the costs for the actual model (*L(M)*) are computed according to default formulas. (g) Via a correlation test with the continuous gene-expression data, significantly associated segments are identified, yielding one segment associated to gene expression (h).

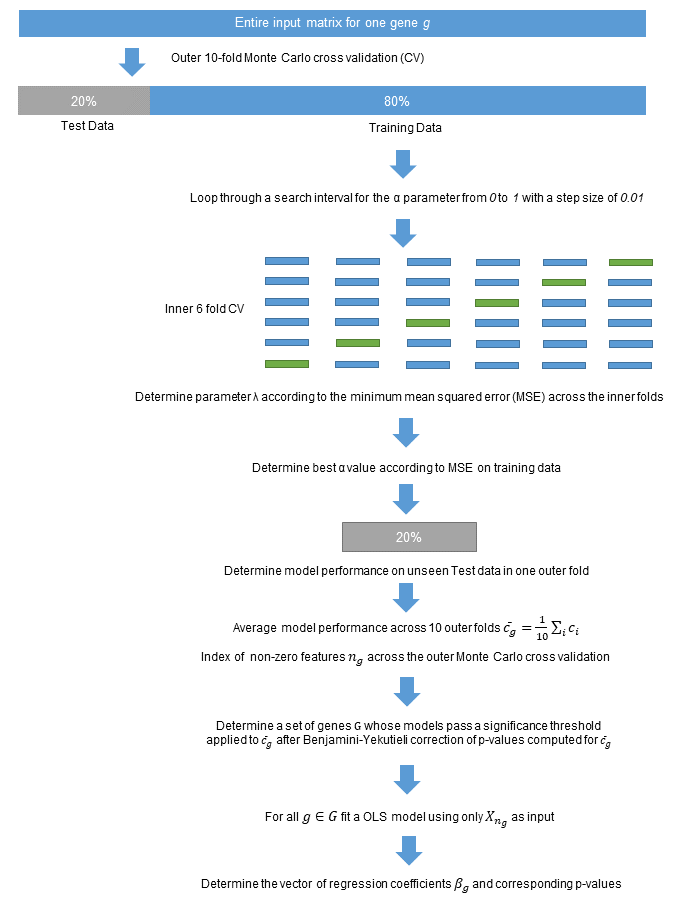

**Supplementary Figure S3**: Two level learning approach used in the manuscript. Using elastic net regularization, a set of candidate features per gene is selected. For all models achieving a sufficient model performance that is passing a significance test and a p-value correction according to Benjamini-Yekutieli, an OLS model is fitted to learn the final coefficient values and their corresponding p-values.

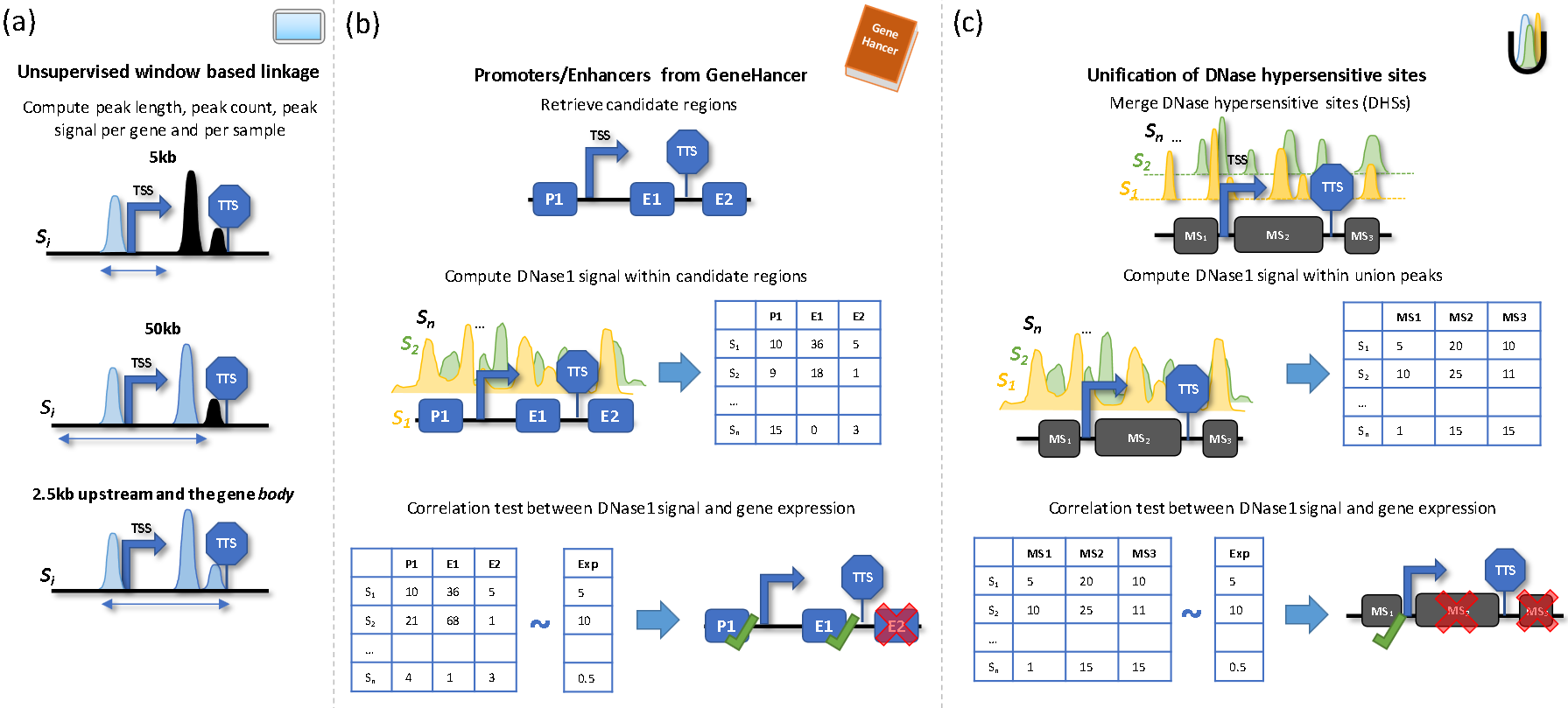

**Supplementary Figure S4**: Overview on additional gene-enhancer linkage approaches. Following a window based approach, as normally applied in per sample models (a), we assess length, count and signal of DNase hypersensitive sites (DHSs) per gene and per sample using three different windows: 5kb, 50kb, and a gene body window. Also, we consider a curated set of promoters and enhancers contained in the GeneHancer database (b). Here, we select regions that exhibit DNase1-seq signal that correlates with the expression of a target gene. Another approach is depicted in (c). Here, we identify the union of all DHS sites and select regions that exhibit DNase1-seq signal that correlates with the expression of the target gene.

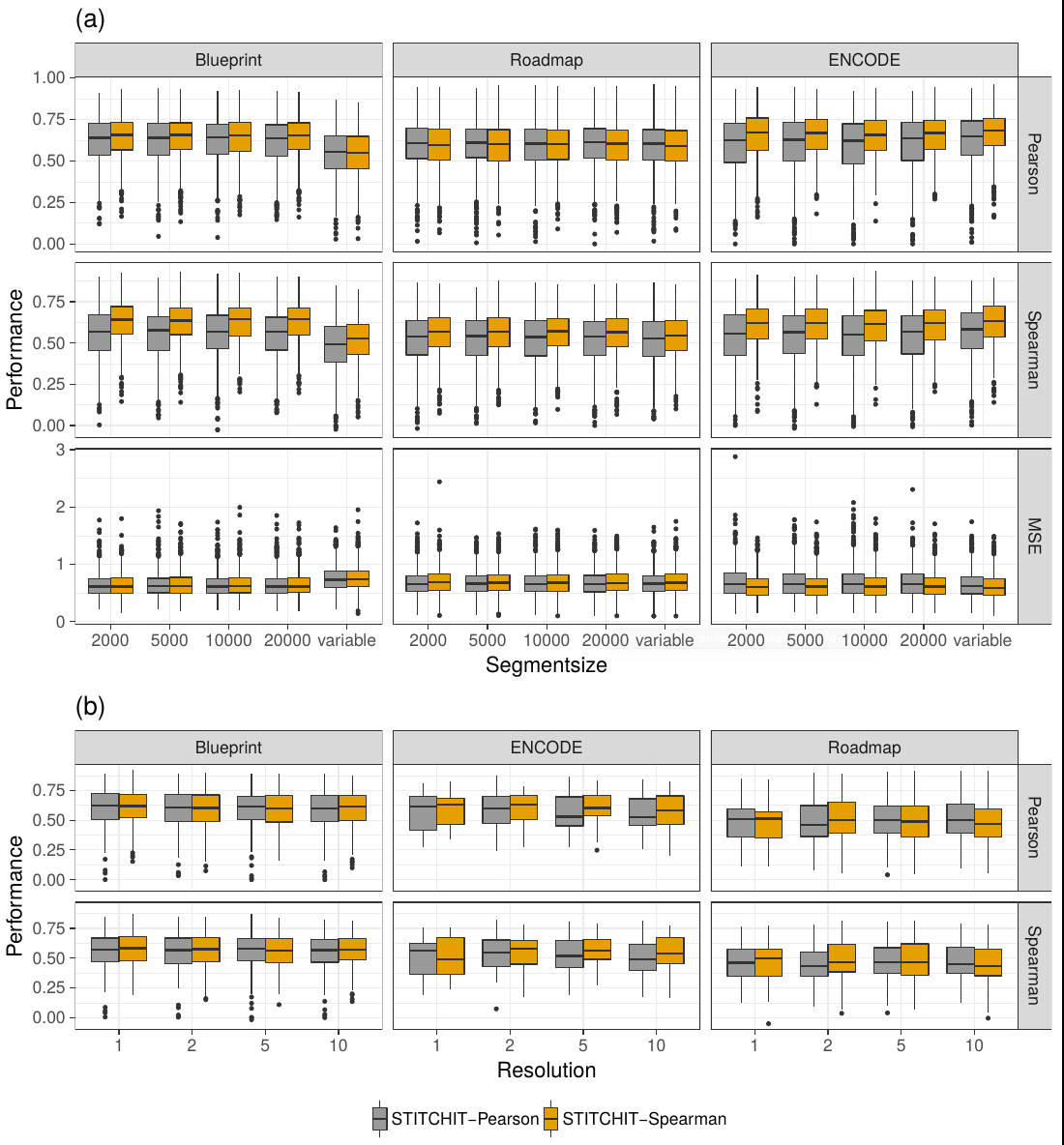

**Supplementary Figure S5:** (a) Influence of different values for the step size parameter used in STITCHIT on model performance. Importantly we note that a fixed value of the segment size is beneficial in the case of Blueprint data. (b) Influence of different values for the resolution parameter used in STITCHIT on model performance. Increasing the resolution leads to an improved runtime of STITCHIT as the initial binning is smaller.

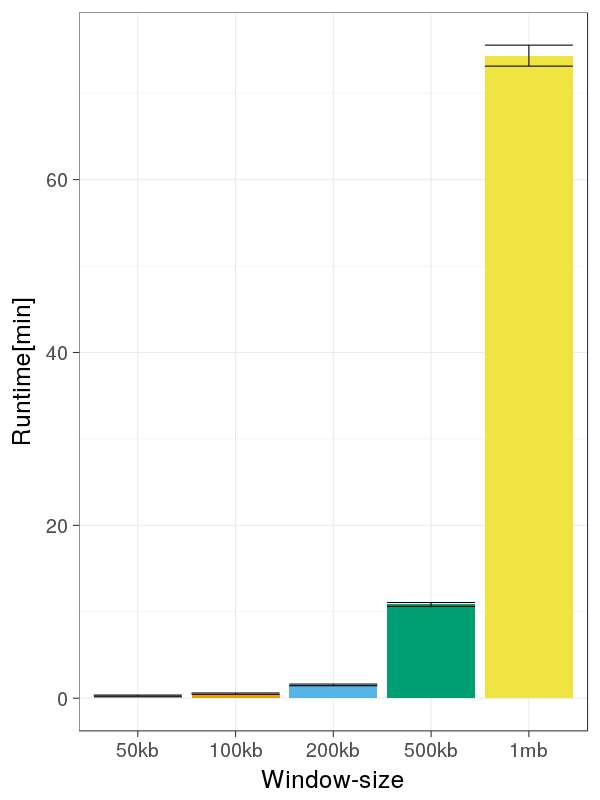

**Supplementary Figure S6**: Runtime of STITCHIT [min] depending on the used extension up- and downstream of the target gene.

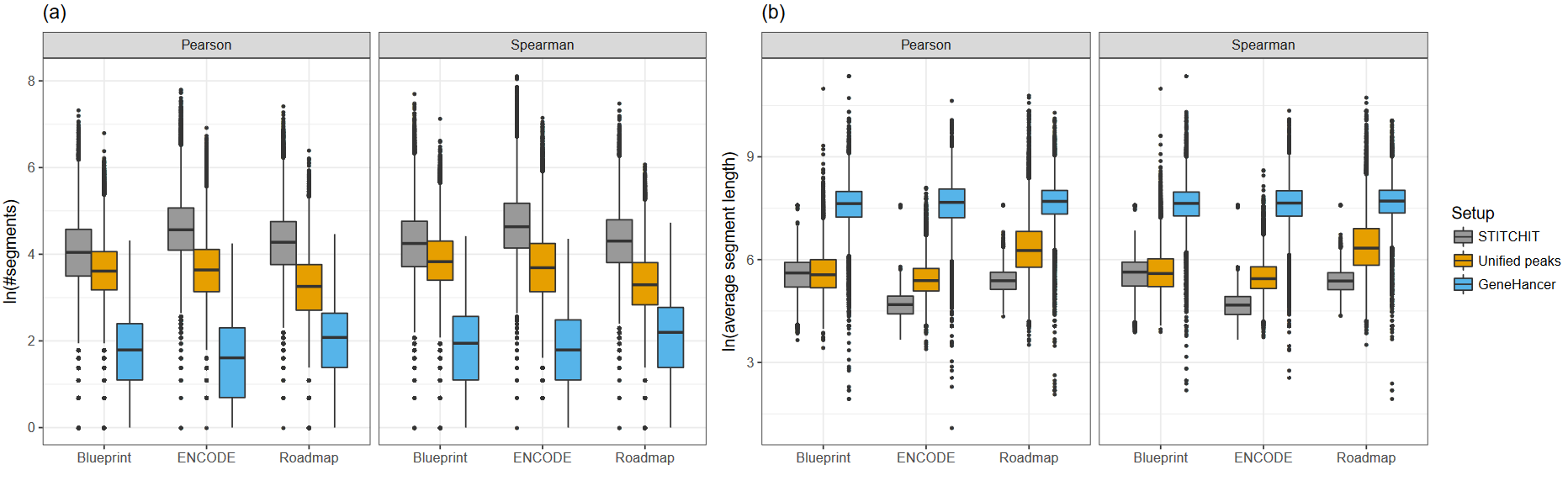

**Supplementary Figure S7:** (a) Number of segments suggested by STITCHIT, Unified peaks or GeneHancer using either Pearson or Spearman correlation as an initial filtering. (b) Length of segments suggested by STITCHIT, Unified peaks or GeneHancer using either Pearson or Spearman correlation as an initial filtering.

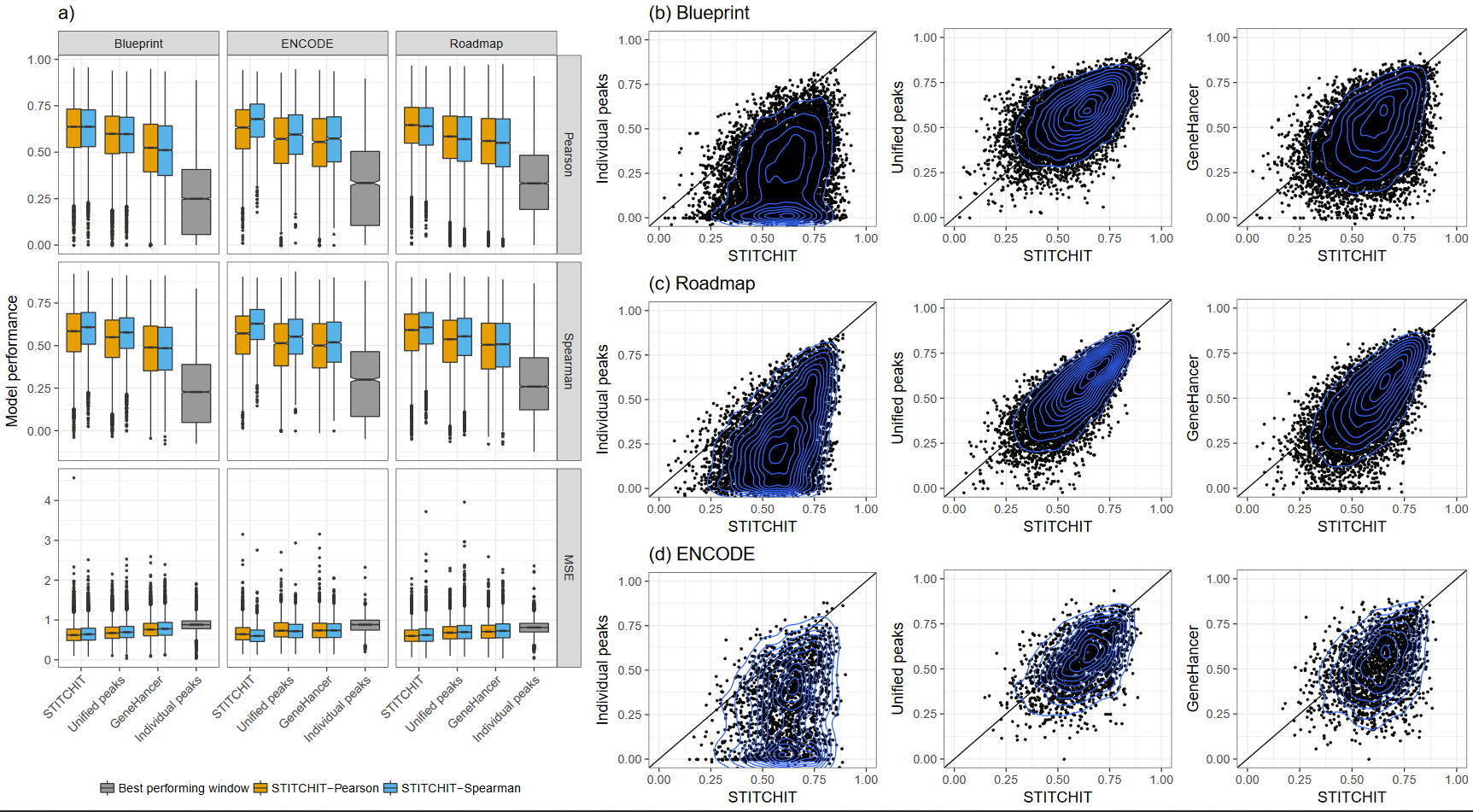

**Supplementary Figure S8:** (a) Model performance assessed in terms of Pearson and Spearman correlation as well as using the Mean Squared error (MSE) for all consortia. The models are compared on the same gene sets within one consortia. (b-d) Scatter plots comparing the spearman correlation of genes for two distinct models on Blueprint, Roadmap and ENCODE data respectively.

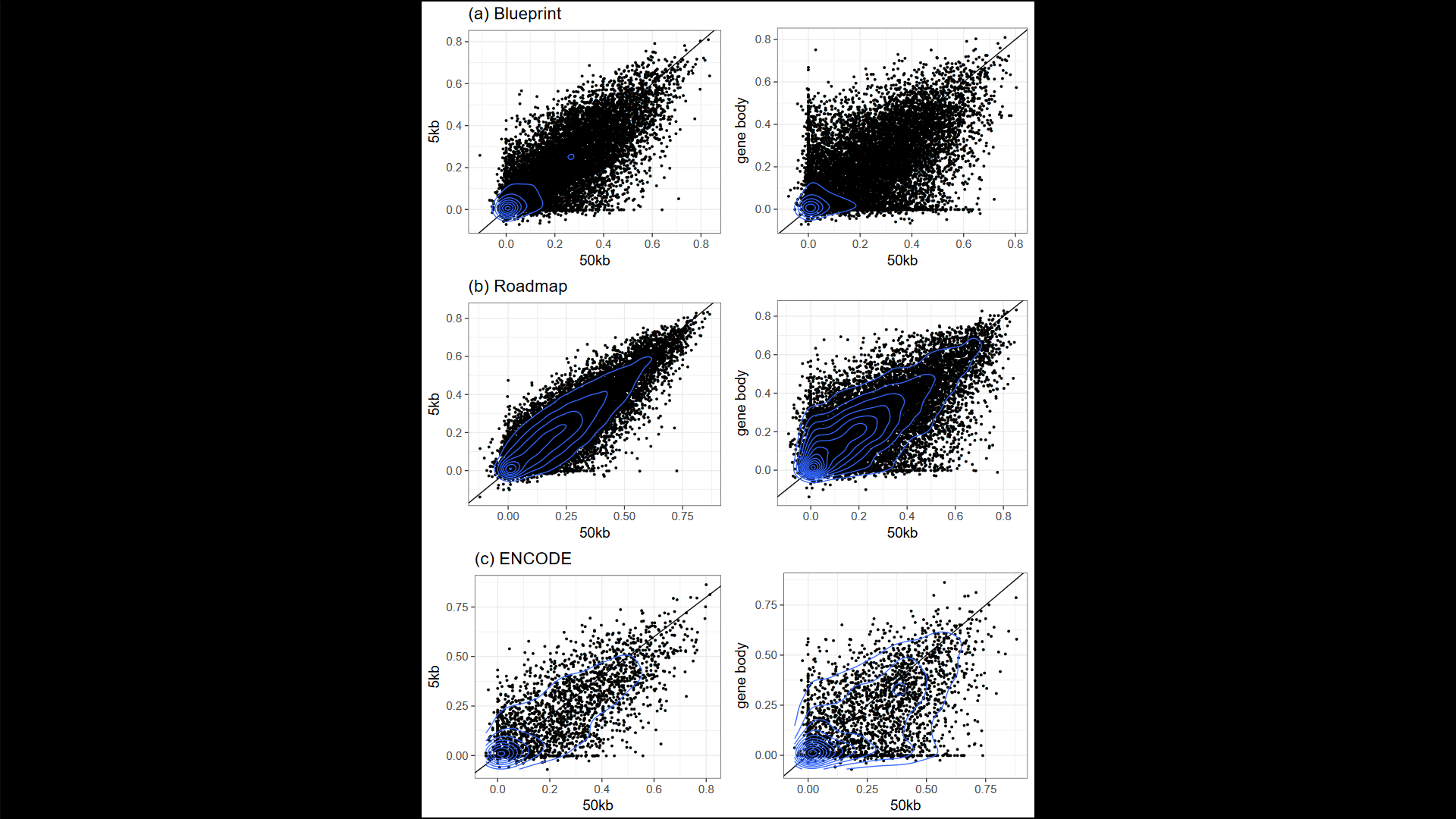

**Supplementary Figure S9:** Scatterplots comparing different unsupervised annotation approaches (5kb, 50kb, genebody) on Blueprint (a), Roadmap (b) and ENCODE (c) data.

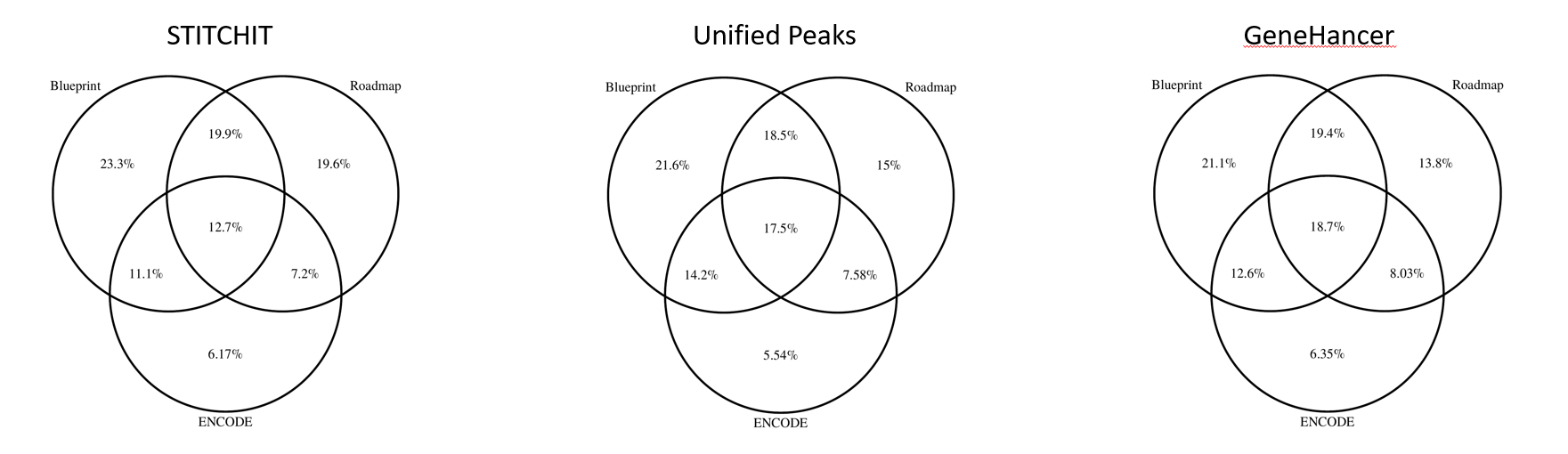

**Supplementary Figure S10**: Venn Diagramms indicating the overlap in terms of covered genes for STITCHIT, Unified Peaks, and GeneHancer data respectively.

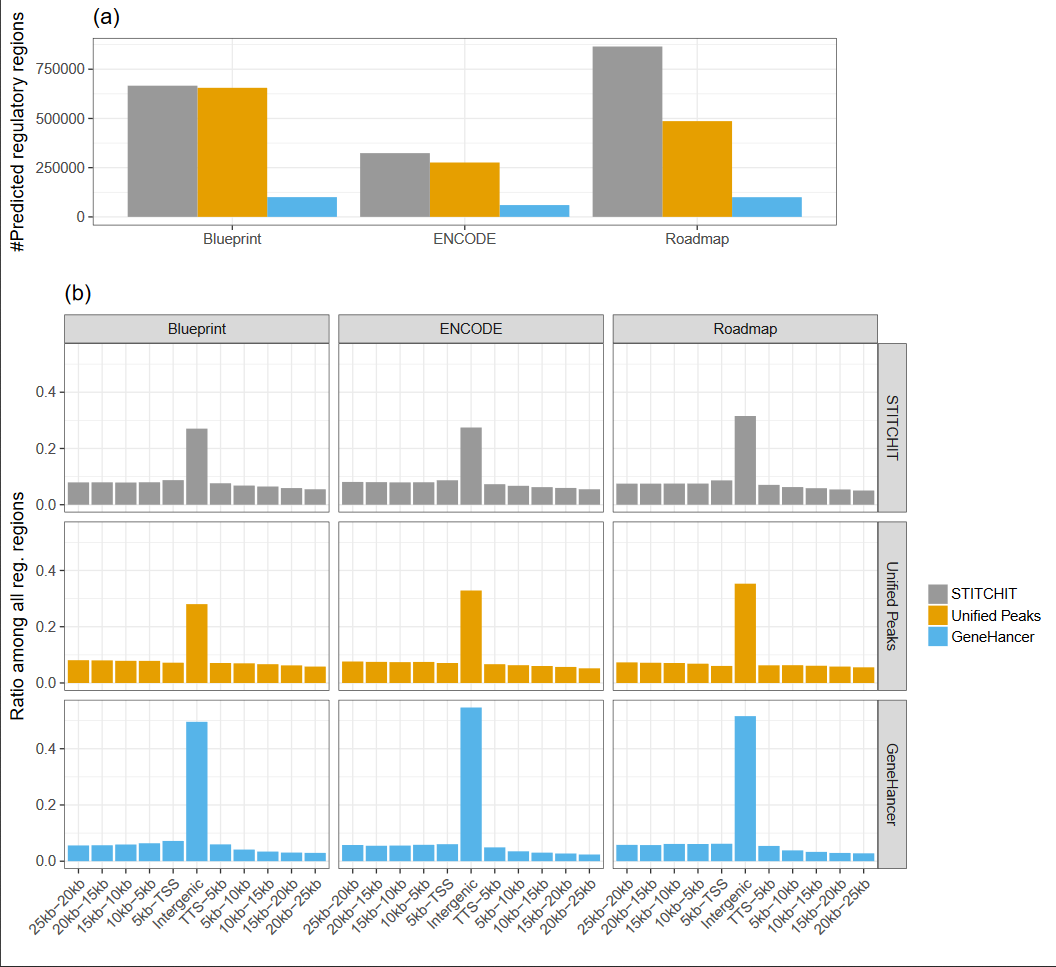

**Supplementary Figure S11**: Distribution of PEIs around genes for all datasets and all methods. We observe a peak in the intergenic part of the gene and a decline downstream of the gene.

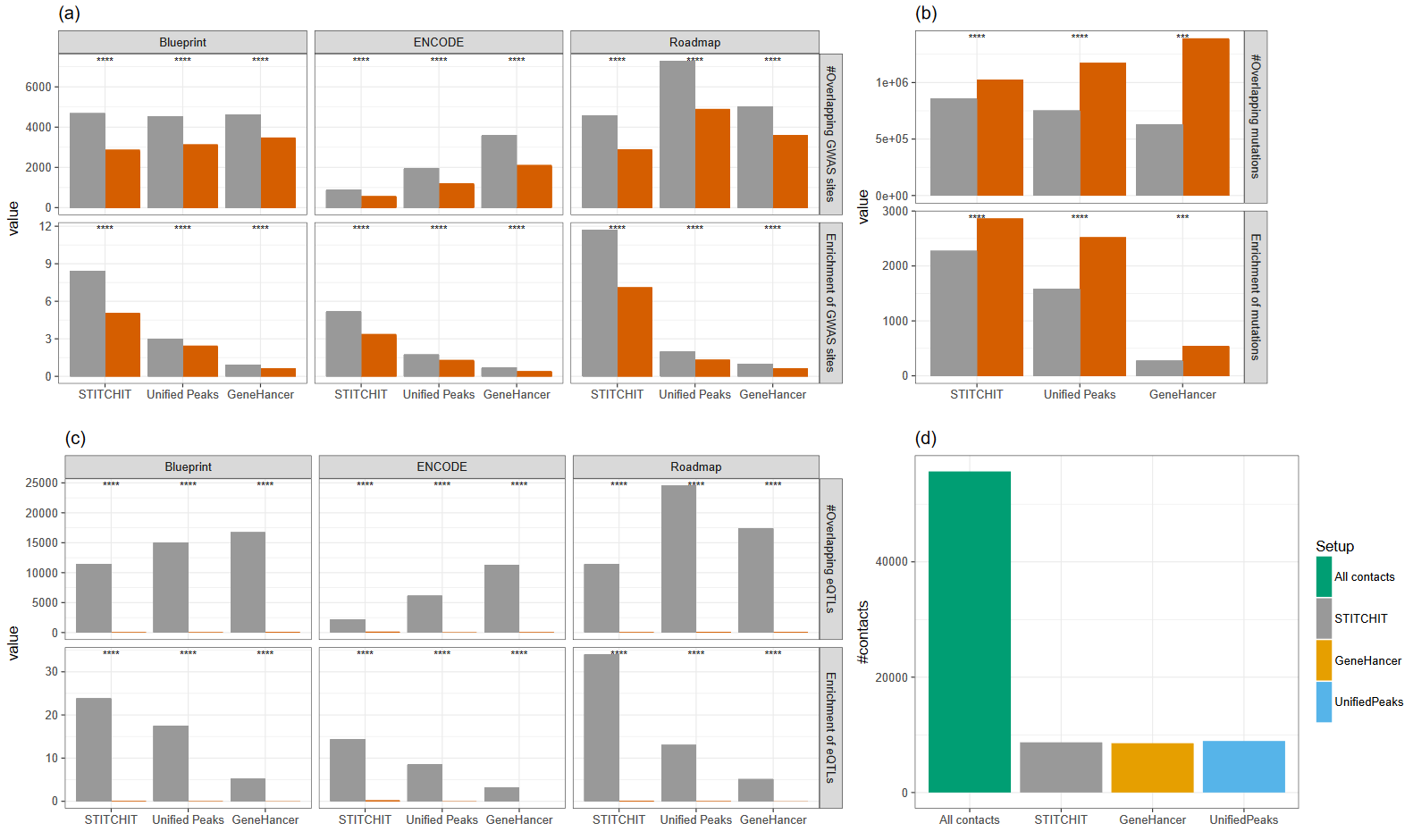

**Supplementary Figure S12**: (a) Random comparison to overlapping GWAS sites indicating the total number of overlapping sites as well as the enrichment score. (b) Total number of overlapping mutations from the COSMIC database and the enrichment score compared between randomized and original data. (c) Random analysis of the eQTL data from expSNP showing the total number of correctly overlapping eQTLs as well as the enrichment score. (d) Total number and retrieved number of contacts from a Capture Hi-C experiment. Significance in a)-c) is indicated by a two-sided t-test (****: p≤ 0.0001).

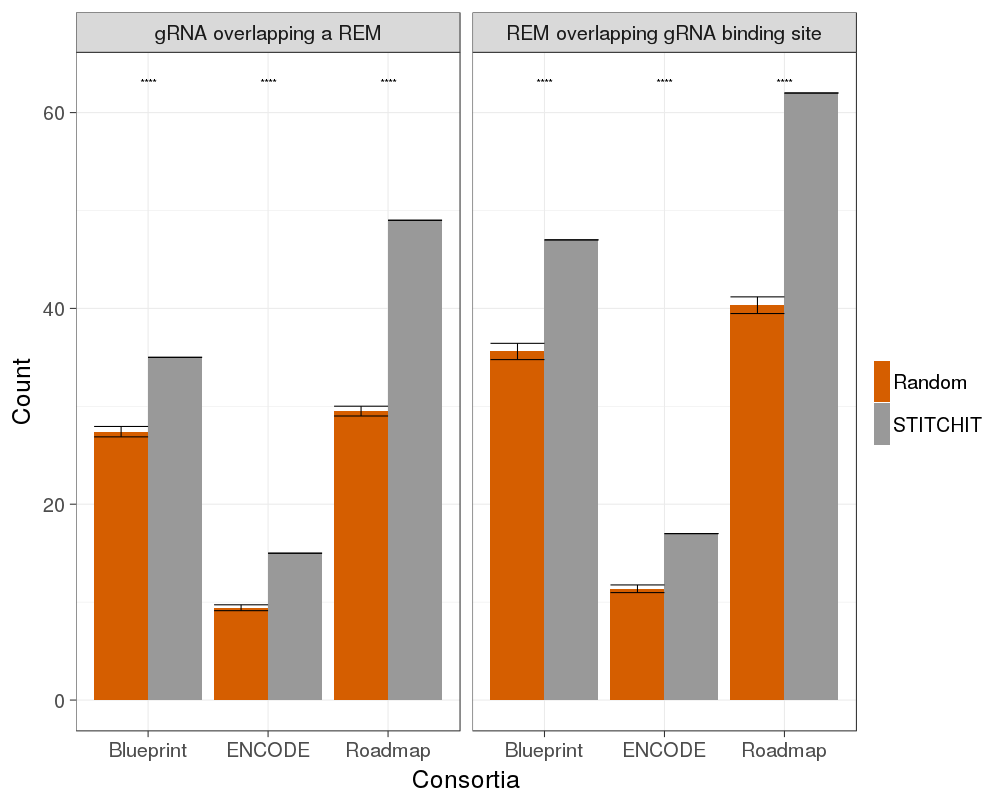

**Supplementary Figure S13**: The Figure shows the mean number of gRNAs overlapping at least one REM, and the mean number of REMs overlapping at least one gRNA binding site for STITCHIT predicted REMs as well as for Randomly ten rounds of randomly selected regions in the genome with the same length distribution as the STITCHIT regions. Significance in a)-c) is indicated by a two-sided t-test (****: p≤ 0.0001). Error bars indicate standard error. The standard deviation of gRNA overlapping a REM is 10,1 for the random samples across the consortia and 14.58 for REM overlapping gRNA binding sties.

(a)

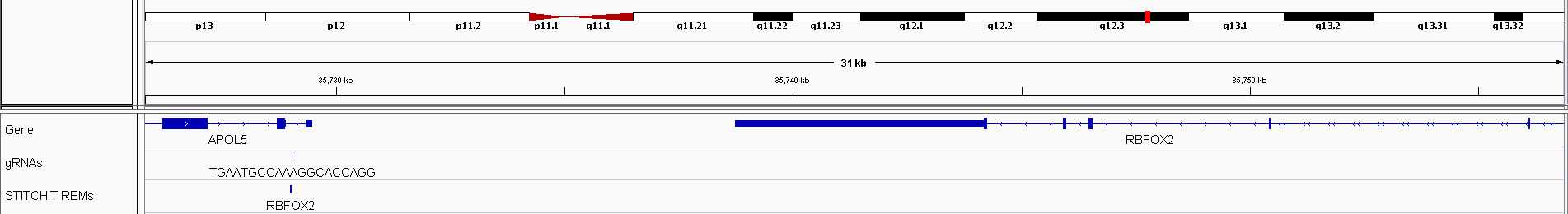

(b)

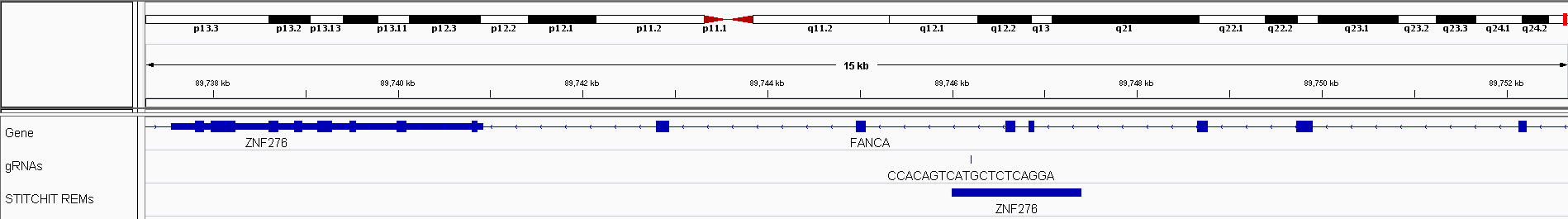

(c)

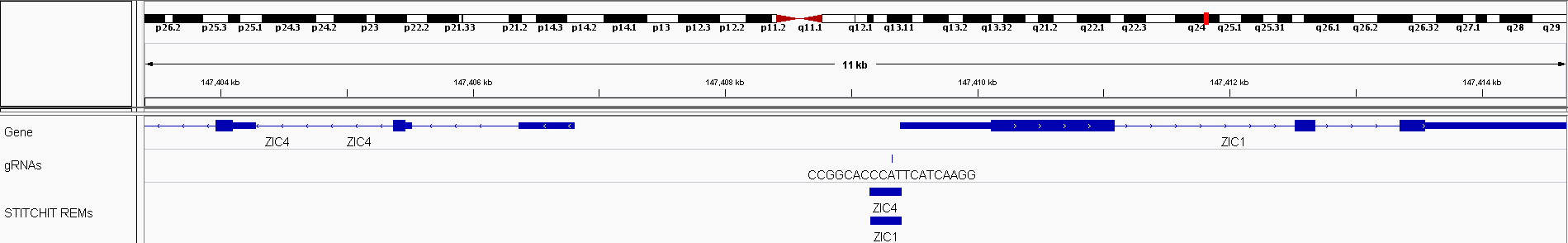

(d)

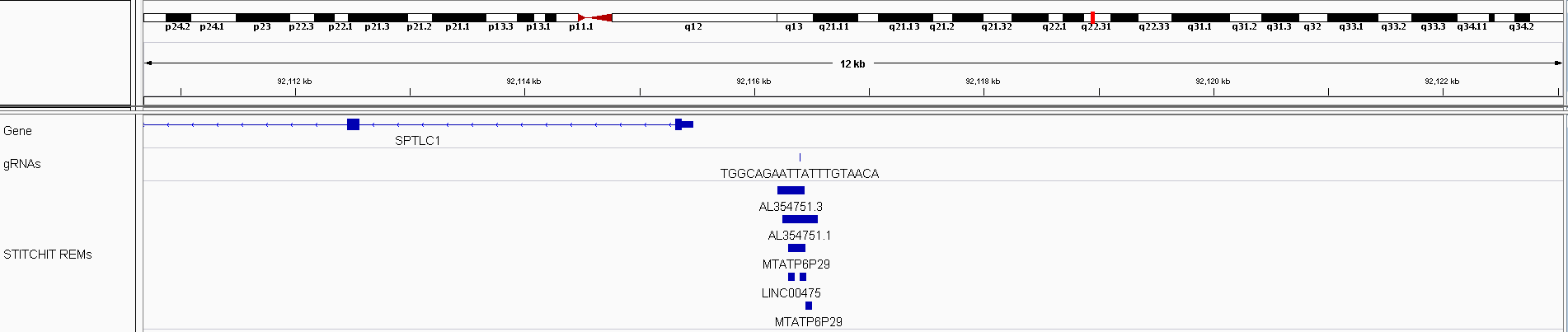
**Supplementary Figure S14**: In (a) and (b), REMs detected in introns of APOL5 and FANCA, serving as enhancers for RBFOX2 and ZNF276, respectively, are shown. In (c) a gRNA site overlapping with REMs for ZIC4 and ZIC1 are depicted while in (d) an extreme case for one gRNA site being linked to several non-coding genes is depicted.

| **gRNA ID** | **DHS** | **Genomic coordinates** |
| --- | --- | --- |
| Chr17.1549.5 | Chr17.1549 | 37856119-37856138 |
| Chr17.1553.4 | Chr17.1553 | 37862556-37862575 |
| Chr17.1561.8 | Chr17.1561 | 37896930-37896949 |
| Chr17.1559.22 | Chr17.1559 | 37894617-37894636 |
| Chr17.1560.27 | Chr17.1560 | 37895566-37895585 |
| Chr17.1562.28 | Chr17.1562 | 37906499-37906518 |
| Chr17.1550.11 | Chr17.1550 | 37858091-37858110 |
| Chr17.1551.30 | Chr17.1551 | 37859298-37859317 |
| Chr17.1552.9 | Chr17.1552 | 37860006-37860025 |
| Chr17.1548.4 | Chr17.1548 | 37850995-37851014 |
| Chr17.1563.27 | Chr17.1563 | 37910314-37910333 |

**Supplementary Table S2**: gRNA IDs as well as corresponding DHS IDs used for the ERBB2 experiment

| **chr** | **start** | **end** |
| --- | --- | --- |
| chr17 | 39663644 | 39663803 |
| chr17 | 39667004 | 39667153 |
| chr17 | 39672254 | 39672403 |
| chr17 | 39675454 | 39676103 |
| chr17 | 39676294 | 39676453 |
| chr17 | 39687254 | 39687453 |
| chr17 | 39687594 | 39687743 |
| chr17 | 39699894 | 39700003 |
| chr17 | 39702694 | 39702913 |
| chr17 | 39703654 | 39703703 |
| chr17 | 39705854 | 39706053 |
| chr17 | 39723894 | 39724003 |
| chr17 | 39729604 | 39730043 |
| chr17 | 39737204 | 39737753 |
| chr17 | 39739304 | 39739453 |
| chr17 | 39740804 | 39740913 |
| chr17 | 39750104 | 39750203 |
| chr17 | 39754704 | 39754913 |

**Supplementary Table S3:** STITCHIT reems for ERBB2 overlapping DHSs of SKBR3 cells

| **chr** | **start** | **End** |
| --- | --- | --- |
| chr17 | 39674899 | 39675489 |
| chr17 | 39677742 | 39678129 |
| chr17 | 39686754 | 39687005 |
| chr17 | 39687110 | 39689386 |
| chr17 | 39699367 | 39702351 |
| chr17 | 39702548 | 39703300 |
| chr17 | 39704371 | 39706387 |
| chr17 | 39737892 | 39738050 |
| chr17 | 39740310 | 39740914 |
| chr17 | 39753060 | 39757532 |

**Supplementary Table S4:** Reems called using the Unified Peak approach for ERBB2 overlapping DHSs of SKBR3 cells

| **chr** | **start** | **End** |
| --- | --- | --- |
| chr17 | 39630988 | 39635001 |
| chr17 | 39673953 | 39681361 |
| chr17 | 39686731 | 39689728 |
| chr17 | 39694339 | 39697219 |
| chr17 | 39698981 | 39707766 |
| chr17 | 39723315 | 39723967 |
| chr17 | 39728818 | 39731177 |
| chr17 | 39784411 | 39789373 |
| chr17 | 39790476 | 39796594 |
| chr17 | 39812390 | 39815058 |

**Supplementary Table S5:** Regulatory elements from the GeneHancer database for ERBB2 overlapping DHSs of SKBR3 cells

| **chr** | **R start** | **R end** | **GeneID** | **coef** | **p-value** | **ChIA-Pet** | **Gene**  **Hancer** | **gRNA** | **gRNA start** | **gRNA end** |
| --- | --- | --- | --- | --- | --- | --- | --- | --- | --- | --- |
| chr3 | 147409152 | 147409401 | ENSG00000152977 | 0.378326275 | 1.74E-005 |  | X | CCGGCACCCATTCATCAAGG | 147409223 | 147409423 |
| chr11 | 43949847 | 43950306 | ENSG00000244953 | -0.274825376 | 2.95E-005 |  |  | TTATAGAGTTTCCTTTGCCA | 43950223 | 43950423 |
| chr17 | 28404050 | 28404239 | ENSG00000265254 | 0.481973601 | 0.001400099 |  |  | TTTAAGAACATATTAATCGG | 28404183 | 28404383 |
| chr3 | 134378494 | 134378603 | ENSG00000114019 | 0.328670581 | 0.002697551 |  | X | CTGCCTGTTGCCAAAAGCAG | 134378350 | 134378550 |
| chr3 | 134378494 | 134378603 | ENSG00000114019 | 0.328670581 | 0.002697551 |  | X | CTGCCTGTTGCCAAGAGCAG | 134378350 | 134378550 |
| chr17 | 42093046 | 42093155 | ENSG00000187595 | 0.187034942 | 0.002857962 |  |  | TGGCAGAATTATTTGTAACA | 42093081 | 42093281 |
| chr17 | 48740854 | 48740893 | ENSG00000244514 | -0.598451082 | 0.004260338 |  |  | CCATTTTTTCCCCCGATAGA | 48740701 | 48740901 |
| chr14 | 23008203 | 23008892 | ENSG00000279656 | -0.375730282 | 0.008893199 | X |  | AATAGGGCTCCTATAACACA | 23008590 | 23008790 |
| chr3 | 114219101 | 114219600 | ENSG00000174255 | 0.322885945 | 0.009562447 |  |  | TCAAGTAAAGGTGTGCACAG | 114219495 | 114219695 |
| chr6 | 149919000 | 149920999 | ENSG00000131015 | -0.169969304 | 0.01740522 |  |  | GCACTTGTAGCGGCGTTAAA | 149920916 | 149921116 |
| chr6 | 149230595 | 149230744 | ENSG00000283608 | -0.580879058 | 0.018230986 |  |  | GGGAGCCTCCTTGCGCTACA | 149230624 | 149230824 |
| chr6 | 35460847 | 35461346 | ENSG00000007866 | -0.166890804 | 0.021030043 |  |  | GTGCACATATGGGCTAGGCG | 35461211 | 35461411 |
| chr15 | 93351647 | 93351956 | ENSG00000257060 | -0.145040624 | 0.026855206 |  |  | GCTTACTGCACCAATCACCA | 93351822 | 93352022 |
| chr20 | 45993548 | 45993757 | ENSG00000100985 | -0.167059874 | 0.03118279 | X | X | CCTACTCCTCAATTTCCCCA | 45993723 | 45993923 |
| chrX | 19894200 | 19894299 | ENSG00000173681 | -0.122213972 | 0.033450487 |  |  | CCTACCCCACCAAAATCCCA | 19894067 | 19894267 |
| chr20 | 50212698 | 50212807 | ENSG00000277449 | 0.271475446 | 0.037002049 |  |  | CCTCTCGCCAGCCATCTCAA | 50212667 | 50212867 |
| chr10 | 73358503 | 73358702 | ENSG00000233144 | 0.23456791 | 0.037403606 |  |  | CGGAGGCCTCCGACTAGAAG | 73358365 | 73358565 |
| chr9 | 99202096 | 99202205 | ENSG00000119523 | -0.209406396 | 0.038562862 | X |  | CATATCCTTAAGAATTAGCA | 99201985 | 99202185 |
| chr3 | 147409146 | 147409405 | ENSG00000174963 | 0.377582252 | 0.042256958 |  | X | CCGGCACCCATTCATCAAGG | 147409223 | 147409423 |
| chr9 | 92116298 | 92116447 | ENSG00000232179 | -0.296081695 | 0.057251487 | X |  | TGGCAGAATTATTTGTAACA | 92116300 | 92116500 |
| chr9 | 130742706 | 130743055 | ENSG00000224797 | -0.159784083 | 0.069104178 |  |  | AAATCAAGATCTTAACAAAA | 130742909 | 130743109 |
| chr18 | 31917003 | 31917152 | ENSG00000153339 | 0.166626727 | 0.081261789 |  |  | CCATCAATATATATGTCACA | 31916875 | 31917075 |
| chr1 | 212621054 | 212622333 | ENSG00000162772 | 0.853205857 | 0.083075455 | X | X | TGAATTTCCTCGAATTTGCA | 212621220 | 212621420 |
| chr15 | 48651001 | 48651350 | ENSG00000273925 | -0.164491956 | 0.089737634 |  |  | ATTCATAAAATCAAAAAGAG | 48650897 | 48651097 |
| chr10 | 73358499 | 73358698 | ENSG00000156042 | 0.53519506 | 0.092722936 |  | X | CGGAGGCCTCCGACTAGAAG | 73358365 | 73358565 |
| chr9 | 92116448 | 92116507 | ENSG00000232179 | 0.189450866 | 0.106388303 | X |  | TGGCAGAATTATTTGTAACA | 92116300 | 92116500 |
| chr5 | 140090947 | 140091606 | ENSG00000185129 | -0.157041833 | 0.116161307 |  |  | GCAGTAAATGTGGTGAGAGG | 140091078 | 140091278 |
| chr16 | 30020947 | 30021106 | ENSG00000149926 | 0.475787434 | 0.120981011 |  |  | AATGCATCAGCGCAGCACAG | 30020911 | 30021111 |
| chr10 | 113850401 | 113850600 | ENSG00000196865 | -0.219384309 | 0.1481812 |  |  | CTGCCTGCTGGCTCAAAGCA | 113850460 | 113850660 |
| chr8 | 120445460 | 120445499 | ENSG00000172167 | -2.588460612 | 0.155609711 |  | X | CCTGCTCCTGGTGATATGGG | 120445381 | 120445581 |
| chr3 | 186615799 | 186616898 | ENSG00000090512 | 0.152998161 | 0.155877699 |  |  | CCTTCCCCACTTCCTCGACA | 186616458 | 186616658 |
| chr8 | 120445450 | 120445459 | ENSG00000172167 | 3.125332421 | 0.15810417 |  |  | CCTGCTCCTGGTGATATGGG | 120445381 | 120445581 |
| chr10 | 89411649 | 89412148 | ENSG00000152779 | -0.426390776 | 0.159552393 |  |  | GCGACTCTCCAAAAGCGCAA | 89411571 | 89411771 |
| chr17 | 81635351 | 81635400 | ENSG00000182612 | 0.24330982 | 0.162623163 |  | X | GAACCTAGCATTTATTTGAA | 81635398 | 81635598 |
| chr3 | 186616353 | 186616852 | ENSG00000145192 | 0.180161833 | 0.164895397 |  | X | CCTTCCCCACTTCCTCGACA | 186616458 | 186616658 |
| chr9 | 92116398 | 92116457 | ENSG00000225511 | -0.159616831 | 0.169251307 |  |  | TGGCAGAATTATTTGTAACA | 92116300 | 92116500 |
| chr20 | 49830125 | 49831194 | ENSG00000237788 | 0.188372895 | 0.176663252 |  |  | CTATCTCCCTCTTCATCACA | 49831024 | 49831224 |
| chr17 | 65181452 | 65181551 | ENSG00000108370 | 0.165611915 | 0.176843689 | X | X | CAAGGTATTGAAGCTAAAAG | 65181462 | 65181662 |
| chr8 | 93022850 | 93022899 | ENSG00000205133 | 0.195155507 | 0.184574517 |  |  | ACTATTCCTTACAAGTAAAG | 93022700 | 93022900 |
| chr8 | 93022850 | 93022899 | ENSG00000205133 | 0.195155507 | 0.184574517 |  |  | ACTTTTCCTTACCAGTAAAG | 93022700 | 93022900 |
| chr8 | 93022850 | 93022899 | ENSG00000205133 | 0.195155507 | 0.184574517 |  |  | TCTATTCCTTACCAGTAAAG | 93022700 | 93022900 |
| chr12 | 102127943 | 102128052 | ENSG00000075188 | -0.121815411 | 0.187684234 |  |  | TCTAAGTCCACACAGTCAAA | 102127745 | 102127945 |
| chr10 | 73358351 | 73358550 | ENSG00000138279 | 0.214164144 | 0.190596661 |  |  | CGGAGGCCTCCGACTAGAAG | 73358365 | 73358565 |
| chr16 | 30021099 | 30021208 | ENSG00000149927 | 0.181801372 | 0.19075819 | X | X | AATGCATCAGCGCAGCACAG | 30020911 | 30021111 |
| chr17 | 28404306 | 28404455 | ENSG00000076351 | 0.25608548 | 0.1930928 |  | X | TTTAAGAACATATTAATCGG | 28404183 | 28404383 |
| chr16 | 30021001 | 30021100 | ENSG00000149929 | -0.115797749 | 0.198426634 |  |  | AATGCATCAGCGCAGCACAG | 30020911 | 30021111 |
| chr7 | 142929543 | 142929602 | ENSG00000165131 | -0.083241641 | 0.211633889 |  |  | CTTCCGCCCTCGGCTTGGCA | 142929574 | 142929774 |
| chr8 | 120445350 | 120445449 | ENSG00000172167 | -1.026563442 | 0.225818351 |  |  | CCTGCTCCTGGTGATATGGG | 120445381 | 120445581 |
| chr17 | 28404357 | 28404646 | ENSG00000258924 | 0.196362209 | 0.260259001 |  |  | TTTAAGAACATATTAATCGG | 28404183 | 28404383 |
| chr2 | 64089353 | 64089652 | ENSG00000228079 | 0.255300707 | 0.263143276 |  |  | TAAGAATCCCTCCTGATGAA | 64089573 | 64089773 |
| chr4 | 139651558 | 139651647 | ENSG00000085871 | 0.120460039 | 0.264610385 |  | X | CTGCCTGCCCGGTCTCCCAA | 139651536 | 139651736 |
| chr18 | 24627802 | 24627951 | ENSG00000265485 | -0.072310988 | 0.27634866 |  |  | TATATTAATACCATATAGAG | 24627862 | 24628062 |
| chr7 | 140331753 | 140332692 | ENSG00000157800 | 0.09489202 | 0.282261166 |  |  | CTGGCCCCCAAATCAATACA | 140331748 | 140331948 |
| chr7 | 128938897 | 128939156 | ENSG00000275106 | -0.310745846 | 0.296485428 | X |  | GGACCCATAACTACTCGGGG | 128939109 | 128939309 |
| chr17 | 42093194 | 42093353 | ENSG00000108771 | 0.150817299 | 0.297490601 |  |  | TGGCAGAATTATTTGTAACA | 42093081 | 42093281 |
| chrX | 40943902 | 40943981 | ENSG00000216866 | -0.239128373 | 0.299841717 |  |  | GCTGGGAACCTGGCTATAAA | 40943875 | 40944075 |
| chr20 | 50212804 | 50212903 | ENSG00000172216 | 0.122891307 | 0.301708705 | X |  | CCTCTCGCCAGCCATCTCAA | 50212667 | 50212867 |
| chr17 | 80399898 | 80400827 | ENSG00000263069 | 0.092610397 | 0.308724059 |  |  | CCTGCCGTCATTATACACCA | 80400448 | 80400648 |
| chr2 | 64089653 | 64089902 | ENSG00000228079 | 0.162027634 | 0.315780086 |  |  | TAAGAATCCCTCCTGATGAA | 64089573 | 64089773 |
| chr22 | 35728996 | 35729045 | ENSG00000100320 | -0.072352143 | 0.318897319 |  |  | TGAATGCCAAAGGCACCAGG | 35728951 | 35729151 |
| chr19 | 11111302 | 11111451 | ENSG00000130164 | 0.119137874 | 0.319741622 | X |  | CCTGACGTCATTATTCACCA | 11111224 | 11111424 |
| chr1 | 202938350 | 202939879 | ENSG00000199471 | -0.095500771 | 0.330872399 |  |  | CCGCTAGTATAAGAAAAGGG | 202938430 | 202938630 |
| chr1 | 40036653 | 40036742 | ENSG00000131236 | 0.133279466 | 0.33534476 |  | X | CTATCTTGCTTCCCTTCCAA | 40036552 | 40036752 |
| chr17 | 77894027 | 77894456 | ENSG00000204283 | 0.063671599 | 0.3376392 |  | X | CTGAGGAGTTGCTCGAGACA | 77894331 | 77894531 |
| chr18 | 3632150 | 3633249 | ENSG00000262001 | 0.163875499 | 0.358751563 |  | X | TAAAGCCCATAACTTCCCCA | 3632948 | 3633148 |
| chrX | 1352046 | 1352755 | ENSG00000185291 | 0.451293933 | 0.363103106 |  |  | CGACAAACTTATCTGTGCAG | 1352368 | 1352568 |
| chr9 | 92116248 | 92116557 | ENSG00000234537 | 0.119489106 | 0.371706339 |  |  | TGGCAGAATTATTTGTAACA | 92116300 | 92116500 |
| chr16 | 4045102 | 4045201 | ENSG00000263159 | -0.186393535 | 0.372966401 |  |  | GCTTTAGACAAAGTTCTGAA | 4044989 | 4045189 |
| chr13 | 87671045 | 87671194 | ENSG00000165300 | 0.105043238 | 0.37437256 |  | X | CGTGACAGCAGCATACTGAA | 87671002 | 87671202 |
| chr18 | 26135692 | 26136551 | ENSG00000154611 | -0.091288986 | 0.381219399 |  |  | CTATAAGAAAAGCATCAACA | 26136043 | 26136243 |
| chr17 | 48740203 | 48741002 | ENSG00000159184 | 0.132337474 | 0.382191782 |  |  | CCATTTTTTCCCCCGATAGA | 48740701 | 48740901 |
| chr18 | 3632800 | 3632949 | ENSG00000266401 | 0.107568527 | 0.391771083 |  |  | TAAAGCCCATAACTTCCCCA | 3632948 | 3633148 |
| chr8 | 133596053 | 133596182 | ENSG00000261220 | -0.427016955 | 0.393951973 |  |  | GTACTACCTGATGTGCAGGA | 133595904 | 133596104 |
| chr5 | 58872901 | 58873220 | ENSG00000152932 | 0.056573482 | 0.406340934 |  |  | CATTGACCCTTACTGTTCAA | 58872898 | 58873098 |
| chr3 | 75585049 | 75587048 | ENSG00000272710 | 0.148841707 | 0.408538957 |  |  | CATGTTTATACTGTACACAA | 75585784 | 75585984 |
| chr9 | 5402973 | 5403052 | ENSG00000107020 | 0.163367021 | 0.409134247 |  |  | CAATTTATTTCACCAATACG | 5403047 | 5403247 |
| chr9 | 92116205 | 92116444 | ENSG00000275756 | 0.116715855 | 0.4091644 |  |  | TGGCAGAATTATTTGTAACA | 92116300 | 92116500 |
| chr1 | 119894503 | 119894552 | ENSG00000134250 | -0.101041335 | 0.438858365 |  |  | CCTTGGTCAGGTATTCCCAG | 119894425 | 119894625 |
| chr17 | 28404106 | 28404205 | ENSG00000004139 | -0.070742446 | 0.529379867 |  | X | TTTAAGAACATATTAATCGG | 28404183 | 28404383 |
| chr7 | 100337602 | 100337851 | ENSG00000214300 | 0.079248513 | 0.534001349 |  |  | CCTACCCCACCAAAATCCCA | 100337475 | 100337675 |
| chr1 | 89013053 | 89013142 | ENSG00000137944 | -1.084471206 | 0.55306902 |  |  | CCTTTAGTTCTAACAATGAA | 89013039 | 89013239 |
| chr5 | 54204945 | 54206704 | ENSG00000185305 | 0.334802559 | 0.558465875 |  |  | ACATAAAACAAACCAAACAG | 54205969 | 54206169 |
| chr15 | 48650851 | 48650950 | ENSG00000259705 | -0.050045296 | 0.569712888 |  |  | ATTCATAAAATCAAAAAGAG | 48650897 | 48651097 |
| chr5 | 80110701 | 80111360 | ENSG00000251675 | 0.057448286 | 0.58134695 |  |  | CTGCCTGTTGCCAAAAGCAG | 80110714 | 80110914 |
| chr5 | 168344651 | 168344750 | ENSG00000113645 | 0.058299594 | 0.59243178 |  |  | CGGGTTCATATTCCAACGGG | 168344628 | 168344828 |
| chr3 | 186615655 | 186616594 | ENSG00000283149 | -0.059454239 | 0.612820317 |  |  | CCTTCCCCACTTCCTCGACA | 186616458 | 186616658 |
| chr7 | 64682453 | 64683002 | ENSG00000196247 | -0.064595919 | 0.614693193 | X |  | CAGCCTCGATAACAGAGGCG | 64682966 | 64683166 |
| chr14 | 22562545 | 22563054 | ENSG00000129562 | -0.113703126 | 0.624236905 |  | X | TAAGACCCTATTCTTAAAGG | 22562628 | 22562828 |
| chr3 | 136056184 | 136057903 | ENSG00000227267 | 0.076150156 | 0.630676323 |  |  | TGAAAAGAACATCTACAGAG | 136056679 | 136056879 |
| chr3 | 120004050 | 120004099 | ENSG00000239835 | -0.079857132 | 0.650189715 |  |  | CTGCCTGTTGCCAAGAGCAG | 120003957 | 120004157 |
| chr1 | 105967694 | 105968103 | ENSG00000237480 | -0.042757719 | 0.65098277 |  |  | CTGCCTGTTGCCAAAAGCAG | 105967567 | 105967767 |
| chr6 | 143494051 | 143494100 | ENSG00000001036 | -0.130859432 | 0.656522987 |  |  | CCAGGGATACAAATGCCCAG | 143493996 | 143494196 |
| chr7 | 53810509 | 53810748 | ENSG00000205628 | -0.135075699 | 0.659476595 |  |  | ATTTATAAAATCAAAAAGAG | 53810447 | 53810647 |
| chr17 | 69512649 | 69513048 | ENSG00000267653 | -0.06683863 | 0.68457981 |  |  | GCAGACCTTTGGTCTTCAAG | 69512872 | 69513072 |
| chr9 | 92116298 | 92116357 | ENSG00000225511 | 0.050525643 | 0.694387813 |  |  | TGGCAGAATTATTTGTAACA | 92116300 | 92116500 |
| chr17 | 42092854 | 42093093 | ENSG00000108771 | 0.139988791 | 0.698998094 |  |  | TGGCAGAATTATTTGTAACA | 42093081 | 42093281 |
| chrX | 56546449 | 56546548 | ENSG00000188021 | 0.056264059 | 0.732447423 |  |  | TATATTGATACCATATAGAG | 56546443 | 56546643 |
| chr2 | 14517151 | 14517250 | ENSG00000237261 | -0.073140366 | 0.754529835 |  |  | ATTTATAAAATCAAAAAGAG | 14517042 | 14517242 |
| chr5 | 134925755 | 134925854 | ENSG00000279799 | -0.388412602 | 0.771901659 |  |  | TGGGGCCCTCTATTGCTGAG | 134925638 | 134925838 |
| chr3 | 186616546 | 186616855 | ENSG00000275696 | 0.031154766 | 0.793696223 |  |  | CCTTCCCCACTTCCTCGACA | 186616458 | 186616658 |
| chr17 | 28404146 | 28404255 | ENSG00000076351 | -0.207575338 | 0.809089792 |  | X | TTTAAGAACATATTAATCGG | 28404183 | 28404383 |
| chr16 | 89746000 | 89747399 | ENSG00000158805 | -0.040492179 | 0.846043817 |  |  | CCACAGTCATGCTCTCAGGA | 89746101 | 89746301 |
| chr8 | 61642000 | 61642099 | ENSG00000222898 | -0.018021703 | 0.861076186 |  |  | GGAAAATAAAATCTACGGCA | 61641927 | 61642127 |
| chr5 | 134925653 | 134925752 | ENSG00000113621 | 0.040834188 | 0.911680302 |  |  | TGGGGCCCTCTATTGCTGAG | 134925638 | 134925838 |
| chr8 | 120445500 | 120445649 | ENSG00000172167 | -0.035488991 | 0.91619706 |  |  | CCTGCTCCTGGTGATATGGG | 120445381 | 120445581 |
| chr7 | 100337297 | 100337776 | ENSG00000201913 | -0.013467722 | 0.936116363 |  |  | CCTACCCCACCAAAATCCCA | 100337475 | 100337675 |
| chr13 | 22795106 | 22795195 | ENSG00000237952 | -0.008976307 | 0.940049237 |  |  | CGTGACAGCAGCATACTGAA | 22795183 | 22795383 |
| chr10 | 95880848 | 95881147 | ENSG00000270099 | -0.012167852 | 0.946159672 |  |  | CTTCCTTTACCCAATTCAGA | 95880698 | 95880898 |
| chr10 | 50601356 | 50601745 | ENSG00000198964 | 0.04615091 | 0.946432153 | X | X | GTTCCAAGCATGCTGCAAAG | 50601626 | 50601826 |
| chr11 | 90142515 | 90143154 | ENSG00000077616 | -0.003950993 | 0.947656188 |  |  | ACATAAAACAAACCAAACAG | 90142646 | 90142846 |
| chr14 | 22562706 | 22563205 | ENSG00000277734 | 0.007592207 | 0.956742813 |  | X | TAAGACCCTATTCTTAAAGG | 22562628 | 22562828 |

**Supplementary Table S6**: gRNA targets of a Doxorubicin resistance screen overlapping STITCHIT regions

|  | **Blueprint** | **ENCODE** | **Roadmap** |
| --- | --- | --- | --- |
| **STITCHIT** | 162,453,061 | 33,229,288 | 165,805,230 |
| **Unified Peaks** | 165,590,118 | 70,571,718 | 273,259,579 |
| **GeneHancer** | 137,722,391 | 109,432,986 | 155,548,667 |

**Supplementary Table S7**: Total genomic space covered by the suggested regulatory regions shown for each dataset.

| **STITCHIT** |  |  |  |  |  |
| --- | --- | --- | --- | --- | --- |
| **chr** | **start** | **end** | **Reem ID** | **coefficient** | **OLS p-value** |
| chr5 | 138442350 | 138442489 | S4 | 0.12347378 | 0.13772486 |
| chr5 | 138449000 | 138449499 | S7 | 0.06534721 | 0.52094523 |
| chr5 | 138450250 | 138450399 | S10 | -0.03976734 | 0.65534283 |
| chr5 | 138454490 | 138456489 | S9 | 0.061487 | 0.55857386 |
| chr5 | 138457200 | 138457409 | S6 | 0.07735484 | 0.40351668 |
| chr5 | 138465700 | 138465949 | S2 | 0.21150264 | 0.05986547 |
| chr5 | 138469650 | 138470349 | S10 | 0.45303257 | 0.00021482 |
| chr5 | 138475650 | 138476489 | S5 | 0.11132624 | 0.22930495 |
| chr5 | 138486600 | 138486749 | S3 | -0.15254059 | 0.15120576 |
| chr5 | 138486950 | 138487099 | S8 | -0.06192892 | 0.56018674 |
| **Unified Peaks** | | | | | |
| **chr** | **start** | **end** | **Reem ID** | **coefficient** | **OLS p-value** |
| chr5 | 138443499 | 138444336 | U11 | -0.02633325 | 0.84467668 |
| chr5 | 138446871 | 138447822 | U10 | 0.04075988 | 0.76654673 |
| chr5 | 138454281 | 138454728 | U4 | 0.19843172 | 0.11272934 |
| chr5 | 138454842 | 138455474 | U7 | 0.07477763 | 0.58821766 |
| chr5 | 138462649 | 138471619 | U1 | 0.4100908 | 0.00147926 |
| chr5 | 138471717 | 138473719 | U9 | -0.04813992 | 0.77152777 |
| chr5 | 138473888 | 138474432 | U5 | 0.11320651 | 0.45458859 |
| chr5 | 138475851 | 138476436 | U3 | 0.21427014 | 0.12540462 |
| chr5 | 138476499 | 138477251 | U6 | 0.10563739 | 0.4268543 |
| chr5 | 138483081 | 138483386 | U8 | 0.00735699 | 0.95430046 |
| chr5 | 138486580 | 138487561 | U2 | -0.25307334 | 0.09496258 |

**Supplementary Table S8**: Candidate regulatory sites for EGR1.

| **#motif_id** | **sequence_name** | **start** | **stop** | **strand** | **score** | **p-value** | **q-value** | **matched_sequence** |
| --- | --- | --- | --- | --- | --- | --- | --- | --- |
| BHLHE40 | chr5:138469650-138470349 | 595 | 604 | - | 15.3654 | 1.97E-06 | 0.00254 | CTCACGTGCC |
| SP8 | chr5:138469650-138470349 | 565 | 576 | + | 15.7237 | 2.18E-06 | 0.0028 | ACCACTCCCACT |
| BCL6 | chr5:138469650-138470349 | 379 | 392 | - | 15.7097 | 2.22E-06 | 0.00302 | CTTCCTGGAAGGAT |
| TEAD1 | chr5:138469650-138470349 | 656 | 665 | - | 13.4355 | 2.92E-06 | 0.00402 | CACATTCCTC |
| TEAD4 | chr5:138469650-138470349 | 656 | 665 | - | 12.9516 | 2.92E-06 | 0.00403 | CACATTCCTC |
| STAT4 | chr5:138469650-138470349 | 382 | 395 | + | 15 | 4.65E-06 | 0.00636 | CTTCCAGGAAGAAA |
| STAT3 | chr5:138469650-138470349 | 382 | 392 | - | 14.898 | 5.56E-06 | 0.00764 | CTTCCTGGAAG |
| BHLHE41 | chr5:138469650-138470349 | 595 | 604 | + | 14.3846 | 1.20E-05 | 0.00799 | GGCACGTGAG |
| BHLHE41 | chr5:138469650-138470349 | 595 | 604 | - | 13.9038 | 1.35E-05 | 0.00799 | CTCACGTGCC |
| STAT3 | chr5:138469650-138470349 | 382 | 392 | + | 14.0816 | 1.25E-05 | 0.00857 | CTTCCAGGAAG |
| STAT1 | chr5:138469650-138470349 | 382 | 392 | + | 15.2727 | 6.68E-06 | 0.00915 | CTTCCAGGAAG |
| E2F6 | chr5:138469650-138470349 | 294 | 304 | - | 14.7759 | 7.24E-06 | 0.00918 | GGGAGGGAGAG |
| SP3 | chr5:138469650-138470349 | 565 | 575 | + | 13.7561 | 8.16E-06 | 0.0103 | ACCACTCCCAC |
| SOX3 | chr5:138469650-138470349 | 441 | 450 | - | 14.6724 | 8.14E-06 | 0.011 | CCTTTGTCCC |
| HES2 | chr5:138469650-138470349 | 591 | 603 | + | 12.6984 | 9.83E-06 | 0.0131 | TGTGGGCACGTGA |
| KLF16 | chr5:138469650-138470349 | 565 | 575 | + | 13.7241 | 1.16E-05 | 0.0136 | ACCACTCCCAC |
| ID2 | chr5:138469650-138470349 | 596 | 603 | + | 14.3443 | 1.21E-05 | 0.0148 | GCACGTGA |
| TFE3 | chr5:138469650-138470349 | 595 | 604 | - | 12.7347 | 1.48E-05 | 0.0174 | CTCACGTGCC |
| NR5A2 | chr5:138469650-138470349 | 622 | 636 | - | 13.3448 | 1.40E-05 | 0.0174 | CAGGTCAAGGCCTCT |
| STAT1 | chr5:138469650-138470349 | 382 | 392 | - | 12.4909 | 2.57E-05 | 0.0176 | CTTCCTGGAAG |
| BACH1::MAFK | chr5:138469650-138470349 | 281 | 295 | + | 9.75 | 2.87E-05 | 0.018 | GGGCCGACTCAGCCT |
| BACH1::MAFK | chr5:138469650-138470349 | 281 | 295 | - | 10.2105 | 2.42E-05 | 0.018 | AGGCTGAGTCGGCCC |
| SOX6 | chr5:138469650-138470349 | 441 | 450 | - | 13.8714 | 1.34E-05 | 0.0182 | CCTTTGTCCC |
| KLF12 | chr5:138469650-138470349 | 564 | 578 | + | 12.1475 | 1.42E-05 | 0.0182 | GACCACTCCCACTGC |
| TFAP2B(VAR.3) | chr5:138469650-138470349 | 253 | 265 | - | 12.3934 | 2.63E-05 | 0.0187 | TCCCCTAAAGGCC |
| TFAP2B(VAR.3) | chr5:138469650-138470349 | 253 | 265 | + | 11.6885 | 3.56E-05 | 0.0187 | GGCCTTTAGGGGA |
| TFAP2C | chr5:138469650-138470349 | 587 | 598 | - | 11.5102 | 5.45E-05 | 0.0197 | TGCCCACAGGGA |
| TFAP2C | chr5:138469650-138470349 | 554 | 565 | - | 12.0612 | 3.64E-05 | 0.0197 | TCCCTCAGGGGA |
| TFAP2C | chr5:138469650-138470349 | 554 | 565 | + | 11.8061 | 4.42E-05 | 0.0197 | TCCCCTGAGGGA |
| TFAP2C | chr5:138469650-138470349 | 117 | 128 | + | 11.0204 | 7.52E-05 | 0.0204 | tggccaaaggca |
| NFKB2 | chr5:138469650-138470349 | 579 | 591 | + | 11.2576 | 2.33E-05 | 0.0204 | TGGGGCAATCCCT |
| NFKB2 | chr5:138469650-138470349 | 579 | 591 | - | 10.3636 | 3.25E-05 | 0.0204 | AGGGATTGCCCCA |
| ARNTL | chr5:138469650-138470349 | 596 | 605 | - | 13.1639 | 1.83E-05 | 0.0208 | GCTCACGTGC |
| ARNTL | chr5:138469650-138470349 | 594 | 603 | + | 12.541 | 3.11E-05 | 0.0208 | GGGCACGTGA |
| ZNF263 | chr5:138469650-138470349 | 292 | 312 | - | 12.2857 | 1.64E-05 | 0.0211 | TGGGGAATGGGAGGGAGAGGC |
| RREB1 | chr5:138469650-138470349 | 662 | 681 | - | 8.16514 | 1.64E-05 | 0.0212 | TCCCCAACCAACAATCCACA |
| KLF13 | chr5:138469650-138470349 | 563 | 580 | + | 9.2931 | 1.60E-05 | 0.0213 | GGACCACTCCCACTGCTG |
| TFAP2C | chr5:138469650-138470349 | 587 | 598 | + | 10.5714 | 0.000101 | 0.022 | TCCCTGTGGGCA |
| SP2 | chr5:138469650-138470349 | 296 | 310 | + | 12.9828 | 1.78E-05 | 0.022 | CTCCCTCCCATTCCC |
| KLF1 | chr5:138469650-138470349 | 564 | 574 | + | 13.3636 | 1.70E-05 | 0.022 | GACCACTCCCA |
| TFAP2C(VAR.3) | chr5:138469650-138470349 | 253 | 265 | + | 11.1897 | 4.16E-05 | 0.0224 | GGCCTTTAGGGGA |
| TFAP2C(VAR.3) | chr5:138469650-138470349 | 253 | 265 | - | 11.0517 | 4.41E-05 | 0.0224 | TCCCCTAAAGGCC |
| BHLHE40 | chr5:138469650-138470349 | 595 | 604 | + | 13.1538 | 3.53E-05 | 0.0227 | GGCACGTGAG |
| KLF14 | chr5:138469650-138470349 | 564 | 577 | + | 11.7931 | 1.99E-05 | 0.0229 | GACCACTCCCACTG |
| TFAP2A(VAR.3) | chr5:138469650-138470349 | 487 | 499 | - | 9.65574 | 8.86E-05 | 0.0237 | GGCCCGGATGGGA |
| TFAP2A(VAR.3) | chr5:138469650-138470349 | 487 | 499 | + | 9.54098 | 9.20E-05 | 0.0237 | TCCCATCCGGGCC |
| TFAP2A(VAR.3) | chr5:138469650-138470349 | 253 | 265 | + | 11.2459 | 4.61E-05 | 0.0237 | GGCCTTTAGGGGA |
| TFAP2A(VAR.3) | chr5:138469650-138470349 | 253 | 265 | - | 10.5082 | 6.20E-05 | 0.0237 | TCCCCTAAAGGCC |
| FOXH1 | chr5:138469650-138470349 | 662 | 672 | - | 13.6923 | 1.81E-05 | 0.0249 | AACAATCCACA |
| TFAP2B | chr5:138469650-138470349 | 587 | 598 | - | 10.8909 | 8.38E-05 | 0.0251 | TGCCCACAGGGA |
| TFAP2B | chr5:138469650-138470349 | 587 | 598 | + | 10.7636 | 9.03E-05 | 0.0251 | TCCCTGTGGGCA |
| TFAP2B | chr5:138469650-138470349 | 554 | 565 | + | 11.7455 | 4.81E-05 | 0.0251 | TCCCCTGAGGGA |
| TFAP2B | chr5:138469650-138470349 | 554 | 565 | - | 11.7273 | 4.87E-05 | 0.0251 | TCCCTCAGGGGA |
| TFAP2B | chr5:138469650-138470349 | 117 | 128 | + | 10.3818 | 0.000114 | 0.0254 | tggccaaaggca |
| STAT5A::STAT5B | chr5:138469650-138470349 | 383 | 393 | - | 12.3878 | 3.71E-05 | 0.0255 | TCTTCCTGGAA |
| STAT5A::STAT5B | chr5:138469650-138470349 | 381 | 391 | + | 13.1429 | 2.45E-05 | 0.0255 | CCTTCCAGGAA |
| TFEC | chr5:138469650-138470349 | 595 | 604 | + | 12.8171 | 1.99E-05 | 0.0262 | GGCACGTGAG |
| NR5A2 | chr5:138469650-138470349 | 351 | 365 | + | 11.2241 | 4.22E-05 | 0.0263 | GAGGTAAAGGCCAGT |
| SP2 | chr5:138469650-138470349 | 168 | 182 | + | 11.2759 | 4.49E-05 | 0.0276 | CCCCCACCTCTACTC |
| TCF7L2 | chr5:138469650-138470349 | 330 | 343 | + | 13.1273 | 2.04E-05 | 0.0278 | GGGCATCAAAGAAG |
| TFAP2B | chr5:138469650-138470349 | 117 | 128 | - | 9.89091 | 0.000151 | 0.028 | TGCCTTTGGCCA |
| SP4 | chr5:138469650-138470349 | 562 | 578 | + | 7.56364 | 2.32E-05 | 0.0286 | GGGACCACTCCCACTGC |
| REL | chr5:138469650-138470349 | 189 | 198 | - | 12.8283 | 2.11E-05 | 0.0291 | TGGGTTTTCC |
| KLF13 | chr5:138469650-138470349 | 22 | 39 | - | 6.01724 | 4.53E-05 | 0.0301 | GGAACAGGCCCATTATCA |
| TFEB | chr5:138469650-138470349 | 595 | 604 | - | 12.7692 | 2.25E-05 | 0.0302 | CTCACGTGCC |
| EGR1 | chr5:138469650-138470349 | 565 | 578 | + | 9.51923 | 9.51E-05 | 0.031 | ACCACTCCCACTGC |
| EGR1 | chr5:138469650-138470349 | 453 | 466 | - | 9.90385 | 8.01E-05 | 0.031 | CTTCTGCCCACTCT |
| EGR1 | chr5:138469650-138470349 | 296 | 309 | + | 10.9423 | 4.92E-05 | 0.031 | CTCCCTCCCATTCC |
| EGR1 | chr5:138469650-138470349 | 290 | 303 | + | 10.6731 | 5.61E-05 | 0.031 | CAGCCTCTCCCTCC |
| PPARG::RXRA | chr5:138469650-138470349 | 439 | 453 | + | 12.3153 | 2.48E-05 | 0.0313 | GTGGGACAAAGGATA |
| EGR1 | chr5:138469650-138470349 | 168 | 181 | + | 8.84615 | 0.000127 | 0.0329 | CCCCCACCTCTACT |
| TFAP2C | chr5:138469650-138470349 | 117 | 128 | - | 9.59184 | 0.000182 | 0.033 | TGCCTTTGGCCA |
| HEY1 | chr5:138469650-138470349 | 595 | 604 | - | 11.7455 | 5.63E-05 | 0.0341 | CTCACGTGCC |
| HEY1 | chr5:138469650-138470349 | 595 | 604 | + | 11.4727 | 7.01E-05 | 0.0341 | GGCACGTGAG |
| TFAP2A(VAR.2) | chr5:138469650-138470349 | 587 | 598 | - | 10.4 | 0.000115 | 0.0346 | TGCCCACAGGGA |
| TFAP2A(VAR.2) | chr5:138469650-138470349 | 587 | 598 | + | 10.1636 | 0.000132 | 0.0346 | TCCCTGTGGGCA |
| TFAP2A(VAR.2) | chr5:138469650-138470349 | 554 | 565 | + | 11.6182 | 5.38E-05 | 0.0346 | TCCCCTGAGGGA |
| TFAP2A(VAR.2) | chr5:138469650-138470349 | 554 | 565 | - | 10.8727 | 8.68E-05 | 0.0346 | TCCCTCAGGGGA |
| ZNF263 | chr5:138469650-138470349 | 300 | 320 | - | 8.61224 | 0.000111 | 0.0359 | TGGGGACCTGGGGAATGGGAG |
| ZNF263 | chr5:138469650-138470349 | 291 | 311 | - | 9.63265 | 6.77E-05 | 0.0359 | GGGGAATGGGAGGGAGAGGCT |
| ZNF263 | chr5:138469650-138470349 | 229 | 249 | + | 8.69388 | 0.000107 | 0.0359 | AGAGGAGGTAAGAAGTAGGCA |
| KLF5 | chr5:138469650-138470349 | 565 | 574 | + | 10.8776 | 8.24E-05 | 0.0381 | ACCACTCCCA |
| KLF5 | chr5:138469650-138470349 | 296 | 305 | + | 10.3265 | 9.83E-05 | 0.0381 | CTCCCTCCCA |
| KLF5 | chr5:138469650-138470349 | 292 | 301 | + | 11.0204 | 7.50E-05 | 0.0381 | GCCTCTCCCT |
| KLF5 | chr5:138469650-138470349 | 168 | 177 | + | 9.69388 | 0.00012 | 0.0381 | CCCCCACCTC |
| STAT4 | chr5:138469650-138470349 | 379 | 392 | - | 11.6552 | 5.63E-05 | 0.0385 | CTTCCTGGAAGGAT |
| USF2 | chr5:138469650-138470349 | 595 | 605 | + | 11.6939 | 5.47E-05 | 0.0386 | GGCACGTGAGC |
| USF2 | chr5:138469650-138470349 | 594 | 604 | - | 11.5306 | 6.01E-05 | 0.0386 | CTCACGTGCCC |
| SP1 | chr5:138469650-138470349 | 168 | 178 | + | 12.5 | 3.08E-05 | 0.0388 | CCCCCACCTCT |
| TFAP2C(VAR.3) | chr5:138469650-138470349 | 487 | 499 | + | 7.15517 | 0.000151 | 0.0394 | TCCCATCCGGGCC |
| TFAP2C(VAR.3) | chr5:138469650-138470349 | 487 | 499 | - | 7.05172 | 0.000155 | 0.0394 | GGCCCGGATGGGA |
| TFE3 | chr5:138469650-138470349 | 595 | 604 | + | 11.0612 | 7.33E-05 | 0.0431 | GGCACGTGAG |
| TFAP2A(VAR.2) | chr5:138469650-138470349 | 117 | 128 | - | 9.29091 | 0.000213 | 0.0443 | TGCCTTTGGCCA |
| TFAP2A(VAR.2) | chr5:138469650-138470349 | 117 | 128 | + | 8.96364 | 0.000252 | 0.0443 | tggccaaaggca |
| HEY2 | chr5:138469650-138470349 | 595 | 604 | + | 10.7759 | 7.56E-05 | 0.0459 | GGCACGTGAG |
| HEY2 | chr5:138469650-138470349 | 595 | 604 | - | 10.7241 | 7.68E-05 | 0.0459 | CTCACGTGCC |
| ZFX | chr5:138469650-138470349 | 623 | 636 | - | 10.5 | 7.65E-05 | 0.0461 | CAGGTCAAGGCCTC |
| ZFX | chr5:138469650-138470349 | 521 | 534 | - | 10.4545 | 7.78E-05 | 0.0461 | GAACCTCAGGCCTC |
| TCFL5 | chr5:138469650-138470349 | 595 | 604 | + | 9.8871 | 8.31E-05 | 0.0467 | GGCACGTGAG |
| TCFL5 | chr5:138469650-138470349 | 595 | 604 | - | 9.8871 | 8.31E-05 | 0.0467 | CTCACGTGCC |
| TFAP2B(VAR.3) | chr5:138469650-138470349 | 487 | 499 | + | 6.91803 | 0.000164 | 0.0475 | TCCCATCCGGGCC |
| TFAP2B(VAR.3) | chr5:138469650-138470349 | 487 | 499 | - | 6.44262 | 0.000181 | 0.0475 | GGCCCGGATGGGA |
| NR5A2 | chr5:138469650-138470349 | 245 | 259 | + | 8.94828 | 0.000115 | 0.0479 | AGGCACAAGGCCTTT |
| SP1 | chr5:138469650-138470349 | 296 | 306 | + | 10.6346 | 7.65E-05 | 0.0482 | CTCCCTCCCAT |
| HIC2 | chr5:138469650-138470349 | 591 | 599 | - | 11.4182 | 3.72E-05 | 0.0483 | GTGCCCACA |

**Supplementary Table S9**: Results for FIMO for predicting TF binding in STITCHIT EGR1 enhancer E1 (q-value <=0.05)

| **motif_alt_id** | **sequence_name** | **start** | **stop** | **strand** | **score** | **p-value** | **q-value** | **matched_sequence** |
| --- | --- | --- | --- | --- | --- | --- | --- | --- |
| ZNF263 | chr5:138486600-138486749 | 100 | 120 | - | 16.4694 | 1.18E-06 | 0.000244 | AGAGAAGGGAAAGAGAGGGGG |
| ZNF263 | chr5:138486600-138486749 | 108 | 128 | - | 14.7755 | 3.66E-06 | 0.000272 | AGGGGAAGAGAGAAGGGAAAG |
| ZNF263 | chr5:138486600-138486749 | 107 | 127 | - | 14.6531 | 3.96E-06 | 0.000272 | GGGGAAGAGAGAAGGGAAAGA |
| ZNF263 | chr5:138486600-138486749 | 114 | 134 | - | 13.0408 | 1.06E-05 | 0.000365 | CAAGGAAGGGGAAGAGAGAAG |
| ZNF263 | chr5:138486600-138486749 | 113 | 133 | - | 13.2041 | 9.64E-06 | 0.000365 | AAGGAAGGGGAAGAGAGAAGG |
| ZNF263 | chr5:138486600-138486749 | 105 | 125 | - | 13.4694 | 8.23E-06 | 0.000365 | GGAAGAGAGAAGGGAAAGAGA |
| ZNF263 | chr5:138486600-138486749 | 96 | 116 | - | 11.9388 | 2.00E-05 | 0.000446 | AAGGGAAAGAGAGGGGGTAGA |
| ZNF263 | chr5:138486600-138486749 | 92 | 112 | - | 11.9388 | 2.00E-05 | 0.000446 | GAAAGAGAGGGGGTAGAAAGG |
| ZNF263 | chr5:138486600-138486749 | 120 | 140 | - | 11.7959 | 2.16E-05 | 0.000446 | AAAGAACAAGGAAGGGGAAGA |
| ZNF263 | chr5:138486600-138486749 | 101 | 121 | - | 12.0816 | 1.84E-05 | 0.000446 | GAGAGAAGGGAAAGAGAGGGG |
| ZNF263 | chr5:138486600-138486749 | 110 | 130 | - | 10.4286 | 4.51E-05 | 0.000846 | GAAGGGGAAGAGAGAAGGGAA |
| ZNF263 | chr5:138486600-138486749 | 95 | 115 | - | 10.2449 | 4.96E-05 | 0.000853 | AGGGAAAGAGAGGGGGTAGAA |
| ZNF263 | chr5:138486600-138486749 | 111 | 131 | - | 9.61224 | 6.84E-05 | 0.00102 | GGAAGGGGAAGAGAGAAGGGA |
| ZNF263 | chr5:138486600-138486749 | 103 | 123 | - | 9.59184 | 6.91E-05 | 0.00102 | AAGAGAGAAGGGAAAGAGAGG |
| ZNF263 | chr5:138486600-138486749 | 90 | 110 | - | 9.06122 | 8.98E-05 | 0.00124 | AAGAGAGGGGGTAGAAAGGAA |
| ZNF263 | chr5:138486600-138486749 | 87 | 107 | - | 8.08163 | 1.43E-04 | 0.00185 | AGAGGGGGTAGAAAGGAAAGG |
| ZNF263 | chr5:138486600-138486749 | 104 | 124 | - | 7.91837 | 1.54E-04 | 0.00188 | GAAGAGAGAAGGGAAAGAGAG |
| ZNF263 | chr5:138486600-138486749 | 118 | 138 | - | 7.71429 | 1.70E-04 | 0.00195 | AGAACAAGGAAGGGGAAGAGA |
| ZNF263 | chr5:138486600-138486749 | 84 | 104 | - | 7.10204 | 2.24E-04 | 0.00237 | GGGGGTAGAAAGGAAAGGTTA |
| ZNF263 | chr5:138486600-138486749 | 121 | 141 | - | 7.04082 | 2.30E-04 | 0.00237 | TAAAGAACAAGGAAGGGGAAG |
| ZNF263 | chr5:138486600-138486749 | 94 | 114 | - | 6.63265 | 2.75E-04 | 0.0027 | GGGAAAGAGAGGGGGTAGAAA |
| ZNF263 | chr5:138486600-138486749 | 89 | 109 | - | 6.02041 | 3.57E-04 | 0.00335 | AGAGAGGGGGTAGAAAGGAAA |
| PRDM1 | chr5:138486600-138486749 | 105 | 119 | - | 12.6727 | 1.96E-05 | 0.00434 | GAGAAGGGAAAGAGA |
| ZNF263 | chr5:138486600-138486749 | 97 | 117 | - | 4.87755 | 5.71E-04 | 0.00512 | GAAGGGAAAGAGAGGGGGTAG |
| ZNF263 | chr5:138486600-138486749 | 83 | 103 | - | 4.7551 | 5.99E-04 | 0.00516 | GGGGTAGAAAGGAAAGGTTAG |
| ZNF263 | chr5:138486600-138486749 | 76 | 96 | - | 4.4898 | 6.65E-04 | 0.0055 | AAAGGAAAGGTTAGGAGGAAT |
| ZNF263 | chr5:138486600-138486749 | 123 | 143 | - | 4.18367 | 7.49E-04 | 0.00578 | AATAAAGAACAAGGAAGGGGA |
| ZNF263 | chr5:138486600-138486749 | 109 | 129 | - | 4.16327 | 7.55E-04 | 0.00578 | AAGGGGAAGAGAGAAGGGAAA |
| ZNF263 | chr5:138486600-138486749 | 124 | 144 | - | 3.97959 | 8.10E-04 | 0.00598 | TAATAAAGAACAAGGAAGGGG |
| ZNF263 | chr5:138486600-138486749 | 98 | 118 | - | 3.61224 | 9.31E-04 | 0.00601 | AGAAGGGAAAGAGAGGGGGTA |
| ZNF263 | chr5:138486600-138486749 | 86 | 106 | - | 3.65306 | 9.17E-04 | 0.00601 | GAGGGGGTAGAAAGGAAAGGT |
| ZNF263 | chr5:138486600-138486749 | 81 | 101 | - | 3.71429 | 8.96E-04 | 0.00601 | GGTAGAAAGGAAAGGTTAGGA |
| ZNF263 | chr5:138486600-138486749 | 102 | 122 | - | 3.61224 | 9.31E-04 | 0.00601 | AGAGAGAAGGGAAAGAGAGGG |
| ZNF263 | chr5:138486600-138486749 | 88 | 108 | - | 3.30612 | 1.04E-03 | 0.00653 | GAGAGGGGGTAGAAAGGAAAG |
| ZNF263 | chr5:138486600-138486749 | 93 | 113 | - | 2.97959 | 1.18E-03 | 0.00694 | GGAAAGAGAGGGGGTAGAAAG |
| ZNF263 | chr5:138486600-138486749 | 75 | 95 | - | 2.97959 | 1.18E-03 | 0.00694 | AAGGAAAGGTTAGGAGGAATT |
| ZIC1 | chr5:138486600-138486749 | 26 | 39 | - | 8.65957 | 3.09E-05 | 0.00812 | GACACCCTGTAGTT |
| ZNF263 | chr5:138486600-138486749 | 99 | 119 | - | 2.22449 | 0.00154 | 0.00883 | GAGAAGGGAAAGAGAGGGGGT |
| ZNF263 | chr5:138486600-138486749 | 91 | 111 | - | 1.97959 | 1.68E-03 | 0.009 | AAAGAGAGGGGGTAGAAAGGA |
| ZNF263 | chr5:138486600-138486749 | 79 | 99 | - | 1.93878 | 1.70E-03 | 0.009 | TAGAAAGGAAAGGTTAGGAGG |
| ZNF263 | chr5:138486600-138486749 | 112 | 132 | - | 2 | 1.66E-03 | 0.009 | AGGAAGGGGAAGAGAGAAGGG |
| ZNF263 | chr5:138486600-138486749 | 117 | 137 | - | 1.26531 | 2.14E-03 | 0.011 | GAACAAGGAAGGGGAAGAGAG |
| IRF1 | chr5:138486600-138486749 | 103 | 123 | + | 9.41935 | 5.63E-05 | 0.0118 | CCTCTCTTTCCCTTCTCTCTT |
| ZIC4 | chr5:138486600-138486749 | 25 | 39 | - | 11.0182 | 4.77E-05 | 0.0125 | GACACCCTGTAGTTT |
| IRF1 | chr5:138486600-138486749 | 116 | 136 | + | 6.98387 | 1.43E-04 | 0.0149 | TCTCTCTTCCCCTTCCTTGTT |
| PRDM1 | chr5:138486600-138486749 | 99 | 113 | - | 6.27273 | 4.05E-04 | 0.0158 | GGAAAGAGAGGGGGT |
| PRDM1 | chr5:138486600-138486749 | 85 | 99 | - | 8.54545 | 1.60E-04 | 0.0158 | TAGAAAGGAAAGGTT |
| PRDM1 | chr5:138486600-138486749 | 84 | 98 | - | 7.8 | 2.20E-04 | 0.0158 | AGAAAGGAAAGGTTA |
| PRDM1 | chr5:138486600-138486749 | 112 | 126 | - | 6.47273 | 3.75E-04 | 0.0158 | GGGAAGAGAGAAGGG |
| PRDM1 | chr5:138486600-138486749 | 110 | 124 | - | 5.74545 | 4.93E-04 | 0.0158 | GAAGAGAGAAGGGAA |
| PRDM1 | chr5:138486600-138486749 | 101 | 115 | - | 5.70909 | 5.00E-04 | 0.0158 | AGGGAAAGAGAGGGG |
| ZNF263 | chr5:138486600-138486749 | 106 | 126 | - | -0.142857 | 3.35E-03 | 0.0169 | GGGAAGAGAGAAGGGAAAGAG |
| NFAT5 | chr5:138486600-138486749 | 19 | 28 | - | 10.9516 | 7.18E-05 | 0.0177 | GTTTTCCAAG |
| IRF1 | chr5:138486600-138486749 | 110 | 130 | + | 5.20968 | 0.000264 | 0.0184 | TTCCCTTCTCTCTTCCCCTTC |
| TFAP2C | chr5:138486600-138486749 | 47 | 58 | + | 10.7041 | 9.34E-05 | 0.0196 | TGCCTCAGAGCA |
| TFAP2C | chr5:138486600-138486749 | 47 | 58 | - | 10.0204 | 1.43E-04 | 0.0196 | TGCTCTGAGGCA |
| ZIC3 | chr5:138486600-138486749 | 25 | 39 | - | 7.7 | 7.91E-05 | 0.0199 | GACACCCTGTAGTTT |
| NFATC2 | chr5:138486600-138486749 | 21 | 27 | - | 12.2022 | 7.93E-05 | 0.021 | TTTTCCA |
| ZNF263 | chr5:138486600-138486749 | 63 | 83 | - | -1.06122 | 4.42E-03 | 0.0218 | GGAGGAATTTTATAGAGTACA |
| ZNF263 | chr5:138486600-138486749 | 78 | 98 | - | -1.18367 | 4.59E-03 | 0.022 | AGAAAGGAAAGGTTAGGAGGA |
| SPZ1 | chr5:138486600-138486749 | 118 | 128 | - | 9.97273 | 0.000149 | 0.0222 | AGGGGAAGAGA |
| SPZ1 | chr5:138486600-138486749 | 105 | 115 | - | 9.7 | 1.71E-04 | 0.0222 | AGGGAAAGAGA |
| PRDM1 | chr5:138486600-138486749 | 123 | 137 | - | 4.2 | 8.47E-04 | 0.0234 | GAACAAGGAAGGGGA |
| SP2 | chr5:138486600-138486749 | 94 | 108 | + | 6.05172 | 3.42E-04 | 0.0275 | TTTCTACCCCCTCTC |
| SP2 | chr5:138486600-138486749 | 118 | 132 | + | 6.41379 | 3.06E-04 | 0.0275 | TCTCTTCCCCTTCCT |
| SP2 | chr5:138486600-138486749 | 100 | 114 | + | 8.13793 | 1.71E-04 | 0.0275 | CCCCCTCTCTTTCCC |
| TFAP2B | chr5:138486600-138486749 | 47 | 58 | + | 10.5455 | 1.03E-04 | 0.0285 | TGCCTCAGAGCA |
| TFAP2B | chr5:138486600-138486749 | 47 | 58 | - | 9.29091 | 2.10E-04 | 0.0289 | TGCTCTGAGGCA |
| ZNF263 | chr5:138486600-138486749 | 119 | 139 | - | -2.28571 | 6.27E-03 | 0.0294 | AAGAACAAGGAAGGGGAAGAG |
| PRDM1 | chr5:138486600-138486749 | 90 | 104 | - | 3.07273 | 1.22E-03 | 0.0298 | GGGGGTAGAAAGGAA |
| ZNF263 | chr5:138486600-138486749 | 77 | 97 | - | -2.4898 | 0.00663 | 0.0304 | GAAAGGAAAGGTTAGGAGGAA |
| SPZ1 | chr5:138486600-138486749 | 79 | 89 | - | 8.26364 | 0.000371 | 0.0322 | AGGTTAGGAGG |
| SP1 | chr5:138486600-138486749 | 94 | 104 | + | 4.32692 | 3.89E-04 | 0.0325 | TTTCTACCCCC |
| SP1 | chr5:138486600-138486749 | 76 | 86 | + | 3.34615 | 4.53E-04 | 0.0325 | ATTCCTCCTAA |
| SP1 | chr5:138486600-138486749 | 124 | 134 | + | 2.38462 | 0.00052 | 0.0325 | CCCCTTCCTTG |
| SP1 | chr5:138486600-138486749 | 118 | 128 | + | 6 | 0.000288 | 0.0325 | TCTCTTCCCCT |
| SPI1 | chr5:138486600-138486749 | 119 | 132 | - | 0.381818 | 1.27E-04 | 0.0327 | AGGAAGGGGAAGAG |
| CDX2 | chr5:138486600-138486749 | 137 | 147 | - | 10.0339 | 1.43E-04 | 0.0337 | ATATAATAAAG |
| IRF1 | chr5:138486600-138486749 | 83 | 103 | + | 2.29032 | 0.000653 | 0.0341 | CTAACCTTTCCTTTCTACCCC |
| ELF3 | chr5:138486600-138486749 | 124 | 136 | - | 6.37931 | 1.29E-04 | 0.0345 | AACAAGGAAGGGG |
| ZNF263 | chr5:138486600-138486749 | 85 | 105 | - | -3.06122 | 0.00772 | 0.0347 | AGGGGGTAGAAAGGAAAGGTT |
| TEAD4 | chr5:138486600-138486749 | 73 | 82 | + | 10.4677 | 1.44E-04 | 0.0347 | AAAATTCCTC |
| SPIC | chr5:138486600-138486749 | 119 | 132 | - | 9.50704 | 1.46E-04 | 0.0378 | AGGAAGGGGAAGAG |
| SP2 | chr5:138486600-138486749 | 76 | 90 | + | 2.58621 | 8.62E-04 | 0.0383 | ATTCCTCCTAACCTT |
| SP2 | chr5:138486600-138486749 | 124 | 138 | + | 2.51724 | 0.000876 | 0.0383 | CCCCTTCCTTGTTCT |
| SP2 | chr5:138486600-138486749 | 111 | 125 | + | 2.15517 | 9.51E-04 | 0.0383 | TCCCTTCTCTCTTCC |
| RARA | chr5:138486600-138486749 | 86 | 103 | - | -14.1515 | 1.62E-04 | 0.039 | GGGGTAGAAAGGAAAGGT |
| EGR1 | chr5:138486600-138486749 | 94 | 107 | + | 8.28846 | 1.59E-04 | 0.0392 | TTTCTACCCCCTCT |
| ELF5 | chr5:138486600-138486749 | 125 | 135 | - | 10.2909 | 1.51E-04 | 0.0393 | ACAAGGAAGGG |
| EGR1 | chr5:138486600-138486749 | 118 | 131 | + | 6.19231 | 0.000345 | 0.0425 | TCTCTTCCCCTTCC |
| FOSL2 | chr5:138486600-138486749 | 44 | 54 | + | 8.41818 | 0.000159 | 0.0427 | TAATGCCTCAG |
| PRDM1 | chr5:138486600-138486749 | 117 | 131 | - | 1.09091 | 2.16E-03 | 0.0433 | GGAAGGGGAAGAGAG |
| PRDM1 | chr5:138486600-138486749 | 116 | 130 | - | 1.21818 | 0.00208 | 0.0433 | GAAGGGGAAGAGAGA |
| TBP | chr5:138486600-138486749 | 69 | 83 | + | 9.76389 | 0.00019 | 0.0464 | CTATAAAATTCCTCC |
| IRF1 | chr5:138486600-138486749 | 104 | 124 | + | 0.193548 | 1.17E-03 | 0.0488 | CTCTCTTTCCCTTCTCTCTTC |
| GATA4 | chr5:138486600-138486749 | 113 | 123 | + | 9.17241 | 1.90E-04 | 0.0495 | CCTTCTCTCTT |

**Supplementary Table S10:** FIMO results predicting TF binding in STITCHIT EGR1 enhancer E3 (q-value <=0.05)
